# Structural modularity of receptor-binding proteins underlies host-range strategy diversification in *Klebsiella pneumoniae* phages

**DOI:** 10.64898/2026.05.12.724579

**Authors:** Vyshakh R. Panicker, Bogna J. Smug, Victor Klein-Sousa, Mark C. Enright, Nicholas M.I. Taylor, Zuzanna Drulis-Kawa, Rafał J. Mostowy

**Affiliations:** Doctoral School of Exact and Natural Sciences, Jagiellonian University, Kraków, Poland; Malopolska Centre of Biotechnology, Jagiellonian University, Kraków, Poland; Department of Drug Design and Pharmacology, Faculty of Health and Medical Sciences, University of Copenhagen, Universitetsparken 2, 2100 Copenhagen, Denmark; Norvo Nordisk Foundation Center for Protein Research, Department of Cellular and Molecular Medicine, Faculty of Health and Medical Sciences, University of Copenhagen, Blegdamsvej 3B, 2200 Copenhagen, Denmark; Department of Experimental Medical Science, Lund University, Lund, Sweden; Department of Life Sciences, Manchester Metropolitan University, Manchester, United Kingdom; Department of Pathogen Biology and Immunology, University of Wroclaw, Wrocław, Poland

## Abstract

Bacteriophage receptor-binding proteins (RBPs) determine bacterial host recognition and are central to phage host-range evolution, yet the structural principles governing how RBPs diversify and expand host range remain not fully understood. Although phage RBPs are thought to evolve through modular exchange of receptor-binding domains, it is unclear how this modularity is organised across diverse RBP architectures, at what structural scales recombination operates, and how it shapes host-range breadth. Here we combined large-scale AlphaFold3 structural modelling, domain-level annotation, de novo pseudo-domain segmentation, sequence modularity analysis and experimentally determined host-range phenotypes across 192 *Klebsiella pneumoniae* phages and 382 high-confidence RBPs spanning up to 96 K-types. We generated the first system-wide structural atlas of *K. pneumoniae* phage RBPs, resolving 39 structurally distinct RBP-classes. Although capsule-degrading beta-helix depolymerases dominated numerically, 161 RBPs across 37 RBP-classes employed non-depolymerase architectures, including 18 novel RBP-classes. Structural and sequence analyses show that this diversity arose through modular reuse of structural domains rather than independent invention. Conserved N-terminal scaffolds linked depolymerase and non-depolymerase RBPs across morphotypes, while receptor-binding regions diversified through recombination operating at and beyond domain boundaries. We found recent modular exchange both within genera, where capsule-specific and capsule-independent RBPs can be swapped to alter host-range strategy, and across morphotypes, where depolymerase modules moved between distant lineages altering capsule specificity. Together, these architectures resolve into six receptor-recognition strategies, establishing multi-scale modularity as the primary organising principle of RBP diversification and a structure-informed framework for guiding phage isolation and engineering against *K. pneumoniae*. The complete atlas is freely accessible as an interactive community resource at *Klebsiella*-Phage-RBP-Atlas.

## Introduction

Bacteriophages (phages) are the dominant predators of bacteria and a cornerstone of the emerging phage therapy field, yet a fundamental question remains unresolved: what genetic and phenotypic features determine how broadly a phage can infect^1–4^? Phage-bacteria coevolution is characterised by recurring cycles: as bacterial hosts diversify, broad host-range variants gain a selective advantage by infecting across multiple strains or serotypes, but specialists re-emerge as hosts counter-adapt to resist them^5^. These dynamics play out across ecological and evolutionary timescales and have been documented in multiple systems^6^. Broad host-range variants are therefore not evolutionary endpoints but transient advantages — and precisely because of this, they are disproportionately valuable clinically, where matching phages to the diverse surface architectures of drug-resistant isolates is a key limiting factor in phage therapy^7–11^.

Recognition of a bacterial surface receptor by a phage receptor-binding protein (RBP) is the first step in a phage infection and is also the primary determinant of host-range^12–14^. Phages deploy multiple classes of RBPs, typically classified as tail fibers and tailspikes, to engage diverse receptors including the capsular polysaccharide (CPS), lipopolysaccharide (LPS), O-antigen, outer-membrane proteins (OMPs), and appendages like pili and flagella^12,15–18^.

Tail fibers are elongated structures built around an interweaving beta-helical spine that terminates in a distal tip domain of varied geometry, where receptor-binding loops make contact with the surface target; some tail fibers additionally carry adhesin proteins at the distal tip that extend or fine-tune receptor specificity^19–23^. Tailspikes, by contrast, fold into compact globular trimers whose C-terminal domains are enzymatically active, deploying hydrolase or lyase activity against carbohydrate substrates including CPS and LPS^24–27^. Despite these structural differences, both classes share N-terminal anchoring scaffolds for attachment to the phage baseplate or tail apparatus^19^. This structural architecture allows phages to respond to persistent selection pressure to infect by exchanging structural domains, generating chimeric proteins with altered receptor specificity^14,28,29^. This modular architecture – building new binding properties by recombining existing structural elements – is a recurring feature of phage RBP evolution and the primary mechanism by which phages generate novel receptor-recognition configurations, with host-range diversification as one downstream consequence^13,30,31^.

*Klebsiella pneumoniae* (*K. pneumoniae*) represents an especially informative system for studying RBP evolution because its surface architecture is both clinically important and extraordinarily diverse^32^. The CPS is widely regarded as the dominant receptor for *Klebsiella* phages, and genomic surveys now identify over 150 distinct capsular locus types (KL-type), encompassing 77 serologically defined reference capsule types (K-types)^32–36^. Beyond variation in sugar composition and linkage, capsule thickness and density varies markedly across lineages and different K-types^37–40^. Work on isolated *Klebsiella* phages has established a predominantly capsule-centric paradigm of host recognition, in which phages deploy capsule-degrading depolymerases, most commonly adopting single-stranded beta-helix (SSBH) folds to access CPS and typically displaying narrow host-range constrained by the enzyme’s substrate specificity^25,27,41–43^. This paradigm, however, is largely silent on phages that engage non-capsular surface receptors through non-depolymerase RBPs, a category that standard isolation practices have systematically underrepresented^9,44^.

The capsule-centric paradigm does not account for the full diversity of *Klebsiella* phages^9,44^. Collections assembled using standard encapsulated reference strains as isolation hosts — conditions biased towards halo-forming, depolymerase-carrying phages, nevertheless include phages that lack detectable depolymerases yet display broad host-range^41,43,45^. That such phages were recovered despite conditions selective against them suggests their presence in *Klebsiella* phage communities is not incidental. This is reinforced by two recent studies who used capsule-loss mutant hosts of *K. pneumoniae* for phage isolation and recovered a substantially larger and more diverse repertoire of non-depolymerase RBPs with broad host-range activity, demonstrating that standard conditions actively suppress recovery of this diversity rather than merely failing to enrich it^9,44^. *Klebsiella* phages therefore navigate a receptor landscape that is varied, dynamic, and not fully captured by the capsule-centric model — and their RBPs are the primary tools by which they do so. The exchange and reshuffling of structural domains, leading to RBP modularity, is the primary mechanism by which phages continuously generate the receptor-recognition diversity required to keep pace with evolving bacterial hosts^5,29–31^.

What remains unresolved is how RBP modularity is structurally organised across a diverse RBP repertoire, at what scales it operates, and whether RBP domain architecture independently shapes host-range breadth or merely reflects shared evolutionary history. Addressing this requires going beyond cataloguing which RBPs exist, and instead asking three specific questions. First, does modularity operate only at classical domain boundaries, or does it extend to structural units below this level? Second, how is the full RBP structural repertoire organised across a diverse phage collection from a single host system, and what fraction of that repertoire lies outside the currently characterised repertoire? Third, does the domain composition of an RBP independently predict how broadly a phage can infect, or is structural architecture merely a reflection of the taxonomic background?

To address these questions, we assembled 192 virulent *Klebsiella* phages drawn predominantly from published collections with experimentally determined host-range phenotypes spanning up to 96 K-types, supplemented by newly isolated phages characterised by their depolymerase repertoire. From these genomes, we predicted 382 high-confidence RBP structures using AlphaFold3 structural modelling, annotated their domain architecture through Evolutionary Classification of protein Domains (ECOD) and pseudo-domain segmentation, and integrated the resulting structural atlas with both experimentally determined host-range phenotypes and a depolymerase-based genetic host-range classification as a complementary, dataset-wide measure^30,46–48^. This structural atlas provides a system-wide view of how RBP diversity is organised, how it maps onto phage host-range, and what it implies for the ecological conditions under which phages with different receptor-recognition strategies are most likely to be recovered.

## Results

### Phage Dataset Overview and Host-Range Distribution

A total of 192 virulent *Klebsiella* phage genomes were assembled and curated: 175 from publicly available host-range collections with experimentally confirmed specificities against 96 K-types, and 17 isolated in this study from wastewater in Manchester with confirmed specificities against 9 K-types. Halo and/or plaque formation was the primary inclusion criterion, with the full infection phenotype distribution shown in Figure 1A and Figure S1. Among all phages, 104 produced both halo and plaque, 76 produced plaques with no halo information reported, 7 explicitly produced plaques only, and one produced only a halo. Four phages lacking both phenotypes were retained because at least one depolymerase had been heterologously expressed with experimentally confirmed K-type specificity (Table S1). All 17 phages isolated in this study produced both halo and plaque.

**Figure 1:**
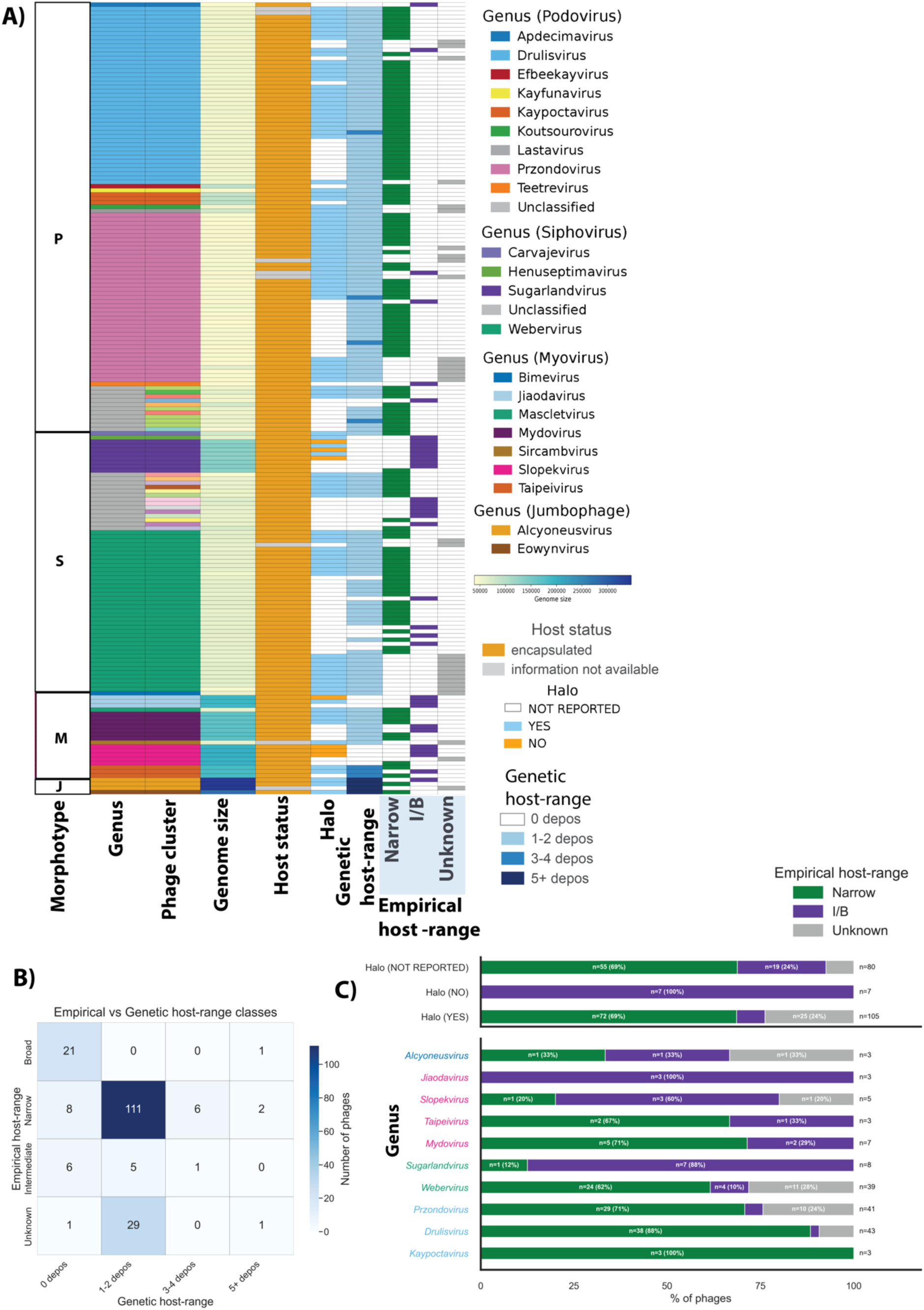
Taxonomic, morphotypical, and phenotypic landscape of 192 virulent *Klebsiella* phages and their host-range distributions. (A) Comprehensive genomic and phenotypic profiling of the curated phage dataset. Each row represents an individual phage genome (n=192). Columns detail (from left to right): morphotype of the phage ((P)odovirus, (S)iphovirus, (M)yovirus, or (J)umbophage), assigned taxonomic genus, wGRR-based phage cluster, Genome size (represented by a continuous color gradient), isolation host status (encapsulated/not reported), reported halo formation phenotype during infection, genetic host-range (calculated based on the number of SSBH-fold depolymerase a phage harbours) and empirical host-range (calculated based on the number of K-types a phage can infect independent of the genetic host-range). Empirical host-range was categorized as Narrow (infecting ≤2 K-types) or Intermediate/Broad (I/B; infecting ≥3 K-types), with phages tested against insufficient panel sizes (<5 K-types) designated as Unknown. (B) A 4×4 confusion matrix illustrating the concordance and discordance between empirical host-range categories (rows: Broad, Narrow, Intermediate, Unknown) and genetic host-range inferred from SSBH-fold depolymerase counts (columns: 0, 1–2, 3–4, 5+ depolymerases). Concordant cells are those where empirical and genetic host-range classifications are mutually consistent: 21 broad host-range phages encoding zero SSBH-fold depolymerases, 111 narrow host-range phages encoding 1–2 SSBH-fold depolymerases, and 1 intermediate host-range phage encoding 3–4 SSBH-fold depolymerases. All remaining off-diagonal cells represent discordance between the two schemes. (C - top) Association between observable infection phenotypes and empirical host-range. Halo formation is strongly associated with highly specific receptor targeting, with 69% of halo-forming phages exhibiting a narrow empirical host-range. 7 I/B phages did not produce a halo, and finally, 69% (n = 55) narrow empirical host-range phages had no halo reported in the respective studies. (C-bottom) Empirical host-range distribution across ten phage genera (≥3 phages each). Genera are grouped by morphotype: podovirus (blue), siphovirus (green), myovirus (pink), and jumbophages (dark blue). Only *Kaypoctavirus* and *Jiaodavirus* show homogeneous host-range profiles; others display mixed profiles, reflecting test panel biases and within-genus host-range diversity.

The dataset comprised 104 podoviruses, 63 siphoviruses, 25 myoviruses, of which 4 are jumbophages (henceforth, we refer to jumbophage myoviruses as “jumboviruses” and non-jumbophage myoviruses as “myoviruses”). Genome sizes varied substantially: podoviruses ∼44 Kbp, siphoviruses ∼57 Kbp, myoviruses ∼143 Kbp, and jumbophages ∼326 Kbp (Figure 1A, Table S1). Of the 192 genomes, 167 were assigned to 22 known genera (9 podovirus, 4 siphovirus, 7 myovirus, 2 jumbophage), while 25 remained unclassified. An additional 18 novel genera (hereafter phage clusters), were identified by weighted Gene Repertoire Relatedness (wGRR)-based genome clustering calibrated against known taxa (Table S1, Figures S2, S3). Three genera predominated: *Drulisvirus* (22%), *Przondovirus* (21%), and *Webervirus* (20%).

Because phages across the dataset were tested against host panels of different sizes, direct comparison of raw infection counts was not possible. To enable system-wide comparison of infection breadth, we therefore applied two complementary classification schemes. The first classification scheme was based on the experimental information included in the study (henceforth referred to as “empirical host-range”). This classification grouped phages by the number of K-types they were reported to infect: narrow (≤2), intermediate (≤4), or broad (≥5) (Figure S4). Because collections in the dataset were tested against host panels of different sizes, only phages challenged against at least 5 K-types were assigned a category; below this threshold a single infection event can shift a phage between categories, making the assignment unreliable (Figure 1A, Figure S1). Intermediate and broad categories are merged into a single Intermediate/Broad (I/B) category in Figure 1A. The second classification scheme was based on the inference of K-type specific depolymerases (henceforth referred to as “genetic host-range”). In this classification, we grouped phages independently by the number of SSBH-fold depolymerases detected in each genome: “zero”, “1-2”, “3-4”, or “5+”. The two schemes showed near-complete agreement (Figure 1B): 111 narrow empirical host-range phages encoded 1–2 depolymerases, 1 intermediate empirical host-range phage encoded 3–4 depolymerases, and 21 broad empirical host-range phages encoded none, all consistent with the capsule-centric model (phages with unknown empirical host-range, n=31, were excluded from this comparison). Minor discordances were apparent — 8 phages with multiple depolymerases displayed narrow empirical host-range, possibly reflecting insufficient panel coverage, and 14 phages encoding zero depolymerases displayed narrow or intermediate empirical host-range, possibly due to undetected depolymerases or targeting of alternative receptors such as O-antigen. Overall, the two schemes were in strong agreement, supporting SSBH-fold depolymerase content as a reliable proxy for empirical host-range.

Given the near-complete agreement between the empirical and genetic host-range schemes, we reasoned that halo formation — a visible proxy for enzymatic capsule degradation — should be a strong predictor of narrow host-range. This was largely confirmed: 69% (n=72) of halo-forming phages displayed narrow empirical host-range, concentrated within *Drulisvirus*, *Przondovirus* and *Webervirus*. However, the association was imperfect: 8 halo-forming phages exhibited I/B empirical host-range, and the 7 phages that produced no halo fell exclusively within the I/B empirical host-range category, concordant with the absence of SSBH-fold depolymerases (Figure 1C - top). Halo formation and depolymerase content are therefore correlated but not interchangeable proxies for host-range breadth.

Taxonomy and morphotype were both associated with host-range breadth, though not perfectly. As expected, narrow host-range phages were predominantly podoviruses and siphoviruses with smaller genomes, while I/B phages comprised mainly siphoviruses and myoviruses with larger genomes (Figure 1A). However, even at the genus level host-range was not uniform: 8 of 10 genera with at least three representatives contained phages spanning different empirical host-range categories, with only *Kaypoctavirus* and *Jiaodavirus* showing a homogeneous profile (Figure 1C-bottom). Because phages within a genus share the majority of their gene content, this within-genus variation points to a small number of rapidly evolving loci — most likely the RBP locus — rather than genome-wide divergence.

In summary, since almost all isolation hosts across the dataset were encapsulated K-type reference strains, the collection was expected to be dominated by narrow host-range, depolymerase-dependent phages — and narrow host-range was indeed the dominant outcome (n=127). The recovery of 34 I/B phages (intermediate n=12, broad n=22) despite these conditions, and the reproducible recurrence of the same depolymerase-independent genera across two independent collections^43,45^, confirms that this diversity is real and not a sampling artefact. What receptor-recognition strategies these phages employ, and what structural features of their RBPs underpin those strategies, remains unknown. The remainder of this study addresses this question through a systematic structural analysis of the full RBP repertoire, linking molecular architecture back to the host-range phenotypes described here.

### Atlas of Receptor Binding Protein Architectures

To catalogue the full structural repertoire of RBPs encoded by the 192 phages in our dataset, we first identified all RBP candidates computationally. 457 RBP candidates were identified across all 192 genomes using functional annotation with PHROGs combined with machine learning-based detection with PhageRBPdetect, filtered by protein length (>=200 aa) and prediction confidence score (>=0.95)^49,50^ (see Methods). Structural models were generated for all candidates using AlphaFold3, with high-quality structures retained when the ranking score exceeded 0.5 or with an average RBPseg merge score ≥80 for models that needed further refinement^30,46^. Of the 457 RBP candidates, 75 were removed after quality filtering or identified as false positives, yielding 382 high-confidence RBP structures carried forward for classification (Table S2, Figure S5).

These 382 structures were first grouped into 148 RBP clusters using Foldseek structural clustering (multimer TMscore threshold 0.65)^51^, which served as the basis for RBP-class assignment. An RBP-class groups one or more RBP clusters sharing a common overall architecture and, where determinable, an inferred functional identity. The 148 RBP clusters were assigned to 39 RBP-classes through a sequential classification pipeline that prioritised confident matches and fell back on progressively more distant comparisons where strong matches were unavailable (see Methods). A functional assignment was considered successful when structural similarity to a characterised RBP was supported by a strong TM-score match to a PDB entry^52^, placement by RBPseg-classify^30^, or direct literature precedent^53–55^; such RBP-classes are referred to as functionally assigned throughout. RBP-classes with no detectable similarity to any previously described RBP are referred to as structurally novel and represent candidate receptor-recognition strategies not previously associated with *Klebsiella* phage infection. Of the 39 RBP-classes, 21 (encompassing 334 RBPs) received functional assignments, while 18 RBP classes (48 RBPs) were structurally novel (see Methods; Text S1 for the full classification pipeline).

The 39 RBP-classes revealed a repertoire that extends well beyond the capsule-targeting depolymerases that dominate standard isolation collections. As expected, the majority of all RBPs were SSBH-fold depolymerases (n=221; 58%) distributed across two RBP-classes: tailspike depolymerases (n=186) and tailspike depolymerases with fiber-like elements (n=35) (Figure 2E). The remaining 161 RBPs distributed across 37 classes encompassed a broad range of architectures: tail fibers of diverse geometries, adhesins, enzymatically active tailspikes targeting non-capsular substrates, and structural classes with no known homologue. Of these 37 RBP-classes, 17 RBP-classes (40 RBPs) lacked detectable structural similarity to any characterised RBP, possibly representing architectures that engage bacterial surface components not yet associated with phage adsorption in *Klebsiella* — hence constitute candidates for structural and functional validation. One additional class, structurally similar to T4 gp37, was retained among the structurally novel due to the absence of literature precedent in *Klebsiella* phages and the substantially different gp37 structure described in the T4 literature; its co-occurrence with T4 gp34-like, gp36-like, and gp12-like proteins in the same phage genomes nonetheless supports an indirect functional inference as part of a T4-like tail fiber system. The distribution of these classes across morphotypes is described in the following paragraphs and summarised in Figure 2, Figure S6 and Text S1.

**Figure 2:**
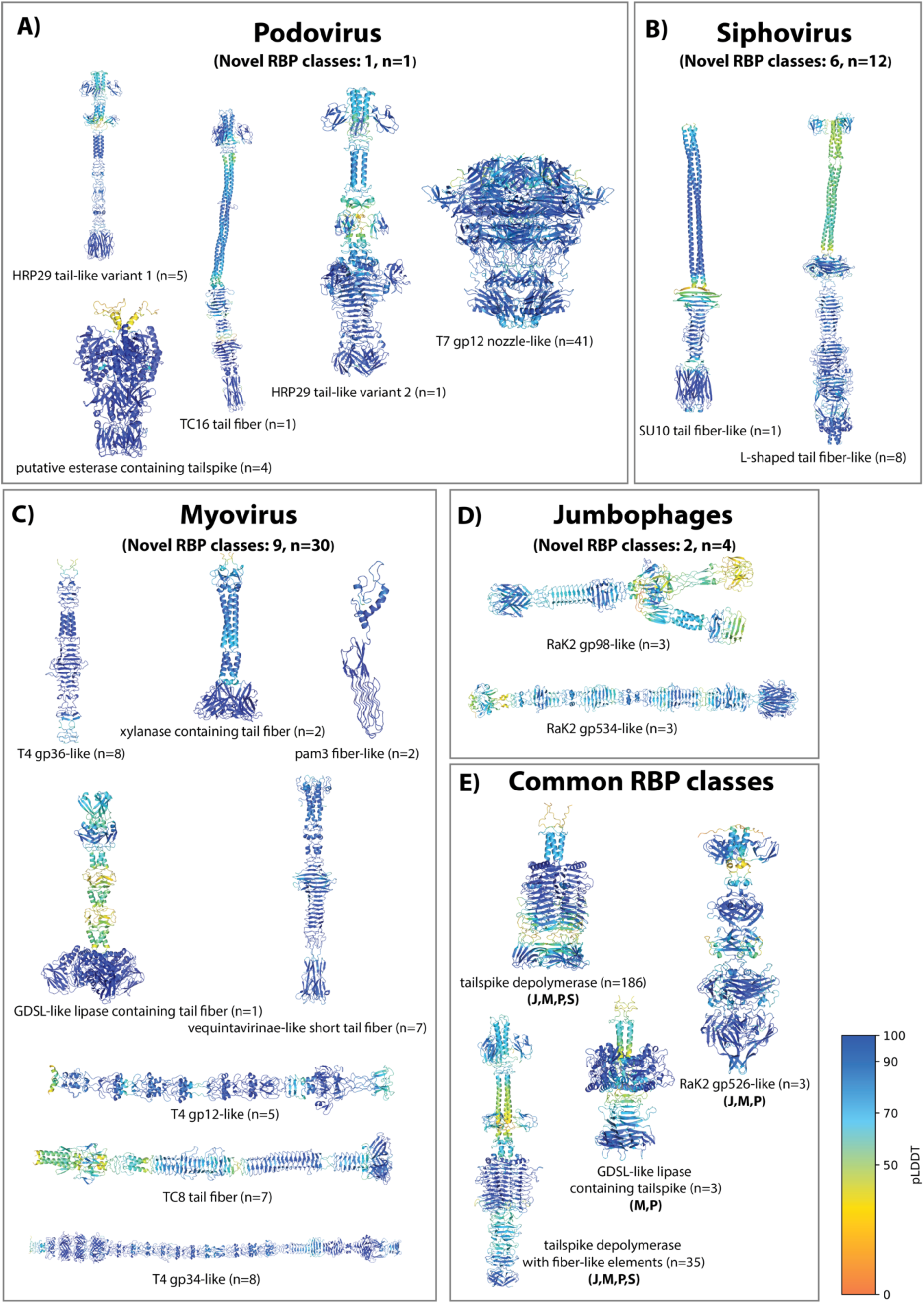
Structural models and morphotype-specific distributions of functionally assigned RBP-classes. High-confidence RBP structural models (n=334) were generated and classified based on homology to known RBP structures. Structures are coloured according to the AlphaFold per-residue confidence metric (pLDDT; 0–100 scale), with dark blue indicating high-confidence regions. Each panel displays only the functionally assigned RBP-classes exclusive to that morphotype, with the number of additional structurally novel RBP-classes and their combined RBP count indicated in the panel header. (A) Podovirus-exclusive functionally assigned RBP-classes (n=52), including T7 gp12 nozzle-like hexamers, HRP29 tail-like variants, TC16 tail fibers, and putative esterase-containing tailspikes. (B) Siphovirus-exclusive functionally assigned RBP-classes (n=9), featuring SU10 and L-shaped tail fiber-like RBP-classes. (C) Myovirus-exclusive functionally assigned RBP-classes (n=40), comprising T4-like tail fiber subunits (gp34-like, gp36-like), T4 gp12-like short tail fibers, TC8 tail fibers, vequintavirinae-like short tail fibers, xylanase-containing tail fibers, GDSL-like lipase-containing tail fibers, and pam3 fiber-like RBP-classes. (D) Jumbophage-exclusive functionally assigned classes (n=6), including RaK2 gp534-like and RaK2 gp98-like fibers. (E) RBP-classes shared across multiple morphotypes (n=227), dominated by SSBH-fold tailspike depolymerases (n=186) and tailspike depolymerases with fiber-like elements (n=35), both present across all four morphotypes (Jumbophage (J), Myovirus (M), Podovirus (P), Siphovirus (S)), alongside RaK2 gp526-like and GDSL-like lipase-containing tailspike classes with narrower cross-morphotype distributions. All protein structures are shown and described in the Klebsiella RBP Atlas Text S1.

RBP-classes were strongly partitioned by morphotype: 35 of 39 RBP-classes were exclusive to a single morphotype (Figure S6, Figures 2A–D). Across all morphotypes, tailspike depolymerases were the numerically dominant RBP-class, but their relative prevalence varied substantially, and each morphotype harboured a distinct complement of non-depolymerase architectures — including structurally novel RBP-classes.

Podoviruses were the most numerically dominant morphotype (n=188 RBPs, 10 RBP-classes) but their repertoire was heavily concentrated in two RBP-classes: tailspike depolymerase (n=132, 70%) and T7 gp12 nozzle-like RBPs (n=41, 22%)^30,56^, which accounted for 92% of all podovirus RBPs (Figure 2A). The remaining RBP-classes were rare, including HRP29-like variants and SGNH hydrolase-containing tailspikes in *Przondovirus*, suggesting possible non-capsular (e.g. O-antigen) targets^26^. Only one uncharacterised RBP-class was identified (Figure 2A, Figure S6, Text S1).

Siphoviruses contributed 60 RBPs across 10 RBP-classes, with tailspike depolymerases again dominant (n=38, 63%). The most informative non-depolymerase RBP class was the L-shaped tail fiber-like (n=8), structurally similar to T5 fibers that recognise oligo-mannose — consistent with targeting a conserved non-capsular surface receptor^21^ (Figure 2B). Six structurally novel RBP-classes covering 13 RBPs were also identified (Figure S6, Text S1).

Myoviruses were the most structurally diverse morphotype (21 RBP-classes, 93 RBPs), with tailspike depolymerases representing only a minority (n=21, 23%). Functionally assigned classes included T4-like tail fiber subunits (gp34-like, gp36-like), T4 gp12-like short tail fibers, TC8 tail fibers, vequintavirinae-like short tail fibers, gp38-like adhesins targeting OMP receptors, and xylanase- and GDSL-like lipase-containing tail fibers (Figure 2C)^31,55,57–60^. Nine uncharacterised RBP-classes covering 22 RBPs represented the largest reservoir of novel architectures in the dataset (Figure S6, Text S1).

Jumbophages contributed 41 RBPs across 7 RBP-classes, dominated by tailspike depolymerases (n=30, 73%). Two functionally assigned RBP-classes in *Alcyoneusvirus* — RaK2 gp534-like and RaK2 gp98-like — both feature jelly-roll receptor-binding domains (Figure 2D)^53^. Two uncharacterised RBP-classes (TC10-like in *Alcyoneusvirus*, TC16-like in *Eowynvirus*) covered 4 RBPs (Figure S6, Text S1).

Four RBP-classes spanned multiple morphotypes, together accounting for the majority of all RBPs (n=227, Figure 2E). Tailspike depolymerases and tailspike depolymerases with fiber-like elements occurred across all four morphotypes, with podoviruses contributing the most (n=118 and n=14) and myoviruses the fewest (n=15 and n=6). The RaK2 gp526-like RBP class spanned jumbophages, myoviruses and podoviruses. GDSL-like lipase-containing tailspikes were detected in myoviruses and podoviruses, invariably co-occurring with tailspike depolymerases^53,61–63^. In total, 18 structurally novel RBP-classes included 40 RBPs across all morphotypes (Figure S6).

In addition to RBP class sharing across morphotypes, we also observed co-occurrence of distinct structural features across RBP-classes. For example, HRP29 tail-like variants share an N-terminal anchor with the HRP29 gp44 tailspike adapter while diverging structurally overall; the SGNH hydrolase domain appeared in both podovirus tailspikes and myovirus tail fibers; and the RaK2 gp526-like class combined structural elements with precedents across three morphotypes. Whether such sharing is widespread and reflects common ancestry or convergence is examined systematically in the next section.

### Structural Space of RBPs is Shaped by Sharing of Homologous Domains

The structural atlas presented in Figure 2 revealed that structurally related RBP-classes recur across phylogenetically distant phages, and that discrete structural modules are shared across otherwise distinct RBP architectures. Whether this reflects convergent evolution toward structurally optimal solutions, or divergent evolution from shared ancestral domains that have since diversified beyond detectable sequence homology, is a fundamental open question in phage biology — one that cannot be resolved from structural similarity alone, as sequence information is needed to establish true homology^64–68^. This question has been extensively debated in the context of capsid and tail evolution, where the recurrence of a small number of folds across distantly related phages is now widely interpreted as evidence for deep common ancestry rather than convergence^66,69^. We hypothesised that the same logic applies at the RBP level: that recurring structural classes reflect modular reuse of a limited set of ancestral domain building blocks, rather than independent invention.

To test the evolutionary divergence hypothesis, we mapped all RBP structures against the ECOD database^48^, which classifies domains using both structural and sequence information organised into a four-level hierarchy: Architecture (A), X-group (X), Homologous superfamily (H), and Topology (T). Critically, domains placed in the same H-group are inferred to share common ancestry based on both structural and sequence evidence — making ECOD classification a direct test of homology rather than mere structural similarity. Mapping was performed at the topology level with a ≥50% domain-length coverage threshold to ensure that only structurally well-supported matches were retained (see Methods). If the same H-group domains recur across structurally distinct RBP-classes, this constitutes evidence for divergent evolution through domain shuffling rather than convergent invention.

The results of the ECOD mapping were consistent with the hypothesis of divergent evolution through modular reuse. At least one ECOD domain was identified in 302 RBP structures spanning 27 of 39 RBP-classes, with complete annotation in 18 RBP-classes and partial annotation in nine (Figure S7). In 24 of these 27 RBP-classes, members shared at least one ECOD domain with a structurally distinct RBP-class, indicating that the domain repertoire underlying *Klebsiella* phage RBPs is drawn from a common, extensively recycled pool rather than independently assembled (Figure 3A). First, N-terminal anchor domains showed the broadest sharing: domains 3856.1.1 (Putative tailspike protein Orf210 N-terminal domain) and 1083.1.1 (Phage T4 gp12 N-terminal repeating units) were each reused across up to seven RBP-classes spanning multiple morphotypes, including otherwise unrelated architectures, consistent with conserved attachment interfaces that permit downstream modular exchange (Figure S8). Second, enzymatic domains exhibited more restricted but positionally flexible reuse: the 2007.5.1 (SGNH hydrolase domain) occurred across podovirus and myovirus RBPs and in different structural positions, while the 207.2.1 (pectin lyase-like domain) was conserved within depolymerases across RBP-classes (Figure 3A, 3C). Third, C-terminal receptor-binding modules showed extensive non-homologous substitution: carbohydrate-binding domains (e.g. 10.1.2 (Galactose-binding domain-like), 3386.1.1 (gp9 C-terminal domain-related), 10.3.1 (TNF-like)) recurred across multiple RBP-classes and could substitute within a shared structural framework, even adopting different oligomeric arrangements (Figure 3C, Figure S9, Figure S10). Finally, structural scaffold domains further support a shared evolutionary origin: the 79.1.1 (Phage tail fiber protein trimerization domain) and 3240.1.1 (Intramolecular chaperone in virus tail spike protein (IMC)) were reused across multiple RBP-classes and morphotypes and occurred in both tailspikes and tail fibers (Figure 3A, Figure S11), indicating common folding and assembly machinery and supporting a structural continuum rather than independent evolution of tailspikes and tail fibres. Together, these patterns provide quantitative evidence that RBP diversity is driven by domain shuffling from a shared evolutionary pool, supporting divergence rather than convergent invention.

**Figure 3:**
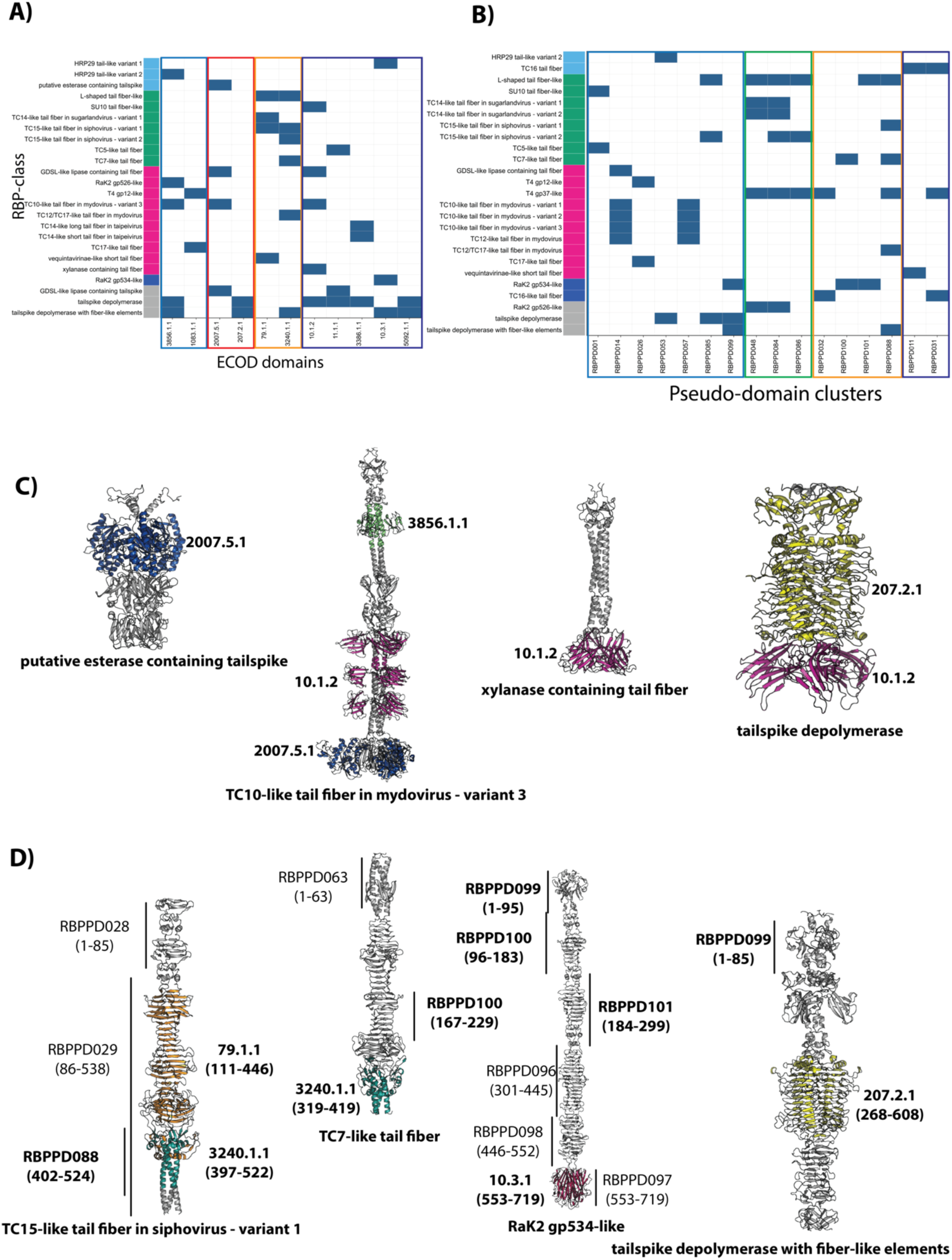
Modular evolutionary architecture of *Klebsiella* phage RBPs revealed through ECOD and pseudo-domain mapping. Comprehensive mapping of ECOD domains and pseudo-domain clusters across RBP-classes. (A) Heatmap of ECOD domains that occur in more than one RBP-class. ECOD domains are displayed as their respective short-hand IDs (3856.1.1 Putative tailspike protein Orf210 N-terminal domain, 1083.1.1 Phage T4 gp12 N-terminal repeating units, 2007.5.1 SGNH hydrolase, 207.2.1 Pectin lyase-like, 79.1.1 Phage tail fiber protein trimerization domain, 3240.1.1 Intramolecular chaperone in virus tail spike protein, 10.1.2 Galactose-binding domain-like, 11.1.1 Immunoglobulin/Fibronectin type III/E set domains/PapD-like, 3386.1.1 gp9 C-terminal domain-related, 10.3.1 TNF-like, 5092.1.1 Domain in virus attachment proteins. (B) Heatmap of pseudo-domain clusters shared across multiple RBP-classes. In A and B, RBP-classes are ordered by their occurrence in different phage morphotypes: podovirus (blue), siphovirus (green), myovirus (pink), jumbophage (dark blue), and RBP-classes spanning more than one phage morphotype (grey). ECOD domains in A and pseudo-domain clusters in B are sorted and shown in the following order of their location - N-terminal (blue, in A & B), enzymatic (red, only in A), knob (green, only in B), scaffold (orange, in A & B) and CBMs (violet, in A & B). Note that in A RaK2 gp526-like is present along with myovirus, and in B it is shown as present in more than one phage morphotype respectively. This discrepancy corresponds to the RBP-class member on which the ECOD and pseudo-domain clusters are mapped. (C) Examples of ECOD domains shared between a few RBP-classes. D) Examples of pseudo-domain clusters mapped onto representative RBPs - coordinates of pseudo-domain clusters and ECOD domain mapped are shown in parentheses. Pseudo-domain clusters shown in bold are the ones present in more than one RBP-classes and shown in panel B of the figure.

Despite this extensive domain sharing, a substantial fraction of RBP structure remained unannotated: 80 RBPs across 21 RBP-classes lacked any detectable ECOD domain, and 57% of all modelled residues fell outside current ECOD classification (Figure S12). This gap is likely the result of both incomplete database coverage—especially for phage tail fiber folds and trimeric assemblies dependent on quaternary structure—and divergence beyond current homology detection limits. As a result, a large portion of RBP structural diversity remains inaccessible to ECOD-based mapping alone. To address this, we applied pseudo-domain segmentation, which partitions each structure into contiguous regions of similar confidence and spatial proximity without relying on predefined domain boundaries or database matches, using a recently proposed approach^30^. These regions were then clustered by pairwise structural similarity using Foldseek at a multimer TM-score threshold of 0.65, and mapped back on the RBP structural models to reveal a pattern of presence and absence of pseudo-domain clusters across all representative RBP-classes (see Methods)^47,51,70^.

Pseudo-domain segmentation revealed modular structural units beyond those captured by ECOD and increased the apparent connectivity of RBP-classes. In total, 102 pseudo-domain clusters were identified across 184 RBPs spanning 38 of 39 RBP-classes, with only a single class (TC14-like short tail fiber in *Taipeivirus*) excluded due to insufficient segmentation signal (Figure S13). Of these, 16 pseudo-domain clusters were shared across multiple RBP-classes, collectively linking 25 RBP-classes across all four phage morphotypes (Figure 3B). This level of cross RBP-class sharing exceeds that observed with ECOD mapping alone, indicating that RBPs are more extensively interconnected at the structural level than suggested by homology-based classification.

Comparison of pseudo-domain clustering with ECOD mapping revealed three consistent patterns. First, multiple pseudo-domain clusters could map to the same ECOD domain, reflecting divergence within homologous folds: for example, RBPPD002 and RBPPD062 both correspond to the 10.1.2 (Galactose-binding domain-like), while RBPPD104 and RBPPD097 map to 10.3.1 (TNF-like), yet were separated due to pairwise multimer TM-scores below the 0.65 clustering threshold. In these cases, TNF-like domains retained TM-scores of 0.5–0.65, consistent with shared fold identity but distinct structural arrangements, whereas galactose-binding domains diverged further (TM ≈ 0.4), indicating more extensive remodelling (Table S3)^48,51,70,71^. Second, pseudo-domain clustering recovered recurrent structural units absent from ECOD classification, including N-terminal regions, C-terminal modules, knob domains, and triple-stranded beta-helix scaffolds; notably, three triple-stranded beta-helix pseudo-domain clusters (RBPPD032, RBPPD100, RBPPD101) were shared across seven RBP-classes, highlighting their role as widely reused structural building blocks (Figure 3B, 3D). Third, a subset of pseudo-domain clusters mapped cleanly onto known ECOD domains, with five pseudo-domain clusters overlapping curated annotations (Table 1), including RBPPD088 corresponding to 3240.1.1 (IMC) and RBPPD097 to 10.3.1 (TNF-like), demonstrating that the approach can also recover established domain units. These three patterns confirm that pseudo-domain segmentation both extends and corroborates ECOD-based annotation.

**Table 1:** Five pseudo-domain clusters shared between multiple RBP-classes that were mapped on to an ECOD domain and RBP-classes in which such cases were detected.

| Pseudo-domain cluster | ECOD domain | RBP-class |
| --- | --- | --- |
| RBPPD011 | 79.1.1 Phage tail fiber protein trimerization domain | Vequintavirinae-like short tail fiber |
| RBPPD026 | 1083.1.1 Phage T4 gp12 N-terminal repeating units | T4 gp12-like; TC17-like tail fiber |
| RBPPD053 | 3856.1.1 Putative tailspike protein Orf210 N-terminal domain | Tailspike depolymerase; HRP29 tail-like variant 2 |
| RBPPD057 | 3856.1.1 Putative tailspike protein Orf210 N-terminal domain | TC10-like tail fiber in mydovirus - variant 3 |
| RBPPD088 | 3240.1.1 Intramolecular chaperone domain in virus tail spike protein | TC15-like tail fiber in siphovirus - variant 1; TC12/TC17-like tail fiber in mydovirus; TC7-like tail fiber; tailspike depolymerase with fiber-like elements |

Taken together, these results show that ECOD mapping and pseudo-domain clustering provide complementary perspectives on RBP architecture. ECOD identifies homologous domains and anchors interpretation in evolutionary relationships but is limited by database coverage and detection thresholds, whereas pseudo-domain clustering extends structural coverage by detecting recurrent units irrespective of prior annotation, at the cost of lacking direct functional or evolutionary assignment. When integrated, the two approaches reveal a highly modular organisation of *Klebsiella* phage RBPs, in which both known and previously uncharacterised structural units are extensively reused and recombined across classes and morphotypes. Yet structural modularity alone does not tell us whether the domain composition of an RBP has functional consequences — specifically, whether particular domain architectures independently shape the host-range breadth of the phage that encodes them. This is the question we address next.

### Pectin lyase-like Topology is the Best Predictor of Narrow Host-Range in *K. pneumoniae* Phages

Having established that RBP diversity is shaped by modular reuse, we next asked whether this structural repertoire relates to host-range. To assess this, we visually inspected ECOD domain architectures in RBPs from narrow and I/B host-range phages by mapping domain positions along each protein (Figure 4). This revealed a single clear pattern: the near-exclusive association of the 207.2.1 (pectin lyase-like) domain with narrow host-range RBPs. In contrast, RBPs from I/B phages displayed substantially greater architectural heterogeneity, largely driven by myoviruses and jumbophages. Overall, we observed 52 domain combinations across 382 RBPs, with a larger share (n=34) in narrow host-range phages and only a subset shared between categories (n=13), but beyond the pectin lyase signal, no consistent domain-level patterns emerged. Instead, domains co-occurred in complex, non-redundant combinations across structurally distinct RBPs, making it difficult to infer individual domain contributions from visual inspection alone. We therefore applied a statistical approach to disentangle domain-specific effects.

**Figure 4:**
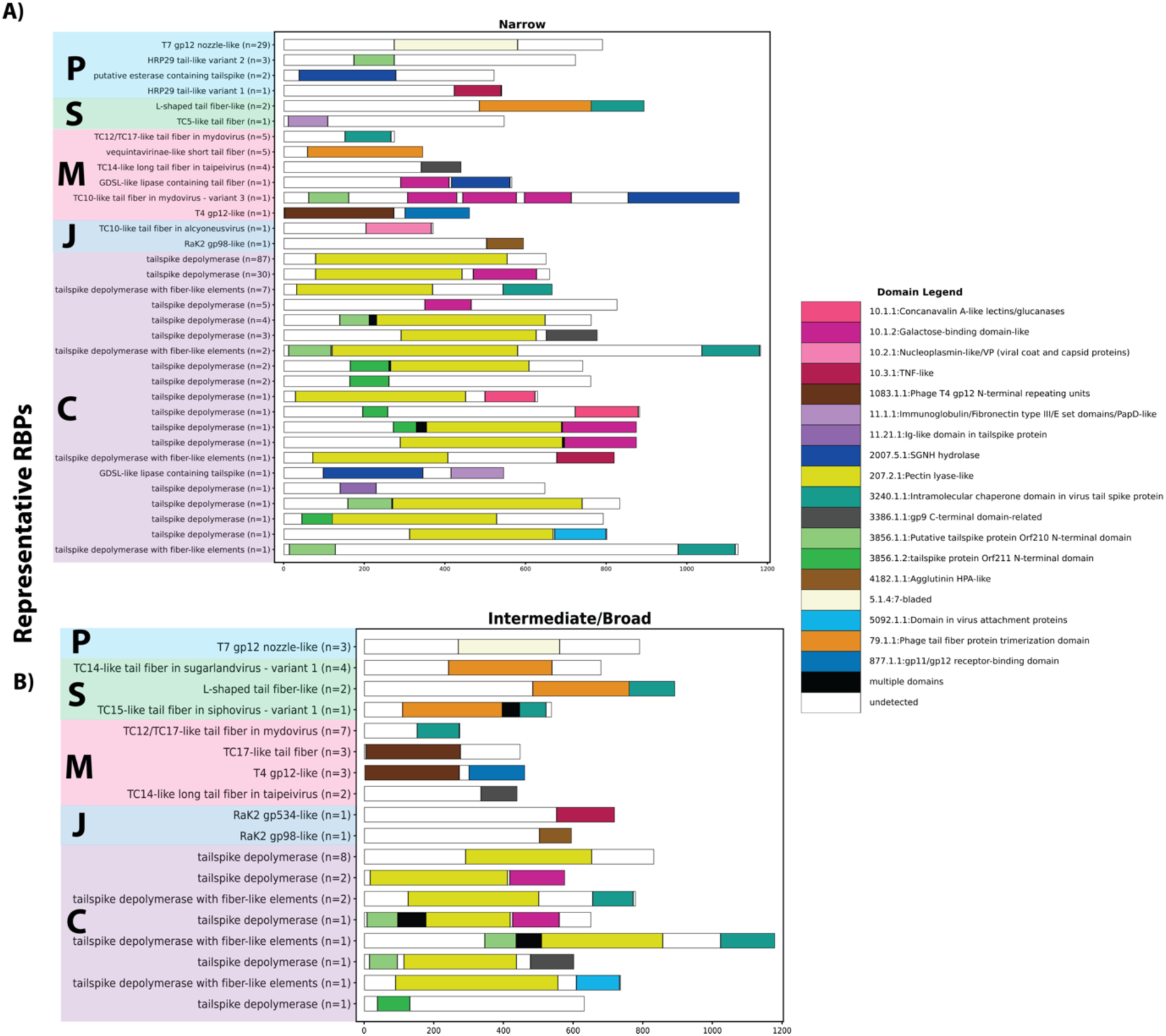
Distribution and combinatorial diversity of ECOD domains across RBPs partitioned by empirical host-range. Schematic representation mapping the presence and linear position of assigned ECOD domains within representative RBPs from phages exhibiting (A) narrow host-range (≤2 K-types) and (B) I/B host-range (≥3 K-types). Individual RBP-classes on the y-axis are grouped and shaded according to their originating phage morphotype: podovirus (P), siphovirus (S), myovirus (M), jumbophage (J), and classes common across multiple phage morphotypes (C).

To quantify domain-level associations with host-range, we fitted a generalized linear mixed-effects model (GLMM) with host-range (narrow vs I/B) as the binary response, domain presence/absence (in ≥5 phage clusters) as fixed effects, and phage cluster (proxy for taxonomic genus) as a random intercept to account for shared evolutionary history. Domains occurring exclusively in one host-range category were excluded from the model as association cannot be estimated without outcome variation. As expected, the 207.2.1 (pectin lyase-like) domain showed the strongest positive association with narrow host-range (OR = 31, 95% CI: 6.8–143; Table 2), reflecting dataset composition dominated by narrow host-range phages carrying SSBH-fold depolymerases, and therefore serving as a validation of model behaviour rather than a novel finding. In contrast, the 3240.1.1 (Intramolecular chaperone in virus tail spike protein) (IMC) domain showed a significant negative association with narrow host-range (OR = 0.13, 95% CI: 0.03–0.60; Table 2). RBPs containing IMC domains are enriched in phages with larger genomes (Figure S17), consistent with more complex receptor-recognition strategies involving either capsule-independent targeting or multiple RBPs and depolymerases. Supporting this, chi-squared tests revealed that IMC domains were overrepresented both in phages lacking SSBH-fold depolymerases (p = 0.01 vs those with 1–2 SSBH-fold depolymerases) and in phages encoding more than two SSBH-fold depolymerases (p < 0.001 vs those with 1–2 depolymerases), confirming that the positive IMC association with I/B host-range reflects a genuine biological signal rather than an indirect consequence of depolymerase content.

**Table 2.** Fixed-effect results from a binomial GLMM relating phage host-range categories (narrow vs I/B empirical host-range) to ECOD domain content in RBPs. Host-range is coded as 1 for narrow and 0 for I/B and modelled with a binomial error distribution and a logit link. Predictors indicate the presence versus absence of ECOD domains detected in at least five phage clusters; Phage cluster is included as a random intercept to account for relatedness among genomes. The table reports odds ratios and corresponding 95% confidence intervals derived from the fixed-effect coefficients, together with p-values for each predictor.

| term | odds.ratio | conf.low | conf.high | p.value |
| --- | --- | --- | --- | --- |
| (Intercept) | 1.05 | 0.46 | 2.4 | 0.90 |
| Galactose binding domain like | 1.26 | 0.18 | 8.7 | 0.81 |
| Intramolecular chaperone domain in virus tail spike protein | 0.13 | 0.03 | 0.60 | 0.008 |
| Pectin lyase like | 31 | 6.8 | 143 | <0.001 |
| Putative tailspike protein Orf210 N terminal domain | 0.75 | 0.09 | 6.5 | 0.80 |
| Tailspike protein Orf211 N terminal domain | 5.37 | 0.36 | 81 | 0.22 |

The remaining commonly detected domains — 3856.1.1 (Putative tailspike protein Orf210 N-terminal domain), 3856.1.2 (tailspike protein Orf211 N-terminal domain), and 10.1.2 (Galactose-binding domain) — showed no association with host-range. This was confirmed using a modified GLMM using genetic rather than empirical host-range classification (Figure 1B), with phages carrying 1–2 SSBH-fold depolymerases classified as narrow and those with 0 or >2 SSBH-fold depolymerases as I/B, and with the pectin lyase-like domain excluded from predictors. The results were consistent with the empirical host-range model: the IMC domain (3240.1.1) remained significantly negatively associated with the narrow host-range (OR=0.03, 95% CI: 0.004-0.21; Table S4), while the other domains showed no effect.

Together, these results point to two structural routes to host-range breadth. The association of pectin lyase-like domains with narrow host-range confirms the expected constraint imposed by capsule-specific depolymerases. In contrast, the enrichment of IMC domains in I/B phages suggests a link to more complex receptor-recognition strategies, including capsule-independent targeting or the use of multiple RBPs and depolymerases. Although IMC itself functions in trimer assembly rather than receptor binding, its independent association with broader host range — after controlling for shared evolutionary history — indicates that more complex RBP architectures are disproportionately associated with expanded host-range, suggesting that structural organisation may actively shape receptor-recognition strategies.

### Sequence Modularity of RBPs extends across different RBP-classes and clear domain boundaries

ECOD and pseudo-domain analyses operate on structural similarity, which captures deep evolutionary relationships but can miss more recent exchange events where sequences have not yet diverged enough to alter the overall fold. Sequence-level comparison complements these approaches because partial sequence matches — in which only a region of two otherwise distinct RBPs is conserved — are a direct readout of mosaicism: the patchwork homology that arises when segments are swapped between proteins by recombination^31^. Here we asked how widespread such sequence mosaicism is across the full RBP repertoire, what forms it takes, and whether its boundaries align with the structural domain organisation established in the previous section.

To detect and classify sequence mosaicism, we performed all-versus-all BLASTP comparisons across all 382 RBP sequences, retaining only pairs where both query and target aligned over at most 70% of their respective lengths^72^. This coverage filter excludes full-length homologues — overwhelmingly within RBP-class pairs sharing overall sequence identity — and enriches for cases where only a specific region is conserved between two otherwise distinct proteins. Detected pairs were classified into five modularity categories based on the position and extent of the shared region (Figure 5A): N-terminal sharing (conserved scaffold anchoring region, boundary set at 250 aa); C-terminal variation (shared scaffold but diverging receptor-binding tip); C-terminal sharing (conserved receptor-binding tip, boundary set to at most last 382 aa); conserved internal regions (shared segments between N- and C-terminal regions); and putative recombination hotspots (short high-identity segments ≤35 aa, ≥80% identity, informed by the hotspot cutoffs defined by Pas et al.)^29^ (Figure S18). This classification provides a conceptual framework for interpreting shared regions rather than a precise delineation of domain boundaries; category boundaries should be regarded as approximations.

**Figure 5:**
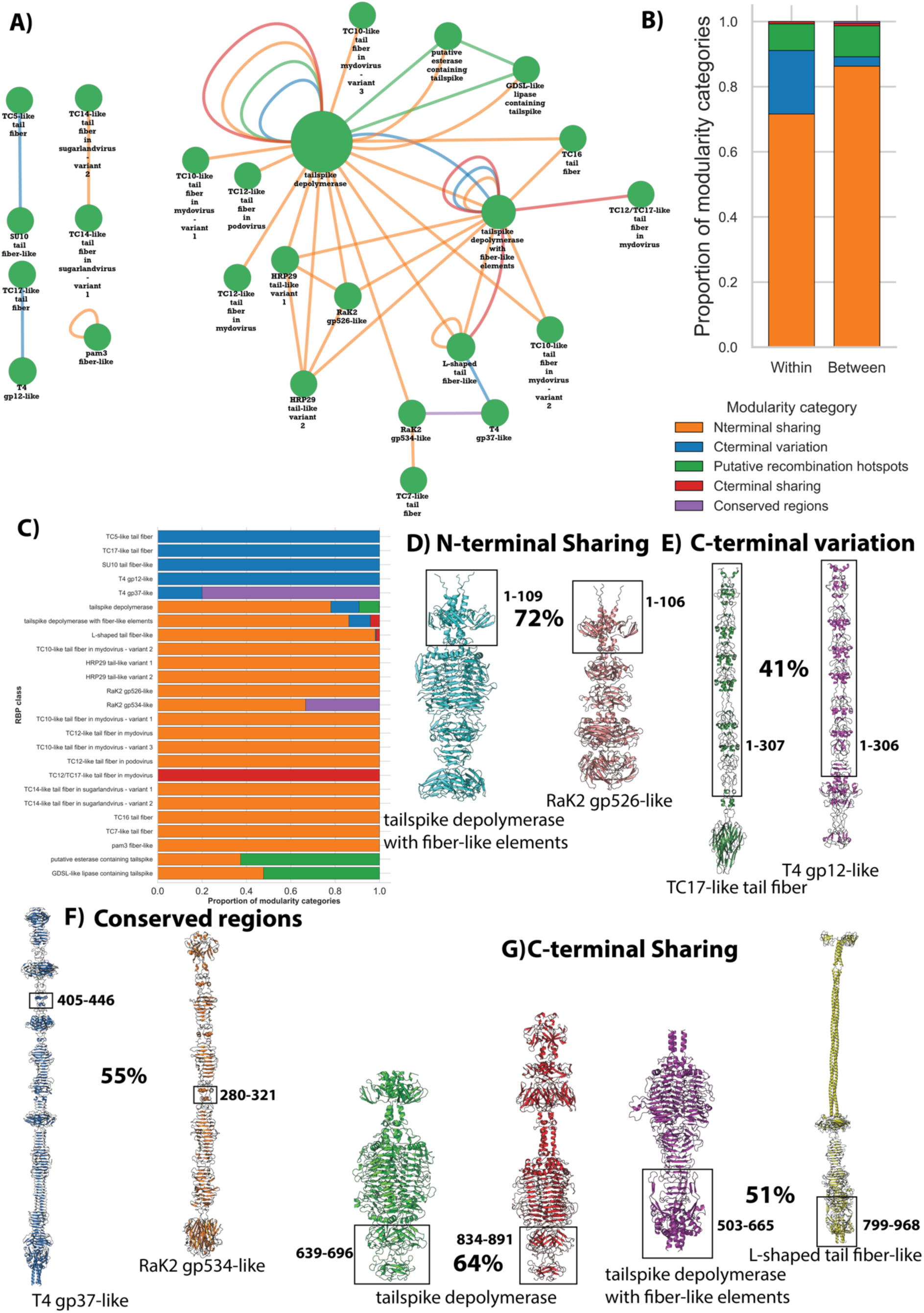
Sequence modularity and structural mapping across RBP-classes. (A) Sequence modularity network where each node represents an RBP-class and node size reflects class membership. Edges connect RBP-classes with evidence of sequence modularity (<70% query/target coverage), and edge colors denote the specific type of modularity category (N-terminal sharing, C-terminal sharing, C-terminal variation, conserved regions, or putative recombination hotspots), consistent with the color scheme in the subsequent panels. (B) Proportion of modularity categories depending on whether they are detected within or between RBP-classes. (C) Relative proportion of modularity categories per RBP-class. (D–G) Illustrative examples of RBP pairs with specific modularity categories mapped onto their structures. Note that the boxes represent approximate structural regions that share sequence similarity, and the exact coordinates where the sequences are similar are shown adjacent to the structures.

Sequence similarity was detected in 1472 RBP pairs involving 245 of 382 RBPs, spanning 25 of 39 RBP-classes (Figure 5A & 5C). Of these, 185 RBPs shared similarity with at least one RBP within the same RBP-class and 199 with at least one RBP from a different RBP-class — indicating that inter RBP-class exchange is approximately as common as within RBP-class similarity at the sequence level. This compares to 17 of the 24 ECOD-domain-sharing RBP-classes (out of 27 RBP-classes with any ECOD annotation) and 20 of the 25 pseudo-domain-cluster-sharing RBP-classes (out of 38 RBP-class with any pseudo-domain clusters detected) showing sequence modularity. Of the 25 RBP-classes with detected modularity, 14 also shared both ECOD domains and pseudo-domain clusters with other RBP-classes, 3 shared only ECOD domains, 6 shared only pseudo-domain clusters, and 2 shared neither. This indicates that the two approaches capture largely overlapping subsets of modular RBP-classes, We detected modularity in additional 2 RBP-classes (TC12-like tail fiber in podovirus & pam3 fiber-like) previously undetected by other methods, suggesting that sequence-level mosaicism captures a similar breadth of inter RBP-class connectivity but at a more recent evolutionary timescale. Three RBP-classes exhibited three distinct modularity categories each — tailspike depolymerase, L-shaped tail fiber-like, and tailspike depolymerase with fiber-like elements — indicating that modularity is not confined to a single structural position within these RBP-classes.

The distribution of modularity categories differed markedly between within and between RBP-class pairs (Figure 5B). Specifically, N-terminal sharing accounted for ∼87% of between RBP-class pairs, whereas C-terminal variation accounted for ∼20% of within RBP-class pairs, consistent with conserved N-terminal scaffolds anchoring RBPs to the virion while receptor-binding tips diversify under selection. C-terminal sharing was rare between RBP-class pairs (0.8%), indicating that convergence on the same receptor-binding solution across distinct RBP architectures is uncommon. Putative recombination hotspots occurred at similar frequencies within and between RBP-class pairs (∼9% and ∼10%; Figure S19) and were confined to the first 100 amino acids regardless of RBP-class (Figure S20), arguing against conserved structural motifs and instead supporting their role as preferred recombination substrates. After normalising for RBP-class size (Figure 5C), 7 RBP classes retained more than one modularity category, confirming that modularity extends across the non-depolymerase repertoire. Notably, SGNH hydrolase-containing classes showed among the highest relative frequencies of putative recombination hotspots, a pattern that may warrant further investigation given their accessory enzymatic roles in *Przondovirus* and *Mydovirus*.

The dominance of N-terminal sharing between RBP-classes has a striking implication. Of the 23 non-depolymerase RBP-classes with detectable sequence similarity, 14 shared N-terminal similarity with at least one depolymerase class. Because these regions correspond to the N-terminal scaffold rather than receptor-binding or enzymatic domains, this supports the idea that both RBP types are assembled on a common N-terminal chassis that maintains compatibility with the baseplate or tail apparatus regardless of the downstream module. This plug-and-play organisation would allow a phage to swap a depolymerase for a structurally unrelated RBP, or vice versa, while preserving virion assembly. For example, a tailspike depolymerase with fiber-like elements from *Taipeivirus* shares 72% N-terminal sequence identity with a RaK2 gp526-like RBP from the same genus, yet diverges completely beyond this region (Figure 5F). The remaining nine non-depolymerase classes showed more restricted modularity, limited to C-terminal variation, conserved internal regions or N-terminal sharing among each other. This suggests that a single recombination event at the shared N-terminal scaffold could shift a phage from capsule-specific, K-type-constrained recognition to a categorically different receptor-recognition strategy with expanded host-range potential.

Mapping sequence modularity onto RBP structures showed that the sequence and structural modularity are not always in perfect accordance. In some cases, boundaries align: the C-terminal variation between TC17-like and T4 gp12-like maps precisely to ECOD domain 1083.1.1 (Phage T4 gp12 N-terminal repeating units), indicating domain-level recombination where intact structural units are exchanged (Figure 5E). However, this alignment is not universal. A conserved internal region shared between RaK2 gp534-like and T4 gp37-like corresponds to an alpha-helical segment with no ECOD annotation (Figure 5D), while C-terminal sharing between tailspike depolymerase with fiber-like elements and L-shaped tail fiber-like extends beyond the IMC domain into an unannotated triple-stranded beta-helix fold (Figure 5G). Similarly, a pair of tailspike depolymerases share 64% identity at their receptor-binding tips, with the shared region spanning domain boundaries (Figure 5G). Together, these examples show that sequence mosaicism does not consistently follow annotated domain boundaries, pointing to a possible role of sub-domain level recombination in RBP evolution.

Taken together, sequence-level analysis shows that RBP modularity operates across multiple structural scales: from shared N-terminal scaffolds spanning depolymerase and non-depolymerase RBP-classes, through C-terminal diversification within RBP-classes, to short recombination hotspots clustered at N-terminal ends. This modularity is not neutral — exchanging receptor-binding modules at any scale can directly alter bacterial targets and host-range breadth. No single method captures this fully: structural approaches miss recent exchange where folds are preserved, while sequence approaches miss deep homology after divergence. We therefore turned to genus-level comparisons to identify clear cases where modular exchange is linked to shifts in receptor specificity and host-range in phages with experimental data.

### Diverse RBP modularity in different genera underpins host-range variation

RBP exchange within phage genera is a recognised driver of host-range variation, and cross-morphotype exchange of depolymerase RBDs has been reported, suggesting that receptor-recognition modules can traverse deep phylogenetic boundaries^14,27–29,42^. However, prior cases were identified without broader genomic context, leaving open whether they reflect recent transfer or ancient shared ancestry, and whether non-depolymerase RBPs undergo equivalent exchanges. Here we present four cases drawn from genus-level genomic comparisons combined with structural modelling and experimental host-range data, representing distinct exchange types, timescales, and functional consequences^30,46,72,73^. These four cases represent distinct RBP exchange modes: a within-genus swap between depolymerase and non-depolymerase RBPs causing a host-range shift (*Webervirus*); within-genus RBD diversification expanding receptor repertoire without category change (*Sugarlandvirus*); cross-morphotype depolymerase sharing at low identity indicating ancient exchange (*Przondovirus*/*Alcyoneusviru*s); and high-identity cross-morphotype sharing consistent with recent exchange and shared capsule tropism (*Webervirus*/*Mydovirus*).

Within genera, we identified near-identical phages diverging specifically at the RBP locus, with the substituted RBP directly mapping to differences in host-range or receptor specificity. The most striking case is *Webervirus*: all 38 phages belong to a single phage cluster but differ at the RBP locus, encoding either a tailspike depolymerase or an L-shaped tail fiber-like protein (Figure 6A). These share 85% N-terminal identity, indicating a precise swap boundary, but have structurally distinct RBDs: an SSBH-fold depolymerase versus a non-enzymatic triple-stranded beta-helix fiber. This difference correlates directly with phenotype: depolymerase-encoding phages are narrow host-range, whereas tail fiber variants are broad, showing that a single RBD substitution can shift host-range category. A related pattern is seen in *Sugarlandvirus* (Figure 6B), where two RBP variants share 66% N-terminal identity but carry distinct RBDs, both supporting broad host-range. Here, the difference may lie in receptor specificity: one variant is predicted to additionally target O-antigens (O1v1, OL14 of *Klebsiella michiganensis* and *Klebsiella oxytoca* respectively)^45^, potentially expanding receptor repertoire without altering host-range category.

**Figure 6:**
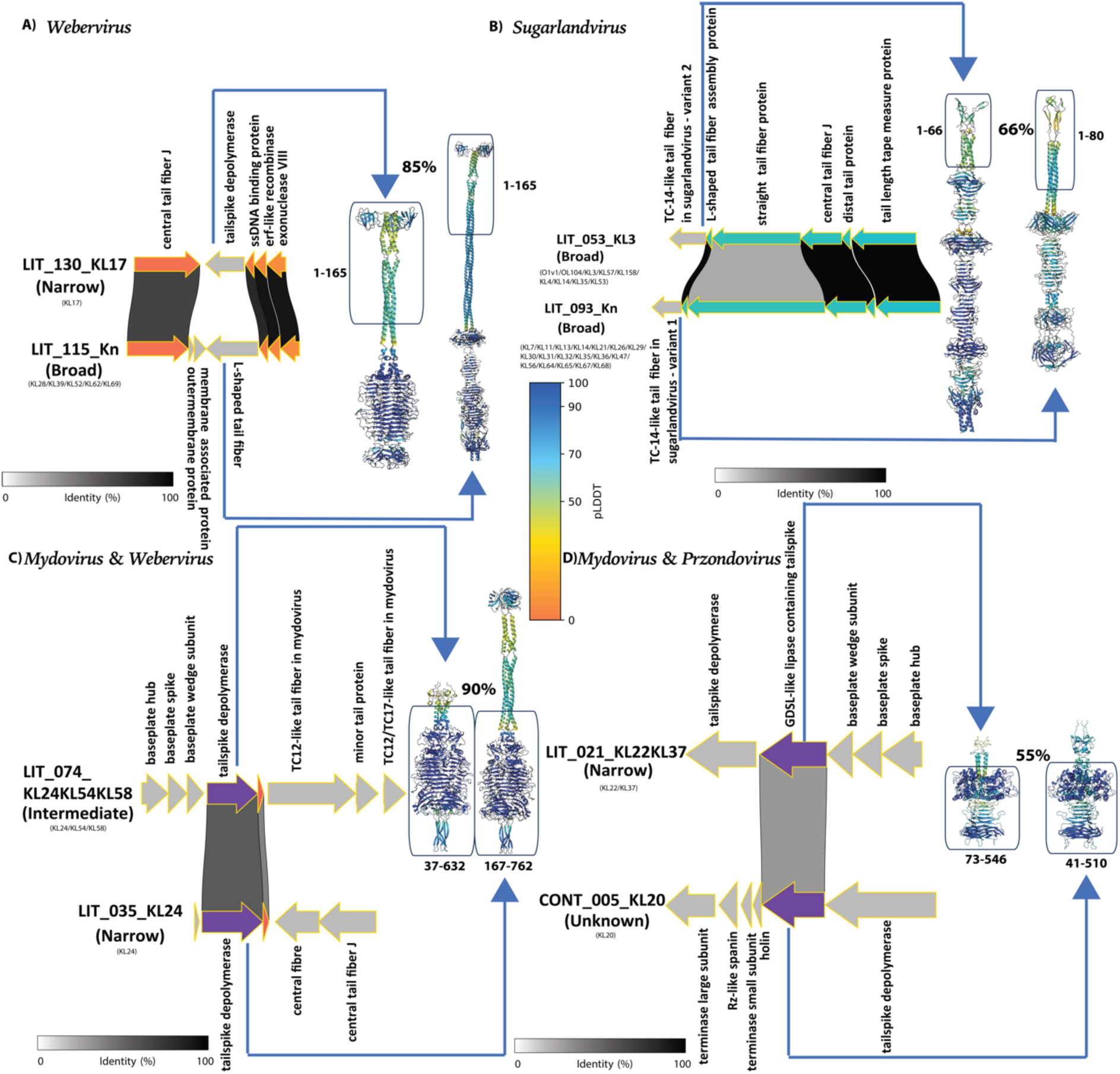
Recombination-driven modular evolution of RBPs within and between *Klebsiella* phage genera and its association with host-range. Panels show RBP gene neighborhoods (drawn to scale; arrows indicate ORFs and of transcriptional direction) and pairwise amino acid identity across the locus (grey-scale shading, 0–100% identity), alongside predicted RBP structures colored by AF3 confidence (pLDDT scale). Square boxes drawn between two RBPs represent the region with amino acid sequence similarity. (A) within *Webervirus* (siphovirus), (B) within *Sugarlandvirus* (siphovirus), (C) between *Przondovirus* (podovirus) & *Alcyoneusvirus* (jumbophage) and (D) between *Mydovirus* (myovirus) & *Przondovirus* (podovirus). Note that the boxes represent approximate structural regions that share sequence similarity, and the exact coordinates where the sequences are similar are shown adjacent to the structures.

We next examined RBD sharing across genera and morphotypes. Cross-morphotype sequence similarity was not restricted to tailspike depolymerases but was observed across all four RBP classes, totalling 982 pairs from 382 RBPs (Figure S21). Tailspike depolymerases contributed the majority (778 pairs; 697 with <50% identity and 81 with ≥50% identity), involving 160 of 186 proteins (86%), consistent with their abundance (49% of all RBPs). Two cases illustrate different timescales. A podovirus (*Przondovirus*) and a jumbophage (*Alcyoneusvirus*) share 47% RBD identity and KL64 specificity (Figure S22), indicating ancient exchange with conserved function. In contrast, *Webervirus* siphoviruses and *Mydovirus* myoviruses share 90% RBD identity and KL24 specificity (Figure 6C), consistent with recent exchange.

Tailspike depolymerases with fiber-like elements showed comparably high cross-morphotype connectivity — 30 of 35 proteins (86%) formed 199 pairs (177 with <50% and 22 with ≥50% identity) — despite representing only 9% of all RBPs (Figure S23). RaK2 gp526-like and GDSL-like lipase-containing tailspikes each contributed a small number of pairs (3 and 2, respectively), with 2 of 3 proteins in each class involved, suggesting that cross-morphotype RBP sharing, while rare, extends to structurally distinct classes (Figure 6D, Figure S21, Figure S24). Together, these examples show not only that receptor-recognition modules can be transferred across genera and morphotypes, but also that such recombination events are ongoing in populations of *Klebsiella* phages and drive their rapid host range adaptation.

Taken together, these cases show that RBP exchange in *Klebsiella* phages is not restricted to within-genus or within RBP-class, nor to depolymerases alone. It operates at two levels: within genera, where near-identical phages differ at the RBP locus and a single RBD substitution can shift host-range category, expand receptor repertoire, or alter KL-type specificity; and across genera and morphotypes, where shared host selection drives convergent receptor-recognition solutions in distant lineages, with sequence identity reflecting exchange recency. The functional outcomes are consistent but graded, spanning host-range shifts, receptor expansion, and specificity changes. In this light, the structural modularity described above — including conserved N-terminal scaffolds, a shared domain inventory, and recombination hotspots clustered at the N-terminal boundary — is not merely a passive record of ancestry. Instead, it constitutes the molecular architecture that enables these exchanges: a permissive substrate for host-range adaptation operating continuously across multiple, nested scales.

## Discussion

The dominant view in *K. pneumoniae* phage biology has long emphasised capsule-specific recognition: depolymerases breach the polysaccharide barrier of an encapsulated host^27,42^ — a framework well-supported by the dominant phage literature but one that leaves unexplained a recurring subset of capsule-independent phages with broad empirical host-range^10,74^. Our dataset recapitulates this: 145 of 192 phages carry one or two SSBH-fold depolymerases and largely display narrow empirical host-range, but also contains 34 phages encoding no detected SSBH-fold depolymerase that nonetheless form plaques or even halos on encapsulated hosts, and a further subset whose empirical host-range exceeds what their SSBH-fold depolymerase content alone could predict. These are not incidental recoveries: the same depolymerase-independent genera recur across two independent collections assembled under conditions that actively select against them^43,45^. Explaining their presence requires asking not just what RBPs *Klebsiella* phages carry, but why the *K. pneumoniae* surface sustains so many structurally distinct infection strategies.

The answer lies in the dynamic heterogeneity of the host surface. Capsule thickness and density vary markedly across and within K-types^38–40,75,76^, and 71% of *K. pneumoniae* strains exhibit within-culture capsule heterogeneity attributable to genetic phase variation^39^. Capsule expression is further reduced by the nutrient-rich conditions used in standard phage enrichment: Buffet et al. report 89% capsule loss in LB over three days^40^. Phage predation compounds this by selecting for capsule-loss escape mutants^77–79^, which persist in nutrient-rich conditions where capsule expression is itself costly^40^. Together, these dynamics mean that any phage isolation experiment exposes a fraction of the bacterial population with reduced or absent capsule, inadvertently creating conditions that can select for phages capable of targeting sub-capsular receptors^44^. Critically, these sub-capsular targets — including OMPs such as OmpA, OmpC and FhuA and the conserved LPS inner core — are under strong purifying selection across *K. pneumoniae* lineages^9,32,44^, making them structurally stable, cross-K-type accessible. These dynamics are amplified when phage enrichment uses a diverse panel of hosts or K-types, further enriching for phages with sub-capsular receptor strategies. The result is that broad host-range phages can be recovered even from conditions that nominally select for K-type specialists, which explains their recurrence across independent collections. The concurrent maintenance of specialist and generalist strategies in this system mirrors the broader coevolutionary pattern in which host diversification sustains phage diversity across ecological timescales^5,6^.

The atlas maps these regimes onto five receptor recognition strategies, each defined by its primary recognition target: K-type dependent (strategies 1-2) and K-type-independent (strategies 3-5) (Figure 7). Most receptor-level assignments for non-depolymerase RBP-classes are inferred from structural homology to characterised *E. coli* phage RBPs; direct experimental evidence for receptor specificity in *Klebsiella* phages is only beginning to emerge and is noted where available. For the three strategies whose primary recognition target lies below the capsule layer (Strategies 3–5), whether the capsule physically facilitates, impedes, or is irrelevant to infection is a mechanistic question that remains to be resolved.

**Figure 7:**
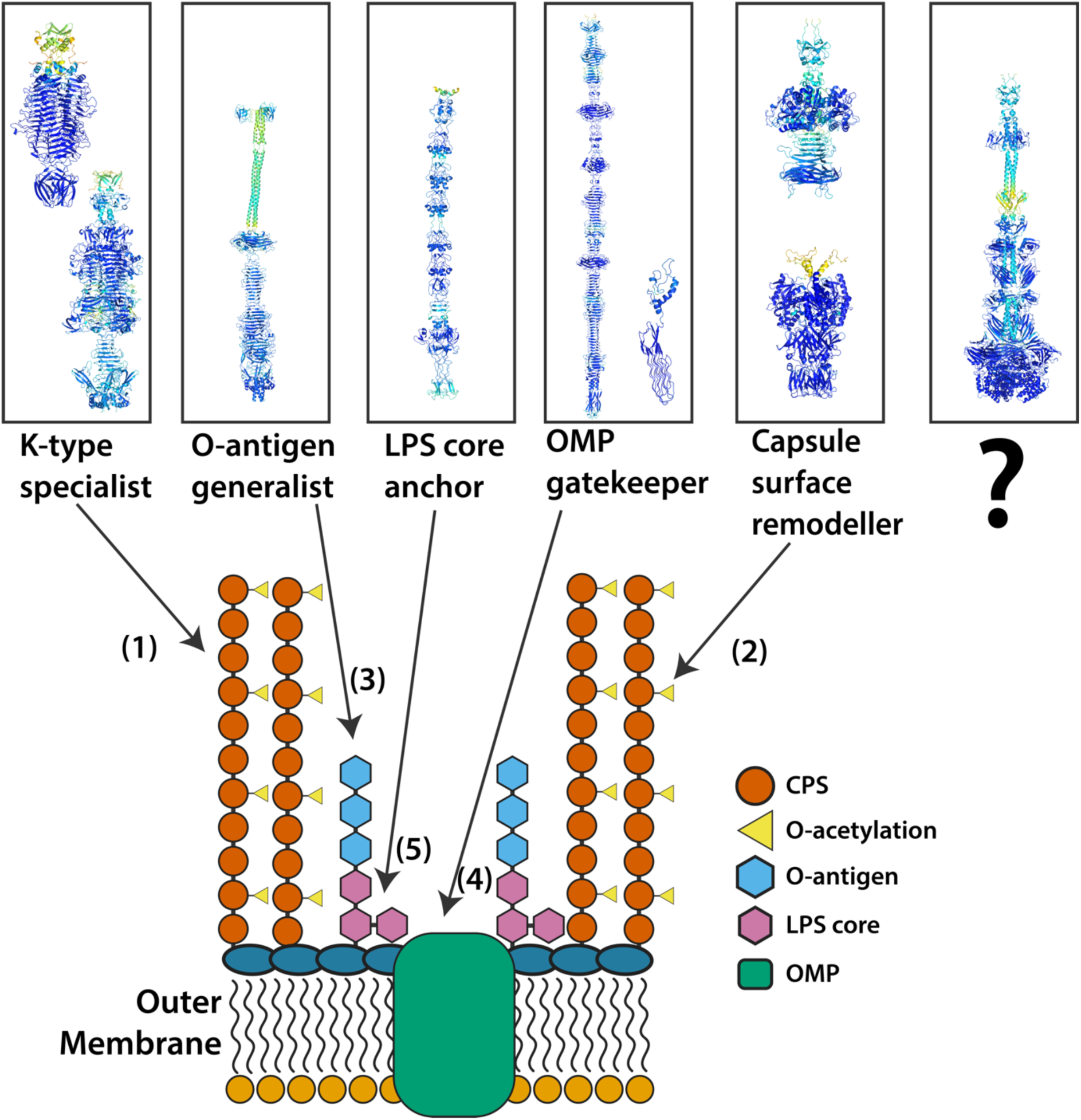
A five-strategy structural atlas of receptor recognition in *Klebsiella* phages. (**Top**) Representative AlphaFold3-predicted structural models of receptor-binding proteins (RBPs) corresponding to each of the five receptor recognition strategies identified in this dataset, coloured by pLDDT confidence score (dark blue, high confidence; cyan, moderate confidence; yellow/green, low confidence). From left to right: an SSBH-fold tailspike depolymerase (Strategy 1, K-type specialist; encompassing tailspike depolymerase and tailspike depolymerase with fiber-like elements RBP-classes); an L-shaped tail fiber-like RBP (Strategy 3, O-antigen generalist); a T4 gp12-like short tail fiber (Strategy 5, LPS core anchor); the gp37 receptor-binding tip of the gp34–gp37 long tail fiber complex alongside a pam3-like adhesin (Strategy 4, OMP gatekeeper); and two capsule surface remodeller RBPs (Strategy 2) — a GDSL-like lipase-containing tailspike (top) and a putative esterase-containing tailspike (bottom). A sixth panel (question mark) represents the RBP-classes sharing no detectable structural similarity with any characterised protein, whose receptor specificities and infection strategies remain entirely unknown. (**Bottom**) Schematic cross-section of the *K. pneumoniae* cell envelope, illustrating the spatial targets of each strategy relative to the layered host surface architecture. The outer membrane is embedded with OMPs (green rectangles). Above the outer membrane, the LPS inner core (pink hexagons) and O-antigen (blue hexagons) extend outward, topped by the densely packed CPS layer (orange circles) decorated with O-acetyl modifications (yellow triangles). Numbered arrows indicate the primary recognition target of each strategy: **(1)** K-type specialist phages (SSBH-fold depolymerases) target the CPS layer in a K-type-dependent manner; **(2)** capsule surface remodellers (SGNH hydrolase-domain RBPs) are proposed to act on O-acetyl or other polysaccharide surface modifications; **(3)** O-antigen generalists (L-shaped tail fiber-like RBPs) target O-antigen residues beneath the CPS; **(4)** OMP gatekeepers engage outer membrane proteins directly, with gp37 forming the receptor-binding tip of the long tail fiber in *Straboviridae* and the pam3-like adhesin providing an architecturally distinct solution in *Jameshumphriesvirinae*; and **(5)** LPS core anchors (gp12-like short tail fibers) are inferred to target the heptose-rich LPS inner core. Strategies 1–2 are K-type dependent; strategies 3–5 operate independently of K-type. For strategies 3–5, whose primary recognition target lies below the capsule layer, whether the CPS physically facilitates, impedes, or is irrelevant to infection remains an open mechanistic question.

Strategy 1 (K-type specialist): phages encoding one or two SSBH-fold depolymerases whose active-site geometry constrains infection to the cognate K-type^7,27,42,80,81^. These are the dominant phages in this and most published *Klebsiella* collections, including the genera like *Drulisvirus*, *Webervirus* and *Przondovirus*. This strategy also includes phages that carry an arsenal of depolymerases: jumbophages like *Alcyoneusvirus* and *Eowynvirus* scale the same enzymatic logic across K-types by encoding eight or nine SSBH-fold depolymerases, respectively ^7,42,82,83^, achieving breadth through RBP accumulation rather than promiscuity (Figure 7).

Strategy 2 (capsule surface remodeler): SGNH hydrolase-domain RBPs including GDSL-like lipase tailspikes and tail fibers, and putative esterase tailspikes, whose characterised homologues, including *E. coli* phage G7C gp63.1 tailspike, experimentally remove O-acetyl modifications from polysaccharide substrates^26^, and for which direct functional evidence in *Klebsiella*’s temperate phages has recently been reported^84^. SGNH hydrolases are however a structurally diverse superfamily; the specific substrates of the novel RBP-classes identified here — whether capsular O-acetyl groups, O-antigen modifications, or other surface glycan decorations — remain to be determined experimentally. A key feature of Strategy 2 is that host-range outcome depends on the full RBP repertoire rather than the esterase/lipase alone: in *Przondovirus*, SGNH hydrolase RBPs co-occur only with SSBH-fold depolymerases and the phage has narrow host-range, whereas in *Mydovirus* they sit alongside SSBH-fold depolymerases and diverse non-capsule-targeting fibers, and the phage has broad host-range. RBP combination rather than individual RBP identity is the phenotypic determinant, a principle likely extending to other multi-RBP architectures in the dataset (Figure 7).

Strategy 3 (O-antigen generalist): L-shaped tail fiber-like RBPs in *Webervirus*, inferred from homology to the T5 L-shaped tail fiber that targets oligo-mannose O-antigen residues in *E. coli*^21^. Consistent with this inference, L-shaped tail fiber-carrying *Webervirus* phages are substantially broader than SSBH-fold depolymerase-carrying members of the same genus, a pattern consistent with limited O-antigen diversity (13 O-loci) imposing far weaker constraints than capsular locus diversity (more than 150 KL-types)^33,34,85^(Figure 7).

Strategy 4 (OMP gatekeeper): phages exploiting this strategy are proposed to operate through OMP-targeted adsorption, but the four genera assigned to this strategy employ architecturally distinct solutions. *Slopekvirus* and *Jiaodavirus* (*Straboviridae*, *Tevenvirinae*) encode the full gp34–gp35-gp36–gp37 long tail fiber complex, engaging OmpC directly via the C-terminal tip of gp37 by close structural analogy with T4^8,86–88^. *Bimevirus* and *Mascletvirus* (*Jameshumphriesvirinae*), phylogenetically distant from *Straboviridae*, entirely lack the gp34–gp37 long tail fiber and instead carry only a pam3-like adhesin with hypervariable C-terminal loops — a structural analogue of the *Straboviridae* gp38 adhesin — paired with a tailspike depolymerase^23,58,59,88^. The recurrence of a gp38-like adhesin module in a phylogenetically distant lineage, decoupled entirely from its ancestral long tail fiber context, is itself an instance of the domain-level modularity that characterises RBP diversification more broadly (Figure 7).

Strategy 5 (LPS core anchor): *Slopekvirus* myoviruses carrying T4 gp12-like short tail fiber are inferred to target the heptose-rich LPS inner core directly^8,89,90^. That 5 of 7 *Slopekvirus* in KlebPhaCol were isolated on capsule-null hosts and 2 of 7 on encapsulated strains is consistent with LPS core access being facilitated by, but not requiring, reduced capsule density or its absence^9^ (Figure 7).

Importantly, these five strategies can be used in combination, leading to more complex infection strategies. This is commonly observed in myoviruses, which accumulate multiple RBP-classes within a single genome. For example, in *Taipeivirus*, SSBH-fold depolymerases are accompanied by TC14-like long and short tail fibers; in *Mydovirus*, eleven RBP-classes co-occur spanning tailspike depolymerases, SGNH hydrolase RBPs, and multiple tail fiber classes. In these phages broad host-range may be an emergent property of RBP repertoire breadth, enabled by the modular domain architecture described below. More than 60% of phages in both genera are empirically I/B host-range; their occasional classification as narrow host-range most likely reflects panel size constraints rather than genuine receptor restriction. The contrast with *Bimevirus* and *Mascletvirus* is potentially instructive: both genera carry a pam3-like OMP-targeting adhesin but pair it exclusively with a tailspike depolymerase rather than an additional non-enzymatic tail fiber. In this configuration, successful OMP engagement would be contingent on either prior capsule loss or sufficient depolymerase-mediated capsule clearance — potentially imposing a K-type dependency at the first step of adsorption and limiting infection breadth to the substrate range of a single enzymatic RBP, regardless of the adhesin’s OMP-targeting capacity. That *Mascletvirus* displays a narrow empirical host-range and *Bimevirus* has unknown empirical host-range is consistent with this interpretation, though direct experimental evidence is required to distinguish an architectural bottleneck from other contributing factors. Finally, 18 RBP-classes in this dataset share no detectable structural similarity with any characterised protein, representing a structural frontier whose receptor specificities and infection strategies remain entirely uncharacterised.

A natural experiment within *Sugarlandvirus* exposes a direct consequence of this receptor diversity. Eight phages, including five from Townsend et al. (23 K-type panel) and three from Ferriol-González et al. (77 K-type panel) are uniformly broad empirical host-range with zero detected SSBH-fold depolymerases, yet two form halos, three are confirmed halo-negative and halo status was not reported for the remaining three^43,45^. This same pattern is replicated in *Jiaodavirus* isolated on KL28^45^. Halo and host-range breadth are therefore regulated by separable molecular processes in these genera. Halo formation requires enzymatic degradation of capsular polysaccharide that diffuses beyond the plaque front^25^; the two halo-positive *Sugarlandvirus* and one *Jiaodavirus* may therefore encode such activity, though alternative explanations — including co-purified phages in the lysate carrying depolymerase activity — cannot be excluded. If the activity is intrinsic to these phages, whether it resides in any of their RBPs or in a non-SSBH-fold depolymerase not detected by current methods remains open.

The persistence of diverse strategies reflects the capacity of RBPs to track receptor heterogeneity through multi-scale modular reshuffling. This modularity is also the structural mechanism that permits multi-RBP generalists to accumulate receptor-binding diversity: N-terminal domain sharing across RBP-classes provides the assembly-compatible scaffold into which new receptor-binding tips can be inserted by recombination. N-terminal domains are shared across 12 of 27 ECOD-annotated RBP-classes while C-terminal tips show marked structural variation within RBP-classes, and putative recombination hotspots cluster exclusively within the first 100 amino acids regardless of RBP-classes, consistent with N-terminal regions serving as assembly-compatible scaffold for RBD exchange^14,28,29^. The IMC domain extends this permissiveness to the folding level: by scaffolding trimeric assembly independently of tip identity, IMC-bearing architectures accommodate tip diversity that would otherwise compromise oligomeric stability, and their enrichment in I/B empirical host-range phages across genera suggests this architectural permissiveness translates into phenotypic breadth, though the causal mechanism remains to be demonstrated^91,92^. That modular substitution can operate even within a single RBP-class is illustrated by *Slopekvirus*, where the T4 gp37-like tail fiber carries an IMC in most phages of the genus but is replaced by an receptor binding adhesin in a minority variant — a swap co-occurring with substitution of the tail fiber assembly protein by a gp38-like adhesin, suggesting C-terminal domain identity co-evolves with the broader assembly pathway, though whether this affects host specificity remains unknown (Figure S25). These features dissolve the classical tailspike-tail fiber binary, placing both on a structural continuum assembled from a common domain inventory^66^. The N-terminal domain-sharing principle is most directly illustrated in *Webervirus*, where near-identical genomes toggle between an SSBH-fold tailspike depolymerase and an L-shaped tail fiber-like RBP at a single locus with 85% flanking N-terminal identity, shifting between narrow and broad host-range through a single modular substitution. At deeper phylogenetic scales, *Mydovirus* and *Webervirus* share 90% RBD identity in KL24-specific tailspike depolymerases, showing that receptor-recognition modules flow across morphotypic and taxonomic boundaries under sustained shared host selection.

Standard isolation recovers this diversity unevenly. K-type specialist (strategy 1) phages form plaques efficiently on encapsulated lawns; phages employing the K-type-independent strategies 3-5, while may be capable of infecting encapsulated hosts directly, form plaques less efficiently under standard conditions because the encapsulated majority presents lower sub-capsular receptor accessibility, and are systematically outcompeted during enrichment. The 34 depolymerase-independent phages in our dataset cannot be minority-cell specialists: clean plaque formation requires repeated productive infection cycles at the plaque front, which is impossible if infection is confined to rare phenotypic variants. Their recurrence across independent collections confirms ecological success rather than fortuitous recovery. The directional effect of isolation protocol on this diversity is quantified most clearly by comparison with KlebPhaCol, a community-assembled collection isolated under conditions designed to broaden receptor access^9^. KlebPhaCol enriched 32 sequence-type-diverse clinical strains in LB — conditions under which substantial capsule loss occurs^40^ — and included five capsule-null strains among the isolation hosts, with a 32 K-type test panel. By contrast, Ferriol-González et al. enriched only on the K20 reference host, recovering 10 of 73 broad host-range phages (14%), and Townsend et al. enriched across all hosts, recovering 11 of 14 (79%)^43,45^. The outcome of the KlebPhaCol approach is stark: 44 of 52 phages (85%) were broad host-range^9^. The 18% depolymerase-independent I/B frequency in our dataset — recovered under conditions that actively select against these strategies — is therefore a lower bound on their true environmental prevalence.

A prediction and a therapeutic design principle follow. Multi-strain or acapsular host enrichment protocols, particularly in nutrient-rich media, will recover O-antigen generalist, OMP gatekeeper, LPS core anchor and multi-RBP generalist phages at higher frequency and broader architectural diversity than single-strain protocols, as the difference between KlebPhaCol and conventional enrichment directly demonstrates^9,43–45^. For phage therapy, K1 and K2 hypervirulent K-types present a specific challenge: reduced capsule heterogeneity and lower spontaneous capsule loss rates^39,40^, mean standard isolation yields K-type specialist dominated collections, and recovering O-antigen generalist, OMP gatekeeper and LPS core anchor against these clinically critical K-types will likely require engineered acapsular host derivatives or ST-pooled enrichment designed to access the sub-capsular receptor landscape^9,44^. The structural atlas, host-range integration and proposed five-strategy taxonomy together provide the first systematic structural classification of RBP diversity in *Klebsiella* phages across 39 RBP-classes, a structure-informed framework for positioning future isolation, engineering and therapeutic efforts against the known receptor landscape of *K. pneumoniae*.

Several limitations constrain the structural and mechanistic conclusions drawn from this analysis. The structural analysis under-samples very long fibers, multicomponent complexes and RBPs requiring partner proteins for stable folding. Fifty-seven percent of RBP residues fall outside ECOD annotations, reflecting genuine structural novelty in phage-specific folds and incomplete database coverage of trimeric assemblies. Discrete host-range bins from heterogeneous panels obscure efficiency-of-plating gradients, making genera such as *Mydovirus*, *Taipeivirus*, *Bimevirus*, and *Mascletvirus* susceptible to apparent narrow empirical host-range classification that likely reflects panel constraint. A further caveat concerns accessory adhesins encoded separately from the primary RBP. In T-even phages other than T4 (T2 & T6), and *Salmonella* S16 phage long tail fibers, gp37 carries an IMC domain and undergoes auto-proteolytic cleavage upon trimerisation, but critically lacks an intrinsic C-terminal receptor-binding domain; receptor specificity is instead provided by gp38, a separately encoded adhesin that associates with the fiber tip^23,59,88^. The pam3-like adhesins identified in this dataset are structural homologues of *Straboviridae* gp38 and represent this RBP-class^58,59^. IMC-bearing fibers and tailspike with fiber-like elements that similarly lack a C-terminal CBM or adhesin domain are therefore likely to follow the same paradigm, relying on undetected external adhesin partners for receptor engagement. Structural analysis of individual polypeptides is by definition blind to such partners, and their prevalence among IMC-bearing RBP classes in the dataset is almost certainly under-represented, reflecting the limited characterisation of this interaction partner in the broader phage literature. Resolving the identity and receptor specificity of these putative adhesins is a tractable and high-priority avenue for future mechanistic studies. Finally, strategy assignments for receptor recognition and the IMC-to-breadth mechanistic interpretation are most-parsimonious structural conclusions consistent with the data; they are not demonstrated mechanisms, and direct receptor-level experimental validation is the immediate priority.

## Methods

### Data

We compiled 192 virulent *Klebsiella* phage genomes from published literature and phages isolated from unfiltered sewage from a wastewater treatment plant in Manchester. All the phages in this study are experimentally tested for their bacterial capsular specificities, and were reported to demonstrate a halo and/or a plaque on a lawn.

#### Public host-range collections

This dataset contains 175 phage genomes for which the host-range of the phages are described based on the formation of either a halo & plaque or just plaques. We assembled published genomes from three host-range collections originating from distinct studies^41,43,45^, totaling 117 genomes, 44 additional genomes of phages with experimental data for depolymerase activity^42^ and 14 genomes curated from nine additional articles^93–101^. To assemble the dataset, we searched PubMed for studies on phages infecting *K. pneumoniae* that reported depolymerase activity and experimentally determined host-range (plaques and/or halos), using combinations of keywords such as ‘bacteriophage’, ‘phage’, ‘*Klebsiella pneumoniae*’, ‘depolymerase’, ‘capsular polysaccharide’, ‘host-range’, ‘plaque’, and ‘halo’. We further selected only those phages for which infection assays demonstrated plaques with or without halos, and we exceptionally also included phages for which at least one depolymerase had been heterologously expressed and its K-type specificity experimentally determined.

#### Phage Isolation and Propagation

Phage isolation from enrichment culture was performed following Blundell-Hunter G. et. al., 2021^102^. Eighty-six *K. pneumoniae* phages were isolated from raw sewage collected in 2017 at the Davyhulme wastewater treatment plant, Manchester, United Kingdom. Isolation was performed using an international panel of 96 *K. pneumoniae* strains comprising multi-, extended-, and pan-drug-resistant clones obtained from bioMérieux. For enrichment, 10 ml sewage aliquots were supplemented with 0.3 g TSB powder, inoculated with 100 μl overnight bacterial culture, and incubated at 37°C overnight with orbital shaking at 200 rpm. Cultures were centrifuged at 4,000 rpm for 20 minutes and supernatants filtered through 0.45 μm membranes (Merck Millipore) before storage at 4°C.

Individual plaques were recovered by preparing ten-fold serial dilutions (10⁻¹ to 10⁻⁸) of each filtered supernatant and applying the double-layer agar method. Overnight bacterial culture (100 μl) was combined with molten top agar, overlaid onto TSA plates, and 10 μl of supernatant spotted onto the surface before overnight incubation at 37°C. Plaques were excised with a sterile tip, resuspended in 300 μl sterile SM buffer, and stored at 4°C overnight. Single-plaque purification was carried out over three successive rounds; in each cycle, serial dilutions prepared in SM buffer were used to inoculate TSB containing 200 μl logarithmic-phase culture, mixed with top agar, and poured onto TSA plates for overnight incubation at 37°C. Following the final purification cycle, phage stocks were amplified by adding 10 μl of purified phage to 10 ml logarithmic-phase bacteria in TSB and incubating overnight at 37°C at 200 rpm. Cultures were centrifuged at 4,000 rpm at 4°C for 30 minutes and supernatants collected by 0.45 μm filtration before storage at 4°C.

#### Genomic DNA Extraction and Sequencing

Genomic DNA was extracted from both bacterial hosts and phages using a phenol:chloroform:isoamyl alcohol (PCI)/SDS protocol. Fresh phage lysates at 10⁷–10⁹ pfu/ml were prepared by the flood plate method. For each phage, 1.5 ml of lysate was centrifuged at 13,000 rpm for 10 minutes at 4°C, and 1 ml of clarified supernatant was treated with DNase I (10 μl of 1 mg/ml) and RNase A (4 μl of 12.5 mg/ml) to remove residual host nucleic acids. DNA was then isolated by sequential extraction with phenol (pH 10), phenol:chloroform (1:1), and phenol:chloroform:isoamyl alcohol (25:24:1 v/v), each step involving 30 seconds vortexing and centrifugation at 13,000 rpm for 10–20 minutes at 4°C. DNA was precipitated overnight at −20°C with two volumes of ice-cold absolute ethanol and one-tenth volume of 7.5 M ammonium acetate, pelleted at 13,000 rpm for 20 minutes at 4°C, washed twice with 70% ethanol, air-dried, and resuspended in 100 μl nuclease-free water at 37°C. Purity was assessed by 260/280 absorbance ratio on a NanoDrop One spectrophotometer (Thermo Scientific), targeting a ratio of approximately 1.8. Final yields were quantified in triplicate using a Qubit 3.0 Fluorometer (dsDNA High Sensitivity assay, Life Technologies) and DNA normalised to approximately 5 ng/μl before dilution to 0.2 ng/μl for library preparation. Sequencing libraries were prepared with the Illumina Nextera XT kit and reads assembled using metaSPAdes^103^. We successfully sequenced and assembled 59 *K. pneumoniae* host genomes, representing 19 MLST sequence types and 20 distinct KL-types, and identified 58 virulent phages at greater than 100× coverage.

#### Phage Genome Selection and Quality Filtering

From the 58 sequenced phage genomes (>100x coverage), 20 candidates displaying clear lytic activity restricted to a single KL-type were shortlisted. For 17 of these, the predominant contig per lysate was selected on the basis of highest assembly coverage; three were excluded because no contig satisfied the predefined quality thresholds of coverage ≥ 1,000× and genome length ≥ 30 kb^104^. Among the 17 retained phages, nine displayed siphovirus morphotype and eight podovirus morphotypes (inferred based on gene synteny and genome similarity to phages from literature), all with genome coverage ≥ 1,000× and length ≥ 30 kb. These 17 phages collectively targeted nine distinct capsular types: K3, K10, K17, K20, K22, K53, K54, K58, and KL127.

Phage specificity was performed only on 3 K-types (including isolation host K-type), but this was insufficient to contribute information towards empirical host-range. Hence, these 17 phages are assigned Unknown empirical host-range and are excluded from empirical host-range comparisons, but are retained in genetic host-range classification and all structural RBP diversity analyses.

Altogether, we analyzed 192 virulent phages with evidence of capsular specificity tested against a range of 96 K-types. All genomes from the literature were retrieved from GenBank using their accessions (Table S1), except *Klebsiella* phage SH-KP156570 (GWHBGZN01000000), which was downloaded from the Genome Warehouse (https://ngdc.cncb.ac.cn/gwh/).

#### Host-range assignment

Host-range assignment was performed using two different approaches.

##### Genetic host-range classification via SSBH-fold depolymerase repertoire

To assess genetic host-range potential, phages were categorized into discrete cohorts based on the total number of SSBH-fold depolymerases encoded per genome. Initially, proteins harboring depolymerase-associated folds were identified using DepoScope (v.1.0)^105^, with candidates filtered using a stringent confidence threshold (score ≥ 0.95). The presence of the defining SSBH-fold was subsequently confirmed for each candidate by visually inspecting AlphaFold 3 homo-trimer models^46^. This structural validation identified a total of 238 SSBH-fold depolymerases across the dataset. All 238 validated depolymerases were retained for genome-level counting, independent of their overall AlphaFold 3 model quality metrics (full details regarding structural prediction and quality estimation are provided in subsequent sections). Based on these final counts, phages were stratified into four distinct genetic host-range bins: 0, 1–2, 3–4, and ≥5 encoded SSBH-fold depolymerases per genome (Figure 1B).

##### Empirical host-range classification

Since the phages in our dataset were compiled from multiple independent studies, each tested against a variable number of reference K-types, we employed an empirical strategy to standardize host-range assignment. A phage was considered eligible for host-range classification only if it had been tested against at least five distinct K-types; otherwise, its host-range was categorized as Unknown. Among the evaluated phages, those infecting up to two distinct K-types were classified as Narrow, those infecting up to four K-types as Intermediate, and those infecting five or more K-types as Broad (Figure S4). In total, we assigned 127 genomes as Narrow, 12 as Intermediate, 22 as Broad, and 31 as Unknown in terms of host-range.

### Genome analysis

#### Clustering

We computed wGRR for all pairwise genome comparisons using MANIAC in coding-sequence (CDS) mode^106^. We used 0.65 (wGRR cutoff at which all phages belonging to a common genus were grouped in one cluster) as the wGRR cut-off and Markov Clustering (MCL) with inflation=2 to define 40 genomic clusters (Figure S3 & S3).

#### Phage Morphotype

Phage morphotypes were assigned in two ways: Published phages retained their reported morphotypes, while our 17 phages—lacking TEM images—were classified based on gene synteny, verified against phages from published literature.

#### Taxonomy

Taxonomic assignments were generated with tax_myPHAGE (v0.3.3) using default parameters^107^. The phages span 22 distinct genera; 25 phages were designated ‘New_genus’ where tax_myPHAGE did not return a genus-level assignment.

#### Functional annotation

We performed functional annotation of phage genomes using an in-house snakemake workflow. ORFs were predicted with Prodigal-gv and Glimmer, then clustered and aligned with MMseqs2 (min. identity 0.5, coverage 0.8, e-value 0.001, sensitivity 7.5, –msa-format-mode 3)^49,108–110^. MSAs were augmented against PHROGs via HHblits and converted to cluster profiles using HHsearch, queried against ECOD, PFAM, and PHROG (min. probability -p 50, query identity -qid 10, coverage -cov 10, max. e-value -E 1)^48,49,111–113^. Per-cluster consolidation prioritized highest bit-score hits: PFAM (non-DUF first), ECOD (highest bit-score), PHROG (up to two hits with e-value <0.001). This identified 18,028 protein-coding sequences across 192 genomes. The complete pipeline is available at: https://github.com/bioinf-mcb/mgg_annotation/tree/prodigal-gv.

### RBP Detection

From the functional annotation of 192 phage genomes, PHROGs annotations were used to retrieve putative RBP candidates by searching for tail fiber or tailspike entries using a regular expression (r’^\W*tail\W*(?:fiber|spike)\W*protein\W*$’), yielding 329 candidates^49^. In parallel, PhageRBPDetect_v4 was applied to all 18,028 CDSs, selecting candidates with either a binary prediction value of 1 or a normalized score ≥0.95, yielding 538 candidates^50^. From the combined set, only those with a minimum length of 200 aa, a PHROGs RBP annotation, and a PhageRBPDetect_v4 score ≥0.95 were retained as high-confidence RBPs. Central fibers were subsequently excluded using a regex (r"(?:central\s+fibre|central\s+fiber|central\s+tail\s+fiber\s+J)"), as AlphaFold 3 and RBPseg failed to generate reliable models for these proteins, and truncated N-terminal fragments ≤350 aa corresponding to the T7 gp17 N-terminal domain (PFAM (PF03906) corresponds precisely to only the N-terminal of gp17) were also removed, as some podoviruses were found to carry only these fragments (r"Phage T7 tail fibre protei\[Phage_T7_tail\]\[PF03906\.14\]")^114^. This filtering yielded a final dataset of 457 RBP candidates (Figure S23).

### Structure Prediction

#### AlphaFold 3

We used AlphaFold 3 (v.3.0.1) to predict the homo-trimeric structures of all the 457 RBP candidates we predicted. We assigned random model seeds between 1 and 100,000 for each protein and used default recycles (n=10) to ensure reproducible predictions^46^.

#### Refine selected RBP predictions

We used RBPseg to refine the structure predictions of 59 RBPs for which AlphaFold 3 was unable to resolve the structure properly^30^. RBP structure prediction was performed using RBPseg (v1.1.1) and AlphaFold 3 on an NVIDIA A100 GPU.

A trimeric AlphaFold 3 prediction of the RBP was generated and used as input for the RBPseg-sDp module to define structural domains:

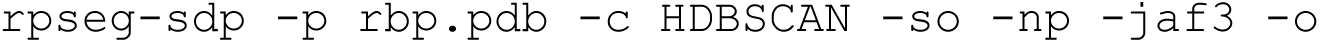

Each identified fraction was predicted individually using AlphaFold 3 with one random seed. Fraction predictions were merged into a full-length model using:

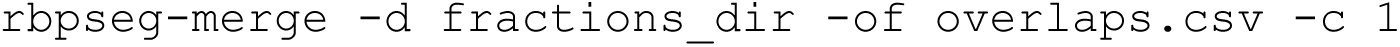

Merged models were screened for structural clashes using rbpseg-validate. Models exhibiting clashes were re-segmented using k-means as the clustering parameter:

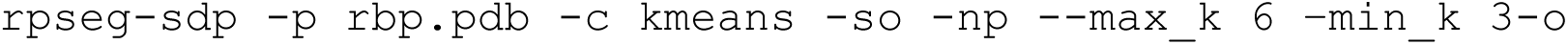

The resulting fractions were re-predicted with Alphafold 3 and merged again using rbpseg-merge as described above.

#### Model Quality Filters

For the 398 RBP structures predicted solely using Alpha Fold 3, we retained only those that met a composite quality threshold defined as a ranking score greater than or equal to 0.5 (Figure S23).

For the 59 RBP structures refined using RBPseg, we calculated the merge score using the RBPseg-validate module^30^. For each structure, we then computed the average merge score across all residues and selected only those with an average merge score greater than or equal to 80.

In total, we shortlisted 410 high-quality RBP structures (356 structures modelled exclusively using Alpha Fold 3 and 54 structures refined using RBPseg and modelled using AlphaFold 3) (Figure S5).

### Structural Analyses

#### Structural clustering of RBPs

We used the Foldseek easy-multimercluster module to cluster the RBP structures at a multimer TMscore threshold of 0.65, a chain TMscore threshold of 0.5, and an interface lDDT threshold of 0.65. The 410 high-quality RBP structures were subsequently clustered into 148 clusters^51^.

#### RBP Classification

The 148 Foldseek clusters formed the basis for our RBP classification scheme. To this end, we defined an “RBP-class” – a group of one or more Foldseek clusters sharing a common overall architecture and, where determinable, an inferred functional identity based on similarity to characterised RBPs. RBP-classes therefore represent discrete structural and functional units within the broader RBP repertoire, operating at a coarser level of granularity than the Foldseek clusters themselves. To assign Foldseek clusters to RBP-classes, we applied a sequential classification pipeline that prioritised confident functional assignments and fell back on progressively more distant structural comparisons where confident matches were unavailable, as described in detail below.

Based on the presence of an SSBH-fold at the central domain, we manually identified 221 depolymerases. Out of which, we classified 186 RBPs as tailspike depolymerase and 35 RBPs as tailspike depolymerase with fiber-like elements based on variations in their C-terminals. The depolymerases in the latter class contained elements resembling tail fibers^27^. We used the RBPseg classify module to assign tail fiber classes (TCs) to the remaining 189 RBPs^30^. Respective TCs were assigned using a minimum probability cut-off of 0.95 and a minimum query TMscore of 0.5. As a result 6 TCs were assigned to 72 RBPs and these were classified into 6 RBP-classes respectively (T4 gp34-like, T4 gp36-like, T7 gp12 nozzle-like, TC16 tail fiber, TC8 tail fiber & *Vequintavirinae*-like short tail fiber).

For the remaining 117 RBPs to which a TC couldn’t be confidently assigned, we ran Foldseek search against PDB100 (v20240101) for each Foldseek cluster representative, and defined the best hit based on the highest TMscore match^47^. A hit was deemed significant only if it had a minimum TMscore of 0.5. Using this approach we defined 9 RBP-classes to 27 RBPs (GDSL-like lipase containing tail fiber, GDSL-like lipase containing tailspike, HRP29 tail-like variant 1, HRP29 tail-like variant 2, L-shaped tail fiber-like, pam3 fiber-like, putative esterase containing tailspike, SU10 tail fiber-like, xylanase containing tail fiber).

Out of the remaining 90 RBPs we defined 4 RBP-classes to 14 RBPs based on similarities to phage RBPs already available in published literature (RaK2 gp526-like, RaK2 gp534-like, RaK2 gp98-like, T4 gp12-like)^53,54^.

For the remaining 76 RBPs without any significant hit to PDB100 or representatives found in literature, we defined RBP-classes based on distant TCs detected with the RBPseg-classify module (probability <= 1 & qTMscore < 0.5)^30^. This way, we assigned 18 RBP-classes with functional assignment to 48 RBPs (T4 gp37-like, TC10-like tail fiber in *Alcyoneusvirus*, TC10-like tail fiber in *Mydovirus* - variant 1, TC10-like tail fiber in *Mydovirus* - variant 2, TC10-like tail fiber in *Mydovirus* - variant 3, TC12-like tail fiber in *Mydovirus*, TC12-like tail fiber in podovirus, TC12/TC17-like tail fiber in *Mydovirus*, TC14-like tail fiber in *Sugarlandvirus* - variant 1, TC14-like tail fiber in *Sugarlandvirus* - variant 2, TC14-like long tail fiber in *Taipeivirus*, TC14-like short tail fiber in *Taipeivirus*, TC15-like tail fiber in siphovirus-variant 1, TC15-like tail fiber in siphovirus - variant 2, TC16-like tail fiber, TC17-like tail fiber, TC5-like tail fiber & TC7-like tail fiber).

Finally, we discarded the remaining 28 RBPs as they were identified as false positives because their HMM hits to PFAMs suggested proteins of function acetyl-CoA acetyltransferase, baseplate tail tube protein, tail tubular protein etc.. Overall, we grouped 382 high-confidence RBPs into 39 RBP-classes (Figure 2, Figure S6, Figure S23 and Text S1).

#### Domain detection in RBPs

We used the Foldseek easy-search module with the 3Di+AA Gotoh-Smith-Waterman method (–alignment-type 2) to map domains onto the high-quality structures of 382 high-confidence RBPs using the ECOD F70 structural database (v20230309)^47,48^. ECOD domain hits were filtered using a minimum target coverage of 50% and an E-value cutoff of 0.001. For each RBP, remaining ECOD hits were then ranked by decreasing bit score, breaking ties in favor of longer query alignments, and iteratively filtered so that any lower-ranked hit whose query interval overlapped an already accepted hit by more than 50% of the shorter domain’s length was discarded, yielding a final set of high-scoring, largely non-overlapping domain assignments per protein. After assigning high-confidence ECOD domains, overlapping intervals corresponding to the same topology within an RBP were merged to generate nonredundant domain annotations for visualization. Overall, we detected 18 unique ECOD domains across 302 RBPs (Figure S7).

#### Pseudo-Domain detection in RBPs

Representative RBP structures from each of the 39 RBP-classes were selected for pseudo-domain segmentation. In most cases, the longest RBP per class was used; however, for the L-shaped tail fiber-like, the second-longest RBP was chosen because the longest one did not form a straight conformation typical of other members of the RBP-class. Pseudo-domain segmentation was performed using the RBPseg-sdp module, with segmentation and clustering executed via HDBSCAN under the following parameters^30^:

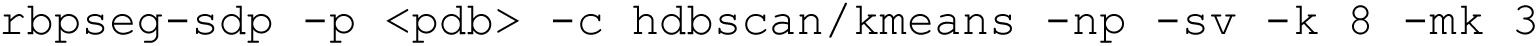

The maximum number of clusters was set to 8, and the minimum cluster size to 3. The resulting 121 pseudo-domains were subsequently clustered by all-against-all structural similarity using Foldseek (v10.941cd33), employing the easy-multimercluster workflow with default settings^47^:

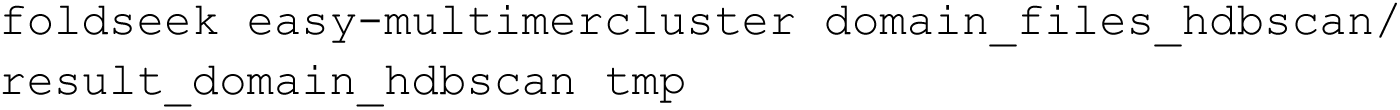

Pseudo-domain clusters were grouped into three categories: 6 cross-protein clusters (present more than one RBP-class), 6 repeat-domain clusters (repeated within the same RBP-class), and 95 orphan domain clusters (present only in one RBP-class). Among the repeat-domain clusters, only one represented genuine repeats; the remaining five were segmentation artefacts and were excluded from further analysis.

### Statistical Modelling

#### Statistical model to predict host-range

Using the 40 phage clusters described above, we retained 36 containing phages with empirical host-range data available, comprising 161 genomes for downstream analysis. Within this subset, we assessed the presence or absence of specific ECOD domains in RBPs, with domain annotations performed as described above. Rare domains, defined as those occurring in fewer than five phage clusters, were excluded to reduce sparsity. The remaining domains were encoded as binary predictors at the genome level.

Associations between ECOD domain content and host-range were analysed using generalized linear mixed-effects models (GLMM) with a binomial error distribution and logit link function, implemented using the glmer function from the lme4 R package. The model estimates the association between the presence or absence of each ECOD domain (T-level) and the probability of a phage displaying narrow versus I/B host-range, while accounting for non-independence among phages with shared evolutionary history (same phage cluster). Host-range was treated as a binary response variable, classifying phages as either having a narrow or an I/B empirical host-range. Phage cluster was included as a random intercept to account for non-independence among genomes within clusters and to capture cluster-level variation in baseline host-range. ECOD domain predictors were included as fixed effects.

The model was specified as follows:

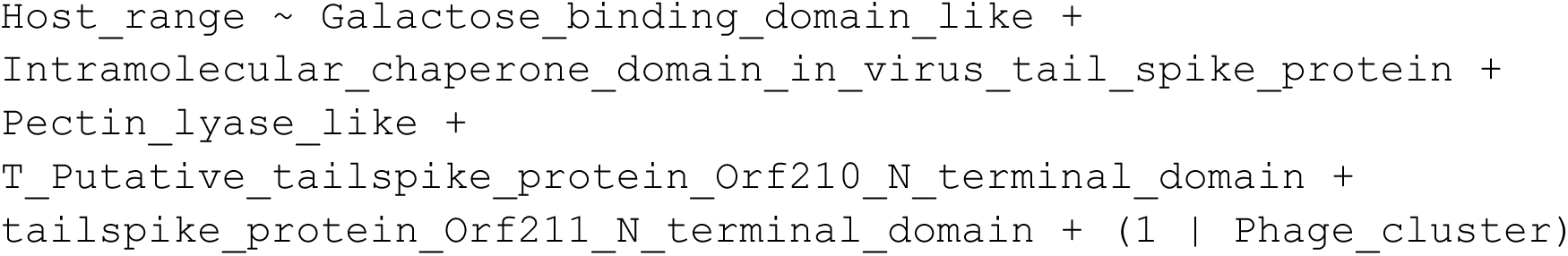

In a GLMM with a logistic link function, model coefficients represent log-odds; exponentiating these yields odds ratios, which quantify the change in odds of narrow versus I/B host-range associated with the presence of each domain.

To investigate the relationship between ECOD domain content and depolymerase-associated host-range, we fitted a modified GLMM:

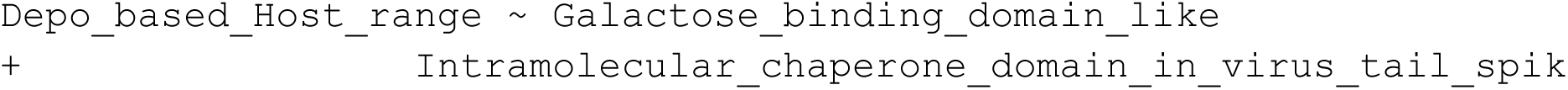
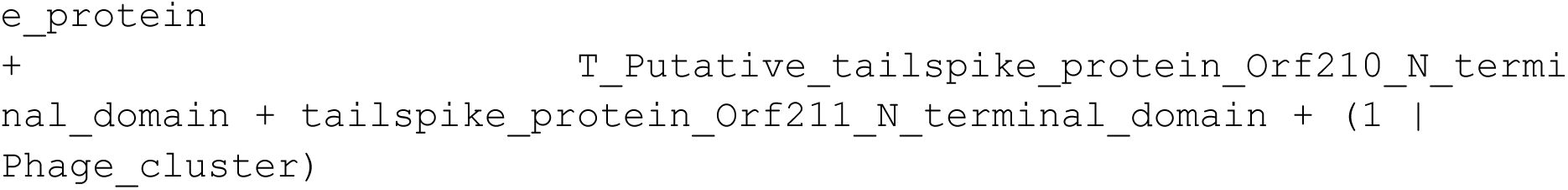

In this model, *Depo_based_Host_range* was defined as “narrow” for phages encoding 1–2 depolymerases, and as “I/B” for phages encoding either no depolymerases or more than two depolymerases.

Finally, chi-squared tests were performed using the chisq.test function in R to evaluate differences in the presence of RBP containing an intramolecular chaperone (IMC) domains between phages with 1–2 depolymerases and: (i) those lacking depolymerases, and (ii) those encoding more than two depolymerases.

### Sequence Mosaicism Analysis

#### Sequence similarity search with BLASTP

We performed an all-against-all BLASTP search on 382 high-confidence RBPs using tabular output format (-outfmt 6)^72^. A dataset-derived threshold N = 382 aa was defined as half the median protein length of the 382 RBPs (median: 763 aa) and applied across multiple filtering schemes below. For modularity categories 1–4, we first removed highly similar RBP pairs by retaining only hits where both query and target coverage were ≤ 70%, and applied an E-value cutoff of ≤ 0.001. We then applied five distinct filtering schemes to capture different modularity categories: N-terminal sharing, C-terminal sharing, C-terminal variation, conserved regions, and putative recombination hotspots.

1. **N-terminal sharing** — RBP pairs assigned to the N-terminal sharing category were detected by selecting hits with sequence identity ≥ 50% and a minimum alignment length of 30 aa (the shortest depolymerase N-terminal identified in the dataset). The aligned fragment was further required to fall entirely within the first 250 aa of both query and target sequences, corresponding to the approximate median N-terminal length of depolymerases in the dataset (237 aa; Table S5).
2. **C-terminal sharing** — RBP pairs assigned to the C-terminal sharing category were identified by selecting hits with sequence identity ≥ 50% and an alignment length of at most 382 aa (N: median protein length of the 382 RBPs). The aligned fragment was additionally required to reach the true C-terminus of both sequences, defined as ending within 10 aa of the terminal residue in both query and target.
3. **C-terminal variation** — RBP pairs assigned to the C-terminal variation category were identified by selecting hits with a minimum alignment length of 250 aa, ensuring that detected similarity extended well beyond the N-terminal segment in both sequences. Both proteins were required to exceed N = 382 aa in length to ensure sufficient internal sequence for an alignable body region. Pairs were further required to have an unaligned C-terminal tail of between 10 aa and 382 aa in both query and target — confirming the presence of a divergent C-terminal region that is neither trivially short nor spanning the full protein.
4. **Conserved regions** — RBP pairs assigned to the conserved regions category were identified by selecting hits with sequence identity ≥ 50% whose aligned segments were located entirely within the internal region of both sequences: beginning after the first 250 aa and ending at least 382 aa (N) before the C-terminus in both query and target.
5. **Putative recombination hotspots** — RBP pairs assigned to the putative recombination hotspots category were identified from the full set of pairwise BLASTP hits (without the ≤ 70% coverage or E-value pre-filters applied to categories 1–4) by selecting short, high-identity matches with sequence identity ≥ 80% and an alignment length of at most 35 aa^29^.

## Author Contributions

The following contributions can be distinguished by different authors based on the CRediT (Contributor Roles Taxonomy) system:

- Conceptualization: VRP, RJM
- Methodology: VRP, BJS, VKS, RJM
- Software: VRP, BJS, VKS
- Validation: VRP
- Formal Analysis: VRP, BJS, VKS
- Investigation: VRP, BJS
- Resources: MCE, NMIT, RJM
- Data Curation: VRP
- Writing – Original Draft: VRP, RJM
- Writing – Review & Editing: VRP, BJS, VKS, MCE, NMIT, ZDK, RJM
- Visualization: VRP, BJS, RJM
- Supervision: NMIT, ZDK, RJM
- Project Administration: ZDK, RJM
- Funding Acquisition: RJM

## Data & code availability

Phage genomes isolated as part of this manuscript are available from NCBI Genank (to be submitted): https://www.ncbi.nlm.nih.gov/genbank/. The annotation pipeline we used is available: https://github.com/bioinf-mcb/mgg_annotation/tree/prodigal-gv. RBP input sequences (FASTA, JSON), AlphaFold 3 models, metadata files generated and all other relevant large data files are available on Zenodo: <to be submitted>. All custom code used for analysis and figure generation is publicly available at: https://github.com/VyshakhRP/RBP-div-hostrange.git. Figures generated in this manuscript are available via figshare: https://figshare.com/s/7b3285de0562781bd745. The *Klebsiella* RBP atlas generated is available as an interactive community resource: https://vyshakhrp.github.io/klebsiella-rbp-atlas/

## Supplementary Figures

**Figure S1:**
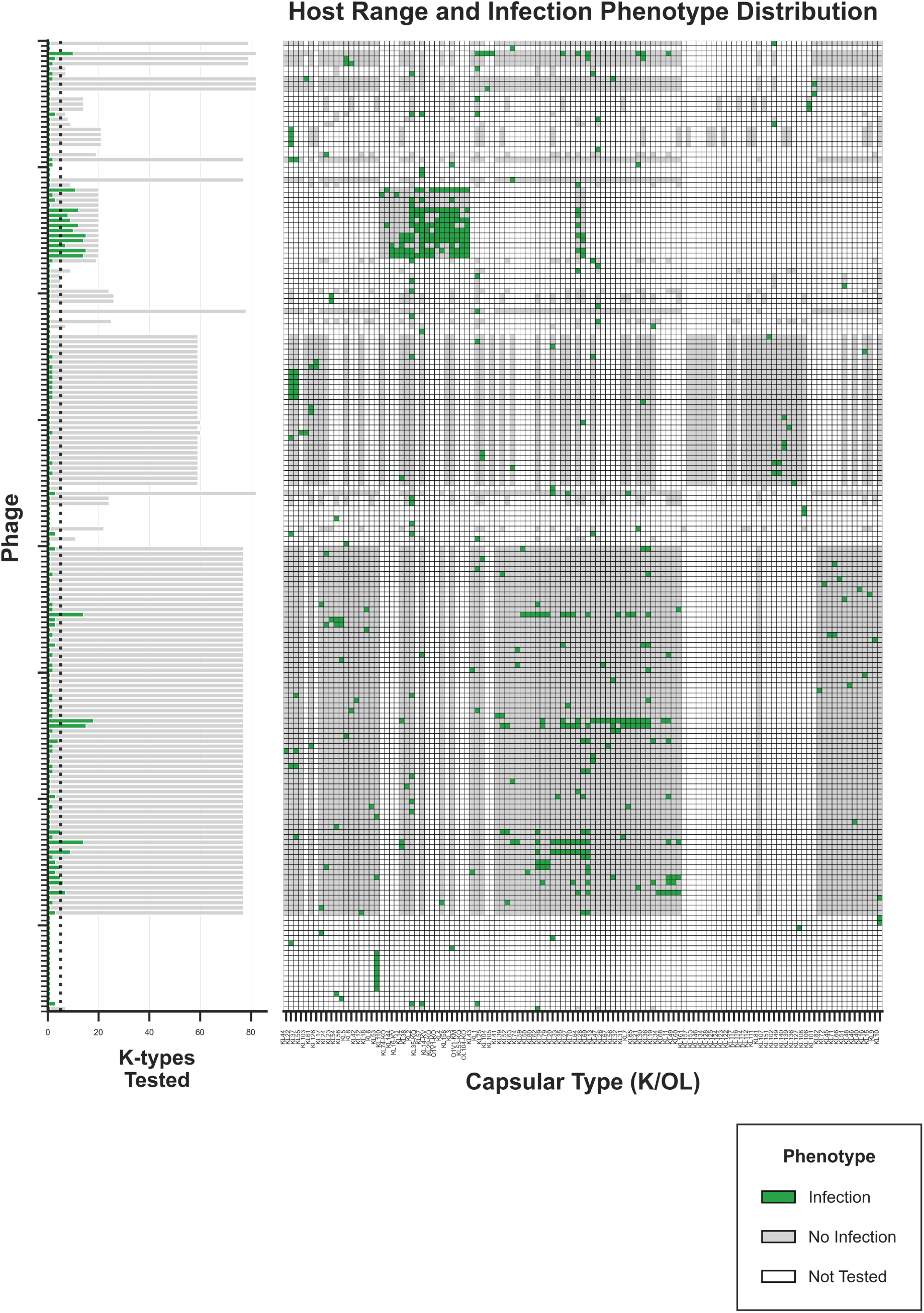
Complete infection panel for the 192 phages analyzed in this study. Host-range data against K-types and O-types were sourced from original publications, except for 17 virulent phages from our collection. The dotted line marks the minimum testing threshold (5 K-types) for assigning empirical host-range categories.

**Figure S2:**
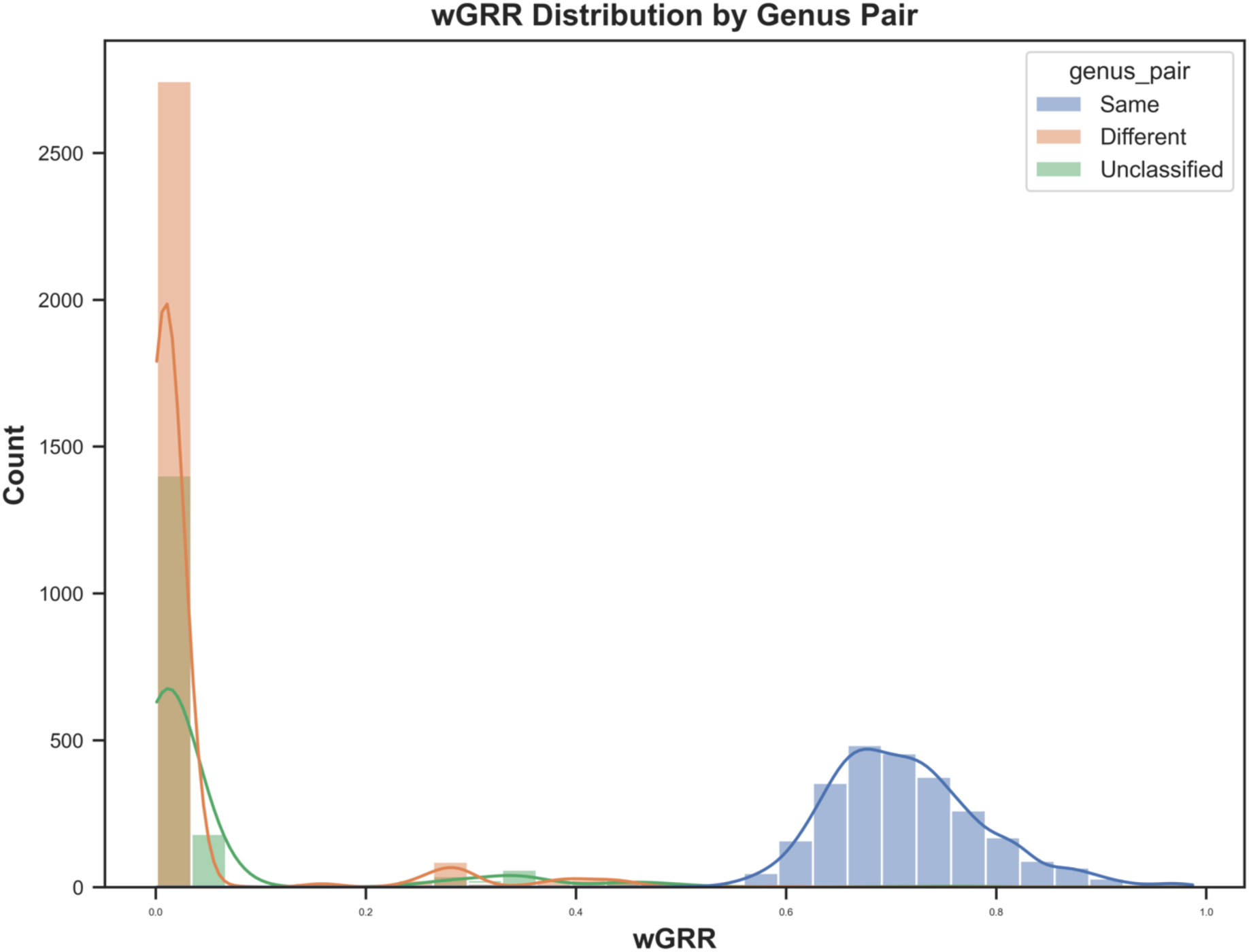
Distribution of pairwise weighted Gene Repertoire Relatedness (wGRR) scores across phage genomes. Scores reflect total protein content comparisons, color-coded by whether genome pairs belong to the same (solid) or different (dashed) genera. The dotted line at wGRR = 0.65 marks the threshold for defining distinct phage clusters.

**Figure S3:**
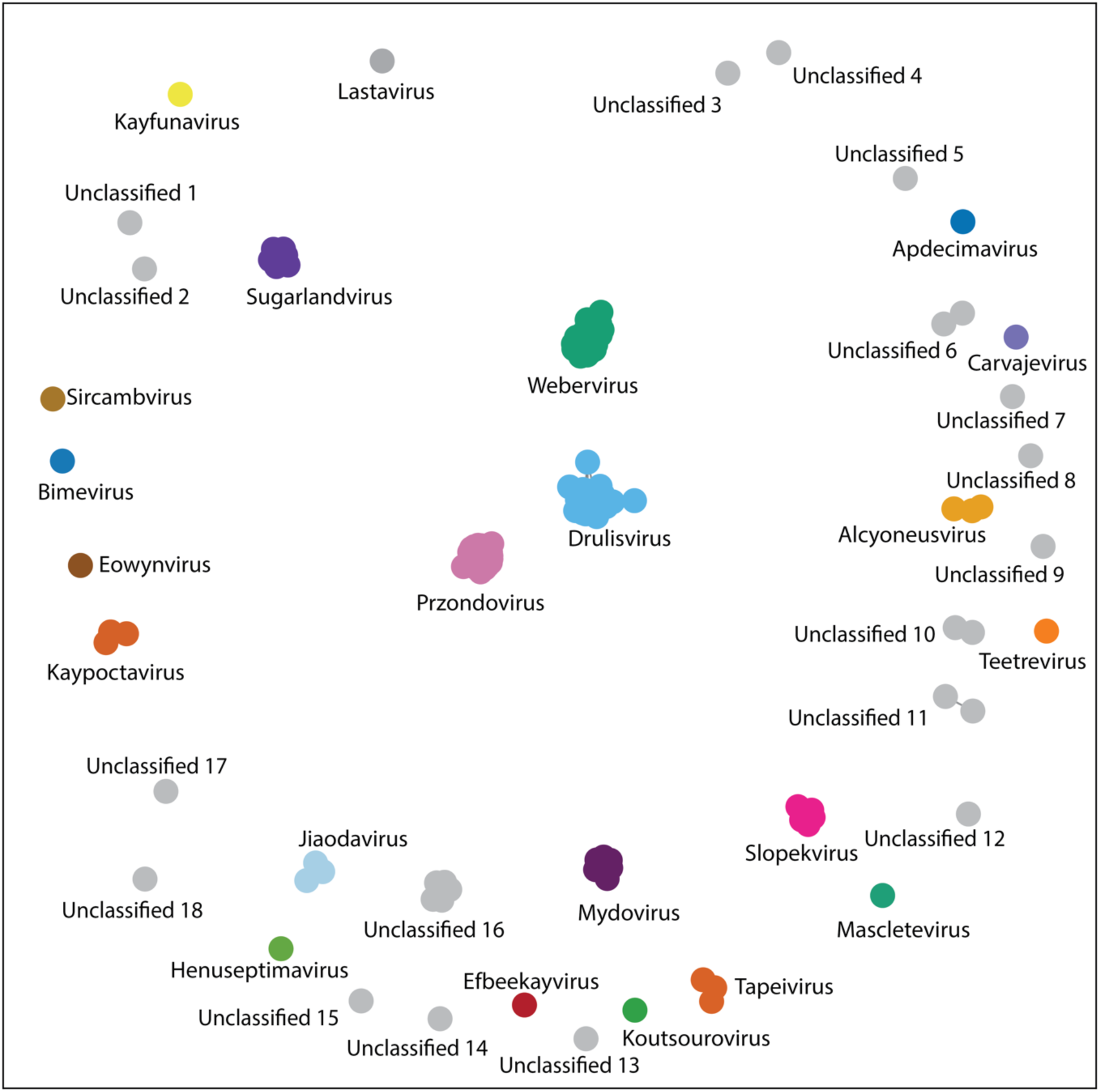
Phage genome clustering by MCL (wGRR = 0.65). This analysis produced 40 clusters, with 22 aligning to known genera and 18 indicating novel genera.

**Figure S4:**
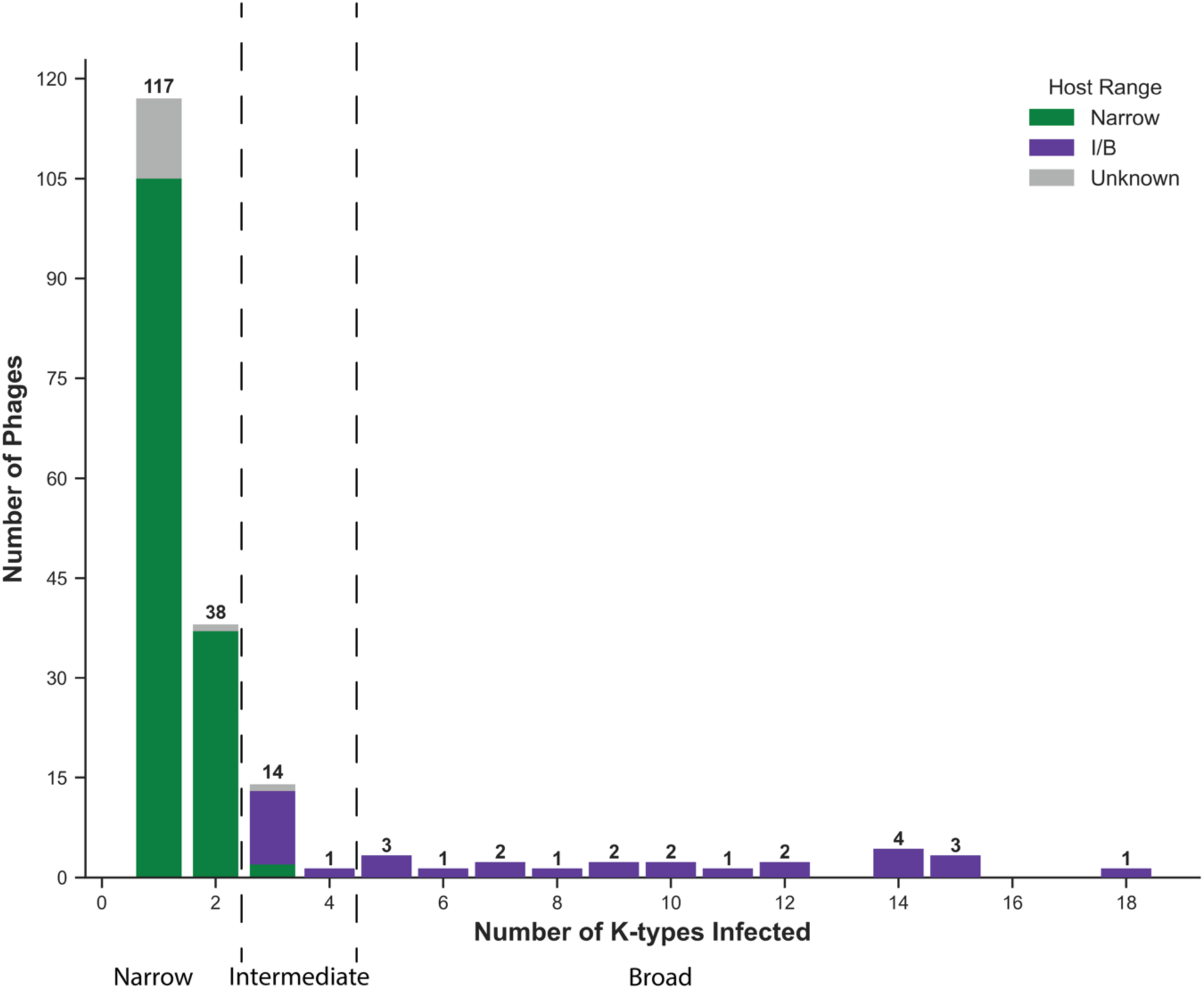
Distribution of phage host-range breadth. The plot shows the number of phages infecting 1–2 K-types (grouped as Narrow empirical host-range), 3–4 K-types (Intermediate, where the peak drops sharply), and 5+ K-types (Broad empirical host-range). The Intermediate category is not biologically defined — it serves as a buffer between Narrow and Broad, and may include some true broad-range phages due to limited testing panels, or rarely narrow-range phages if the isolation host K-type differs from those used for host-range testing.

**Figure S5:**
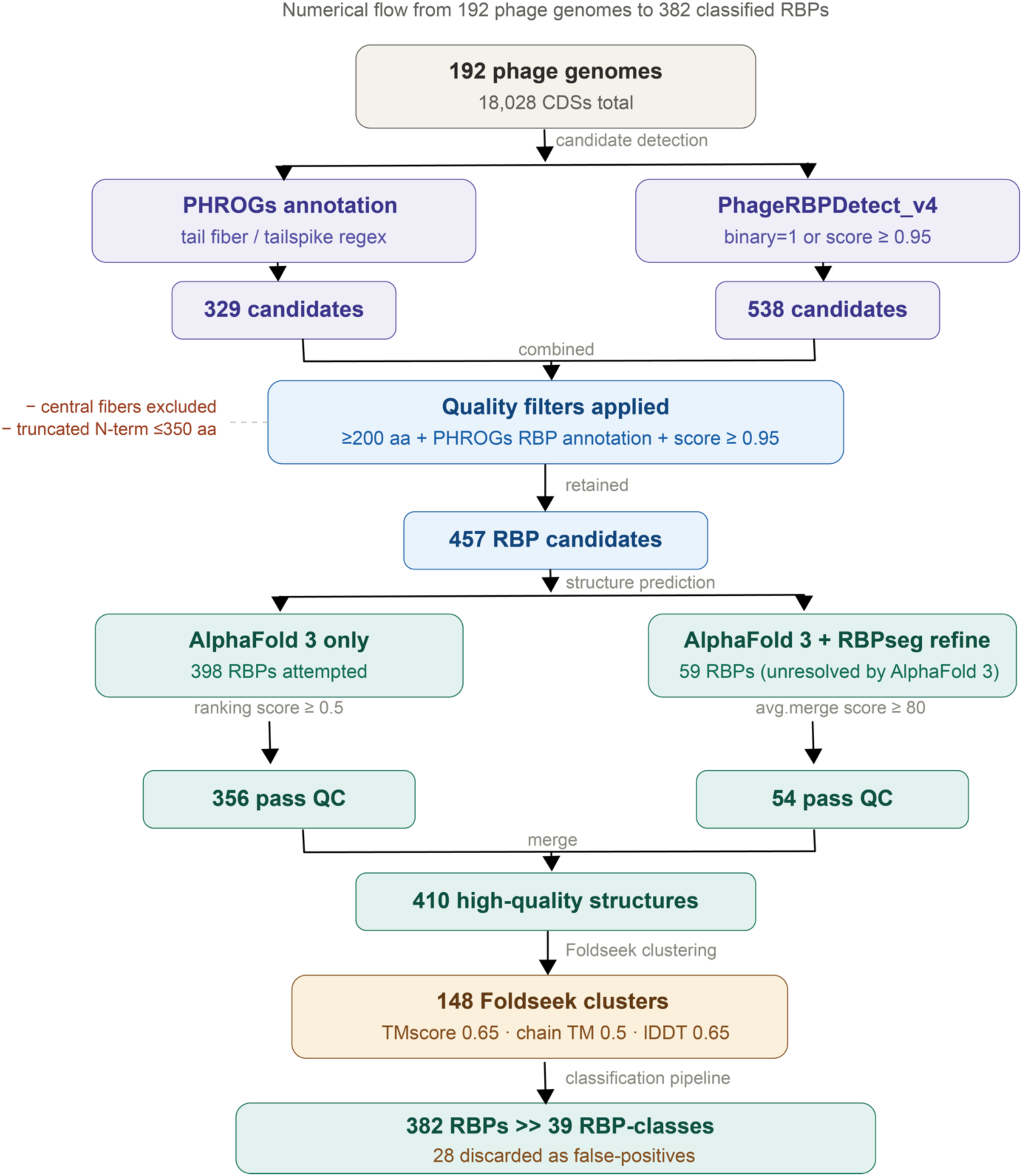
Pipeline for RBP detection, structure prediction, quality filtering, and classification. Numerical flow from 192 phage genomes (18,028 CDSs total) to 382 classified RBPs across 39 RBP-classes. RBP candidates were identified in parallel using PHROGs annotations (tail fiber/tailspike regex; 329 candidates) and PhageRBPDetect_v4 (binary prediction = 1 or normalized score ≥ 0.95; 538 candidates). The combined candidate set was filtered to retain only proteins ≥ 200 aa with both a PHROGs RBP annotation and a PhageRBPDetect_v4 score ≥ 0.95; central fibers and truncated N-terminal fragments ≤ 350 aa were additionally excluded, yielding 457 high-confidence RBP candidates. Homo-trimeric structures were predicted using AlphaFold 3 for all 457 candidates; 59 proteins unresolved by AlphaFold 3 alone were further refined using RBPseg. Quality filtering retained structures with an AlphaFold 3 ranking score ≥ 0.5 (356 of 398) or an average RBPseg merge score ≥ 80 (54 of 59), yielding 410 high-quality structures. These were clustered into 148 Foldseek clusters using Foldseek easy-multimercluster (multimer TMscore ≥ 0.65, chain TMscore ≥ 0.5, interface lDDT ≥ 0.65). A sequential classification pipeline assigned 382 RBPs to 39 RBP-classes; 28 proteins were discarded as likely false positives based on HMM-based functional annotation.

**Figure S6:**
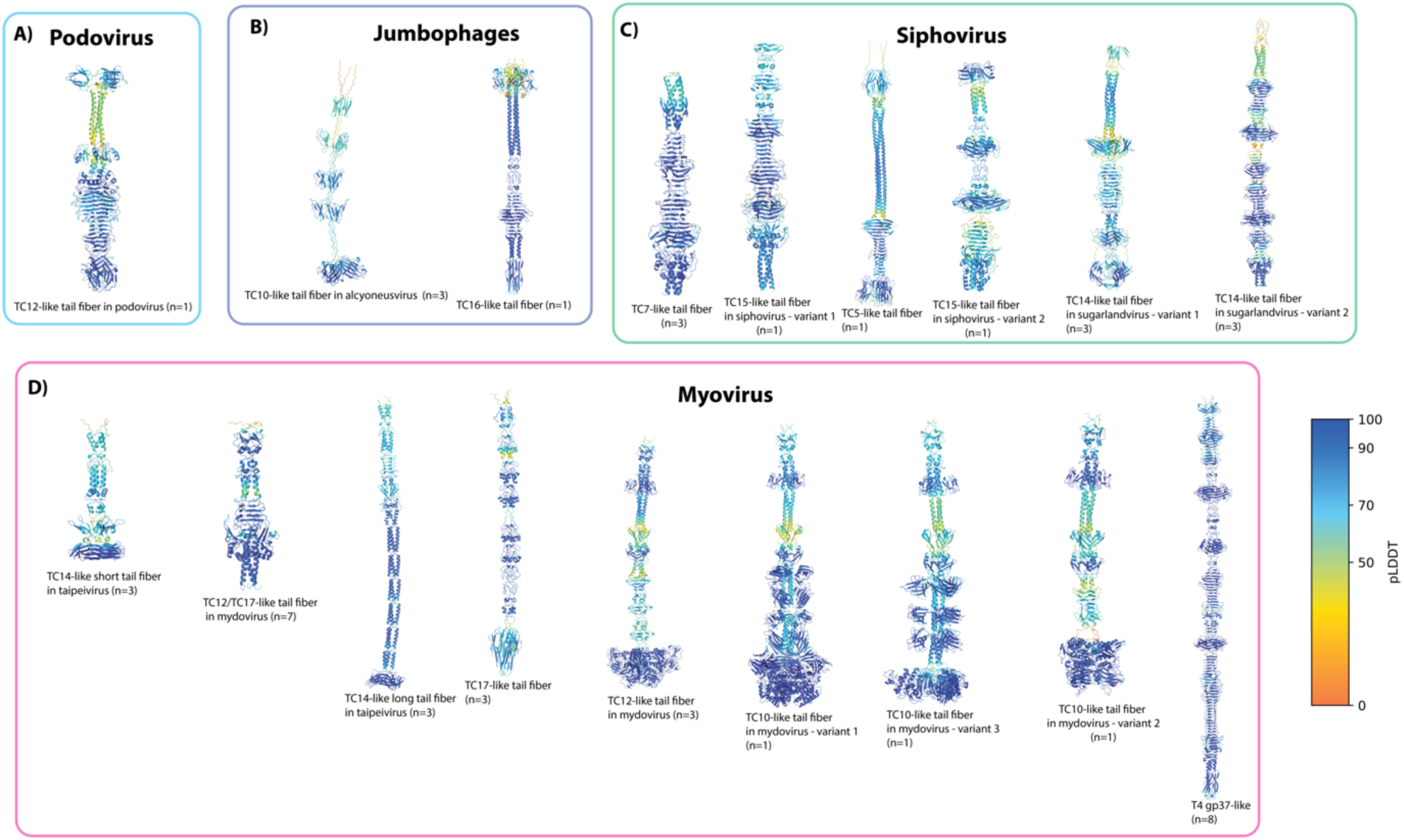
Structural models of the 18 structurally novel RBP-classes, organised by morphotype. Structures are coloured according to the AlphaFold per-residue confidence metric (pLDDT; 0–100 scale), with dark blue indicating high-confidence regions. RBP counts per class are shown in parentheses. (A) One structurally novel RBP-class exclusive to podoviruses. (B) Two structurally novel RBP-classes exclusive to jumbophages. (C) Six structurally novel RBP-classes exclusive to siphoviruses. (D) Nine structurally novel RBP-classes exclusive to myoviruses, including T4 gp37-like, which was retained among the structurally novel due to the absence of literature precedent in *Klebsiella* phages; its function is indirectly inferred from co-occurrence with T4 gp34-like, gp36-like, and gp12-like proteins in the same phage genomes, supporting membership in a T4-like tail fiber system. All protein structures are shown and described in the Klebsiella RBP Atlas Text S1.

**Figure S7:**
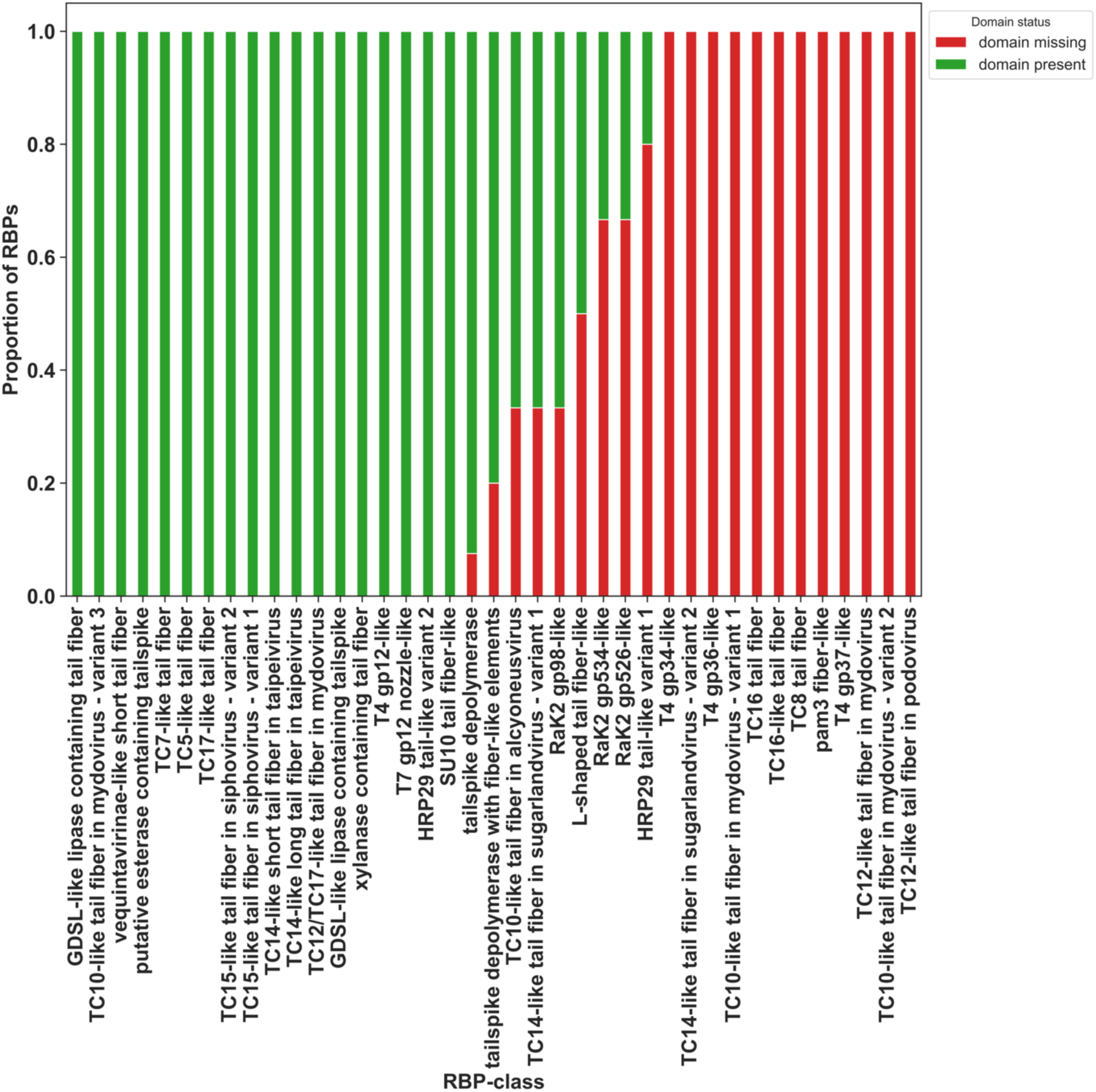
ECOD domain mapping across 39 RBP-classes. RBPs are scored for the presence (at least one confidently mapped domain) or absence of ECOD domains.

**Figure S8:**
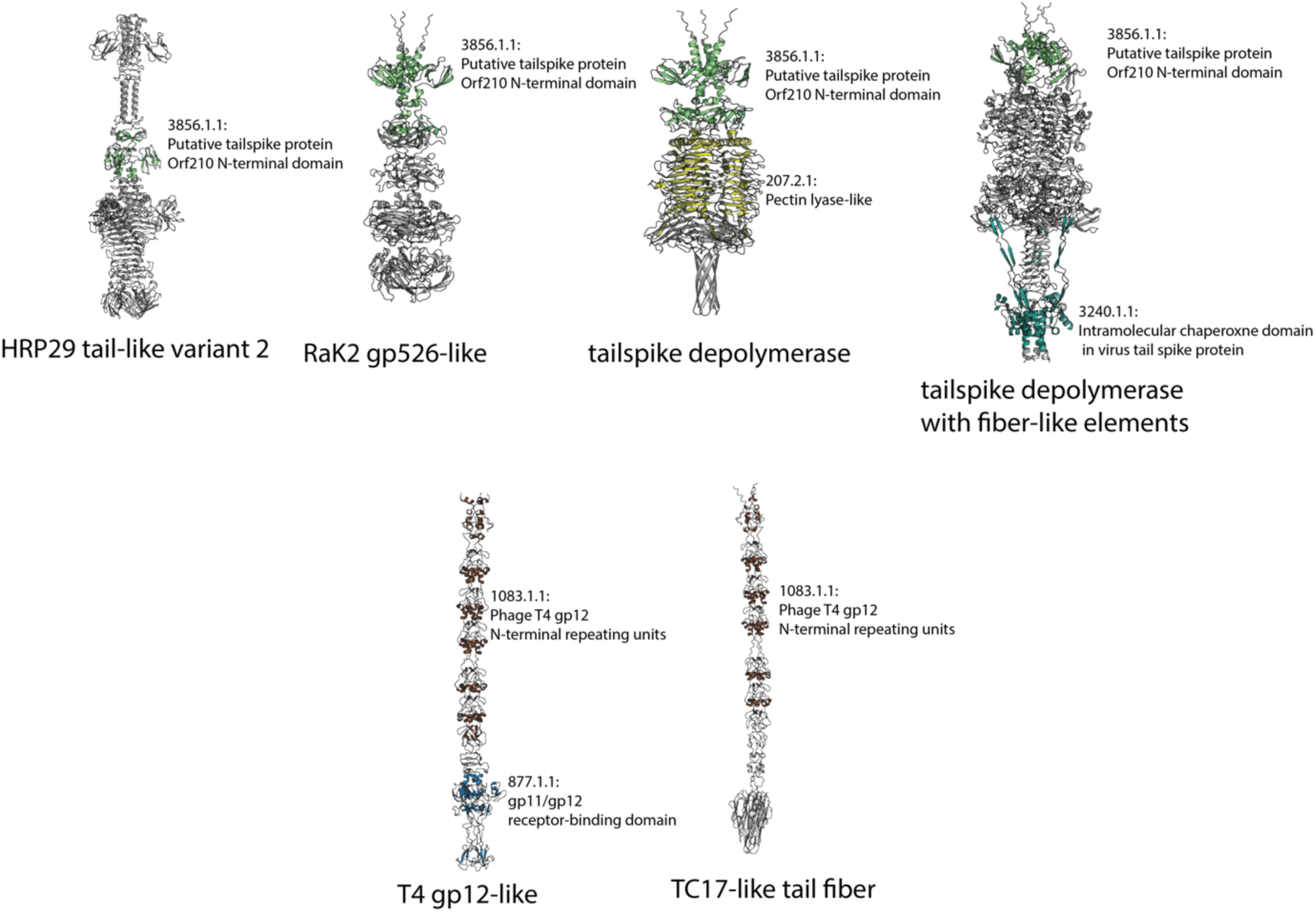
Domain architectures of representative RBPs. The diagram illustrates RBP-classes grouped by their shared N-terminal ECOD domains. Representatives sharing the putative tailspike protein Orf210 N-terminal domain (3856.1.1) are shown in the top row, while those sharing the phage T4 gp12 N-terminal repeating units (1083.1.1) are displayed in the bottom row.

**Figure S9:**
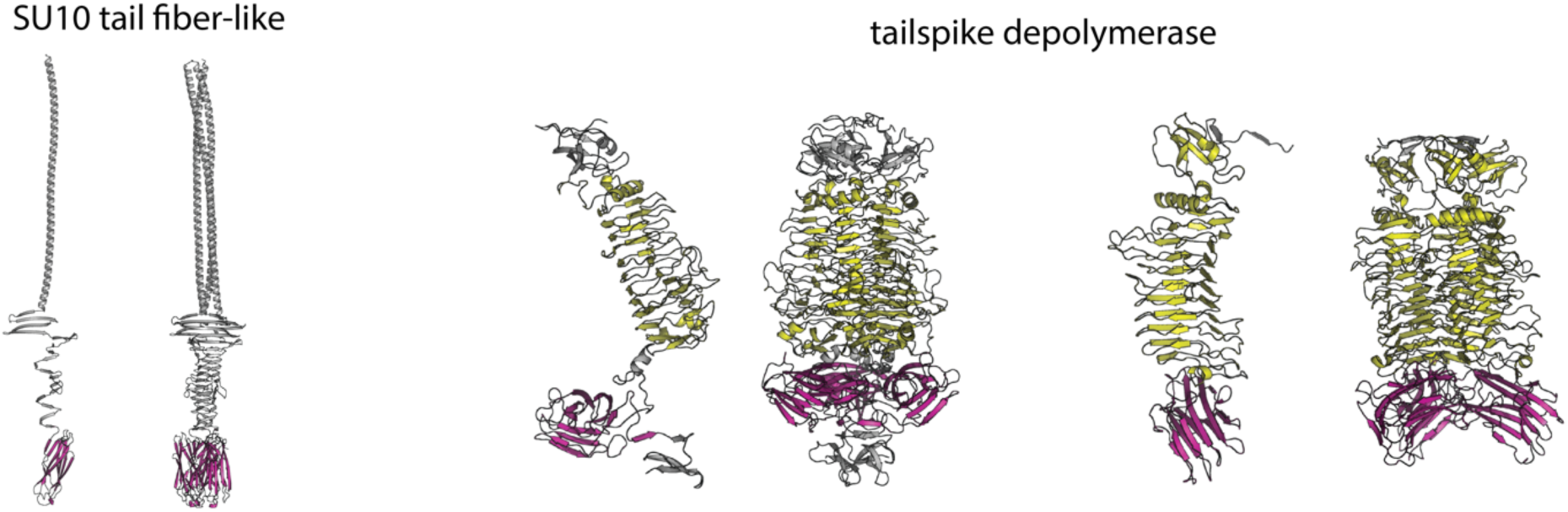
Conformational variation of the Galactose-binding domain (10.1.2) across oligomeric states. The structures illustrate differences between monomeric (left) and trimeric (right) forms from two distinct RBP-classes. The 10.1.2 domain is highlighted in pink, and the Pectin lyase-like domain (207.2.1) is shown in yellow.

**Figure S10:**
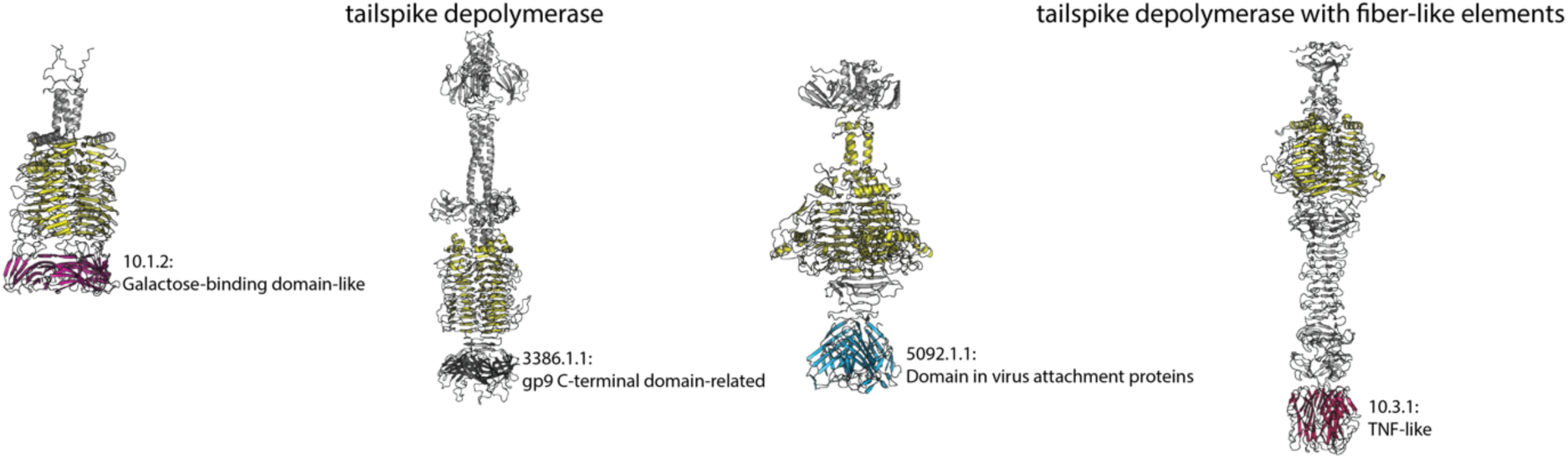
Structural diversity of C-terminal domains in depolymerase-containing RBPs. Representative structures highlight topological variations among tailspike depolymerases (first three from left) and a tailspike depolymerase with fiber-like elements (far right). The Pectin lyase-like ECOD domain (207.2.1) is colored yellow across all models.

**Figure S11:**
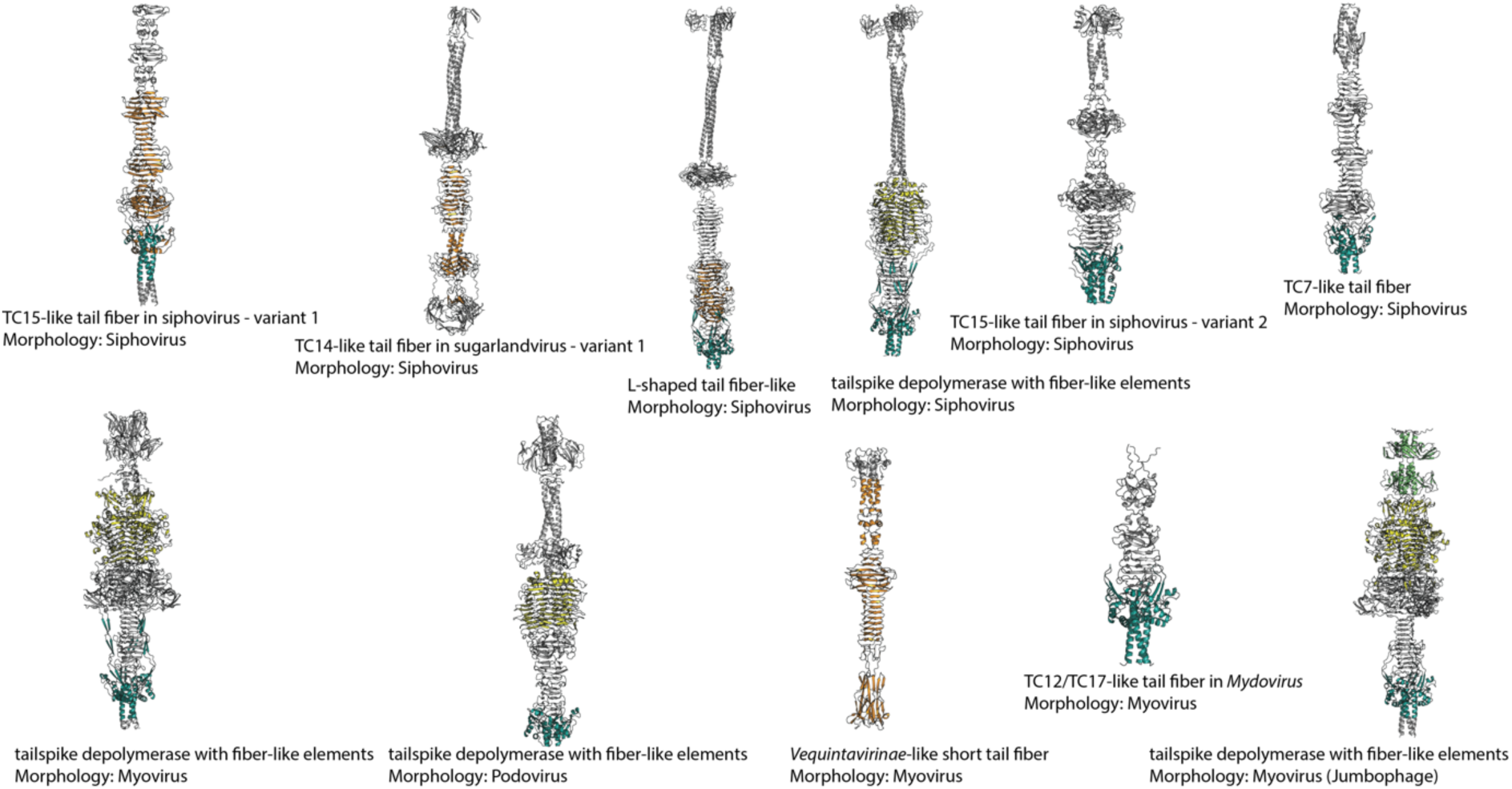
Distribution of scaffold ECOD domains across RBP-classes and phage morphotypes. The conserved IMC (3240.1.1) and tail fiber trimerization (79.1.1) domains are found across diverse structural contexts. Mapped domains include: 3240.1.1 (bluish-green), 79.1.1 (orange), the Pectin lyase-like domain (207.2.1, yellow), and the putative tailspike Orf210 N-terminal domain (3856.1.1, light green).

**Figure S12:**
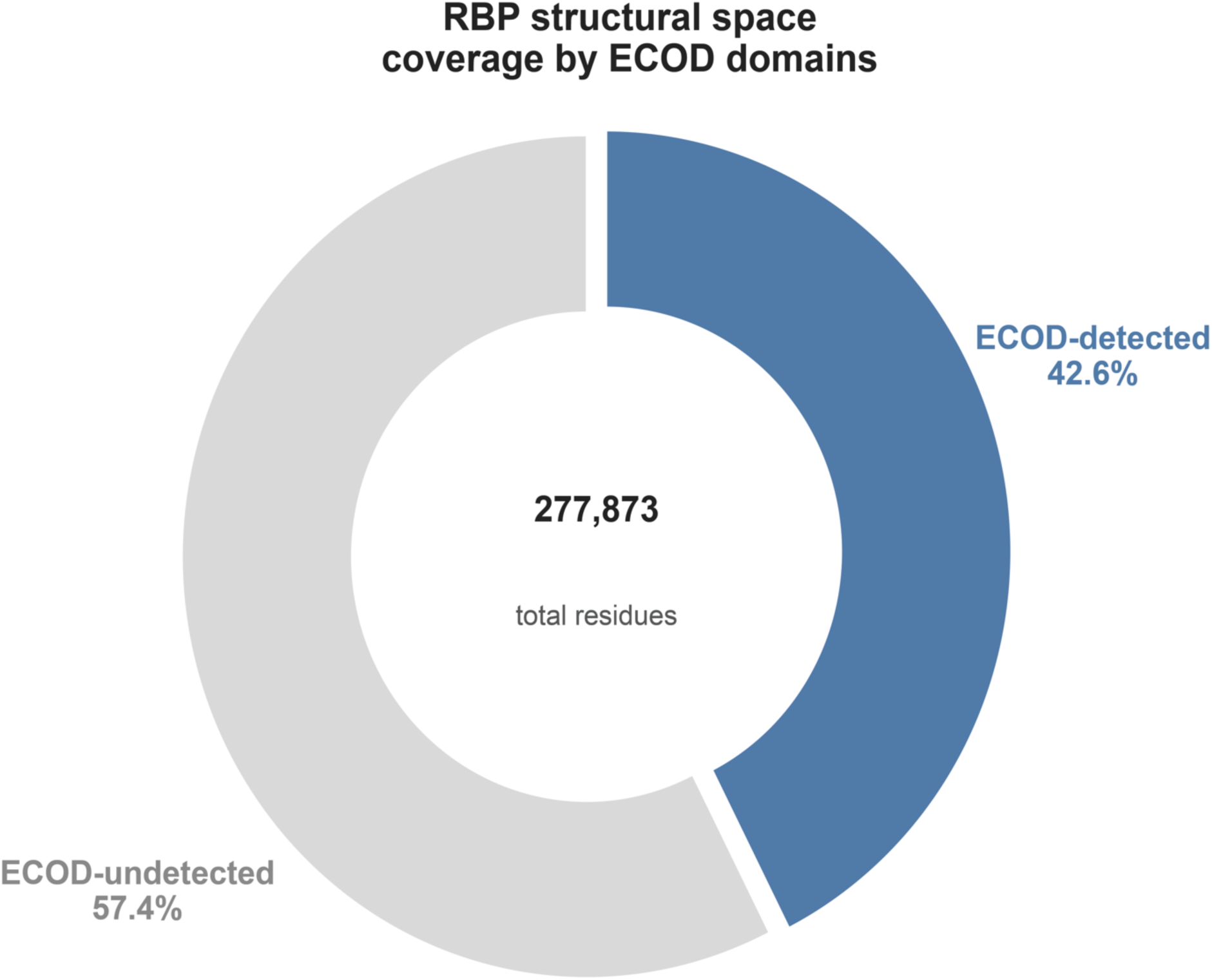
ECOD domain coverage of the RBP structural space (calculated based on monomeric chain length). Donut chart showing the proportion of total amino-acid residues across all 382 confirmed RBPs that are covered by at least one ECOD domain hit (blue, 42.6%) versus residues residing in structural regions with no detectable ECOD homology (grey, 57.4%). Coverage per protein was computed as the union of all ECOD domain hit intervals to avoid double-counting overlapping hits. Total residues analysed: 277,873.

**Figure S13:**
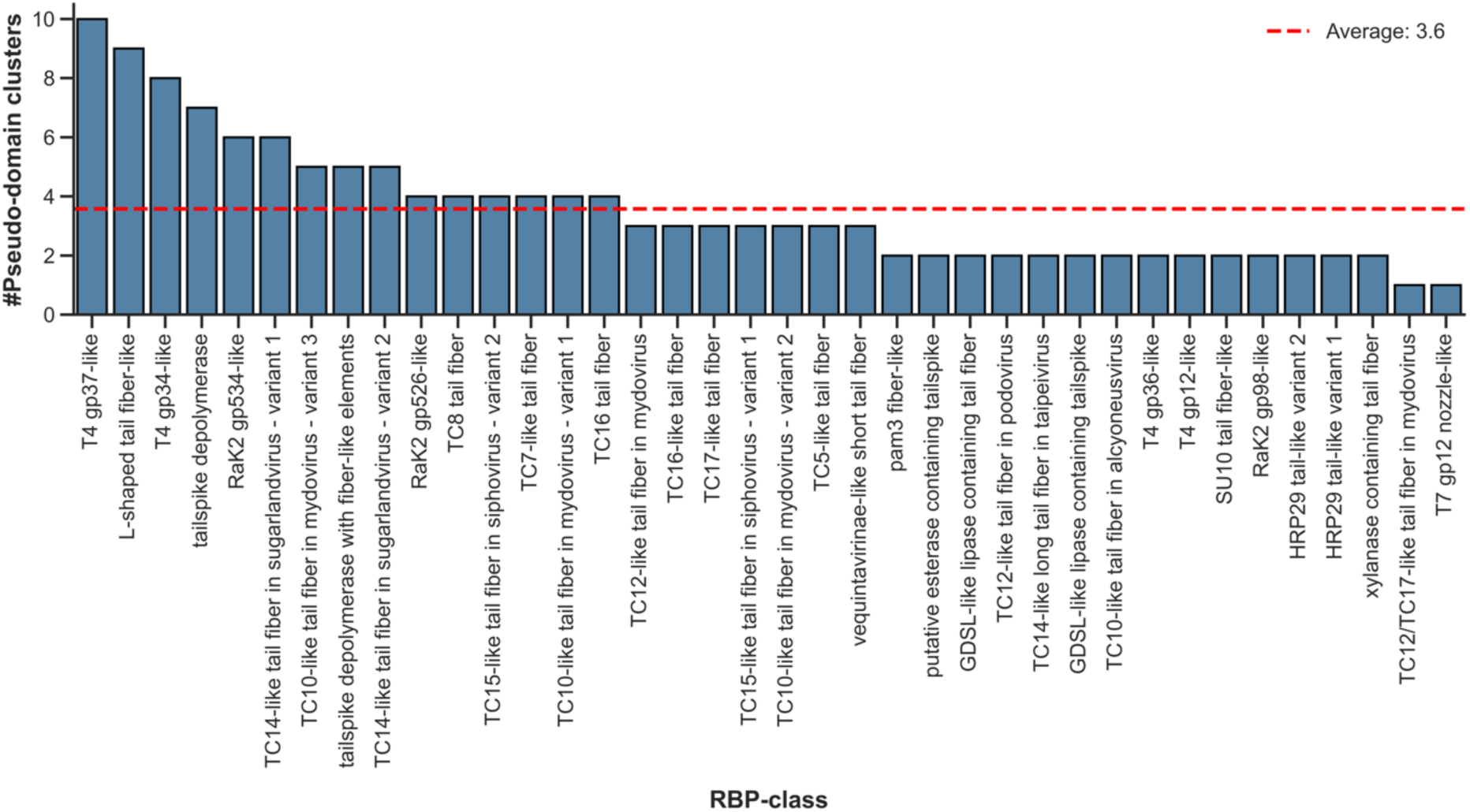
Distribution of pseudo-domain clusters across RBP classes. Pseudo-domains were detected and mapped in 38 of the 39 analyzed RBP-classes. The red dashed line indicates the average number of pseudo-domain clusters per RBP-class.

**Figure S14:**
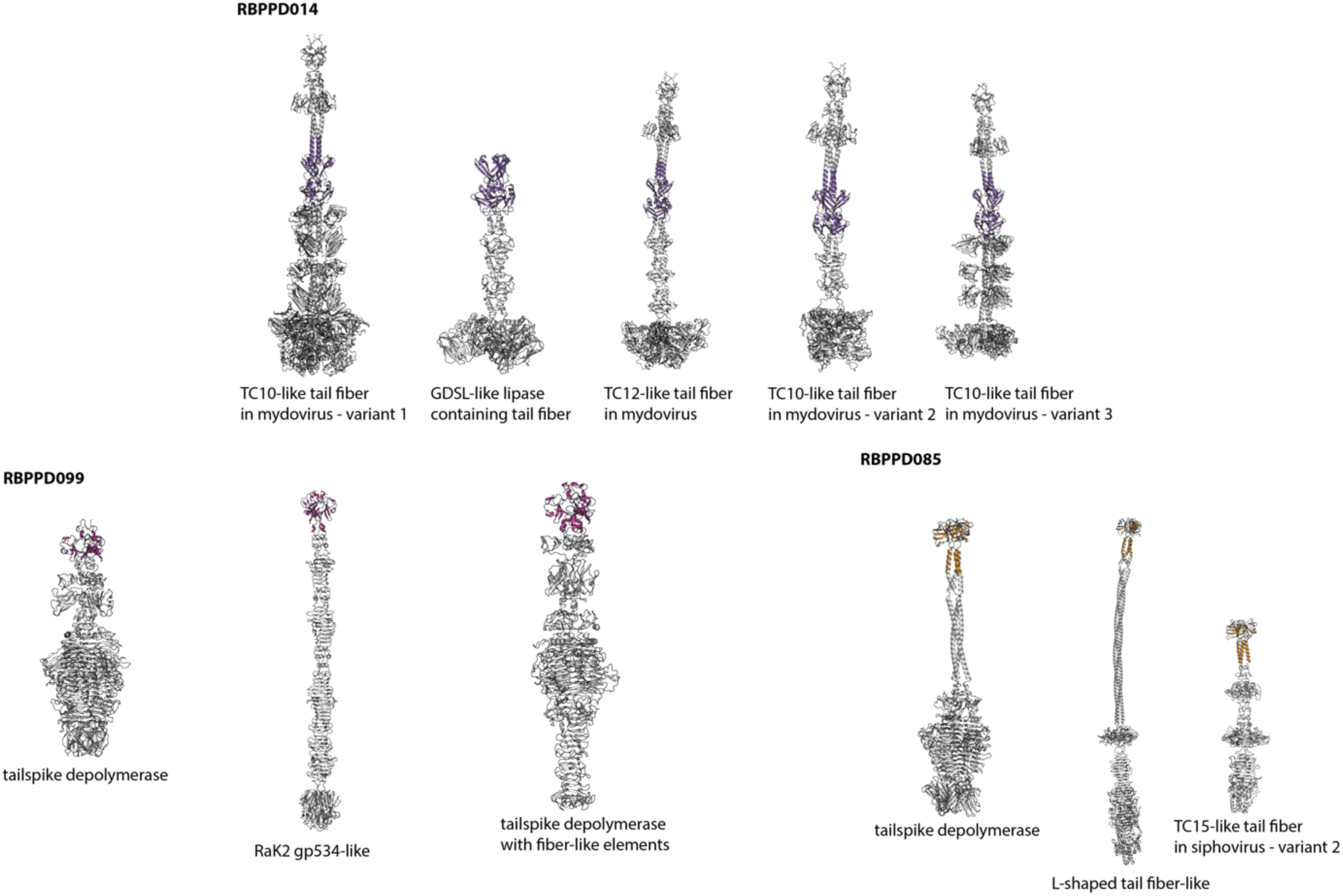
Predominant pseudo-domain clusters shared across multiple RBP-classes (located at N-terminal ends). Representative examples highlight clusters present in more than one RBP-class. Each distinct pseudo-domain cluster is mapped onto a representative RBP structure using a unique color.

**Figure S15:**
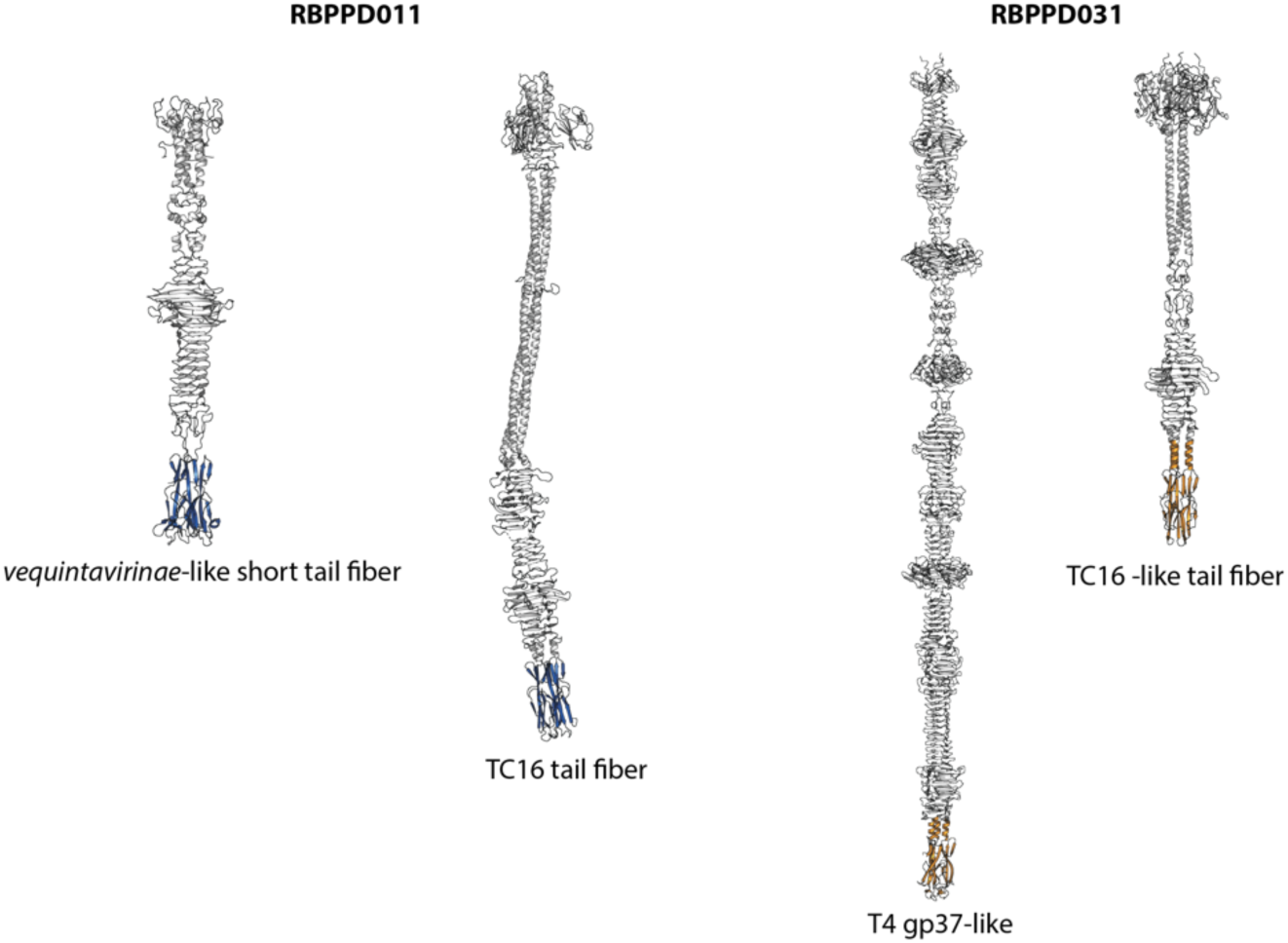
Mapping of C-terminal pseudo-domain clusters across distinct RBP-classes. Two representative pseudo-domain clusters are mapped to the C-terminal regions of various RBP structures. These clusters overlap with potential carbohydrate-binding domains but do not correspond to any currently defined ECOD domains.

**Figure S16:**
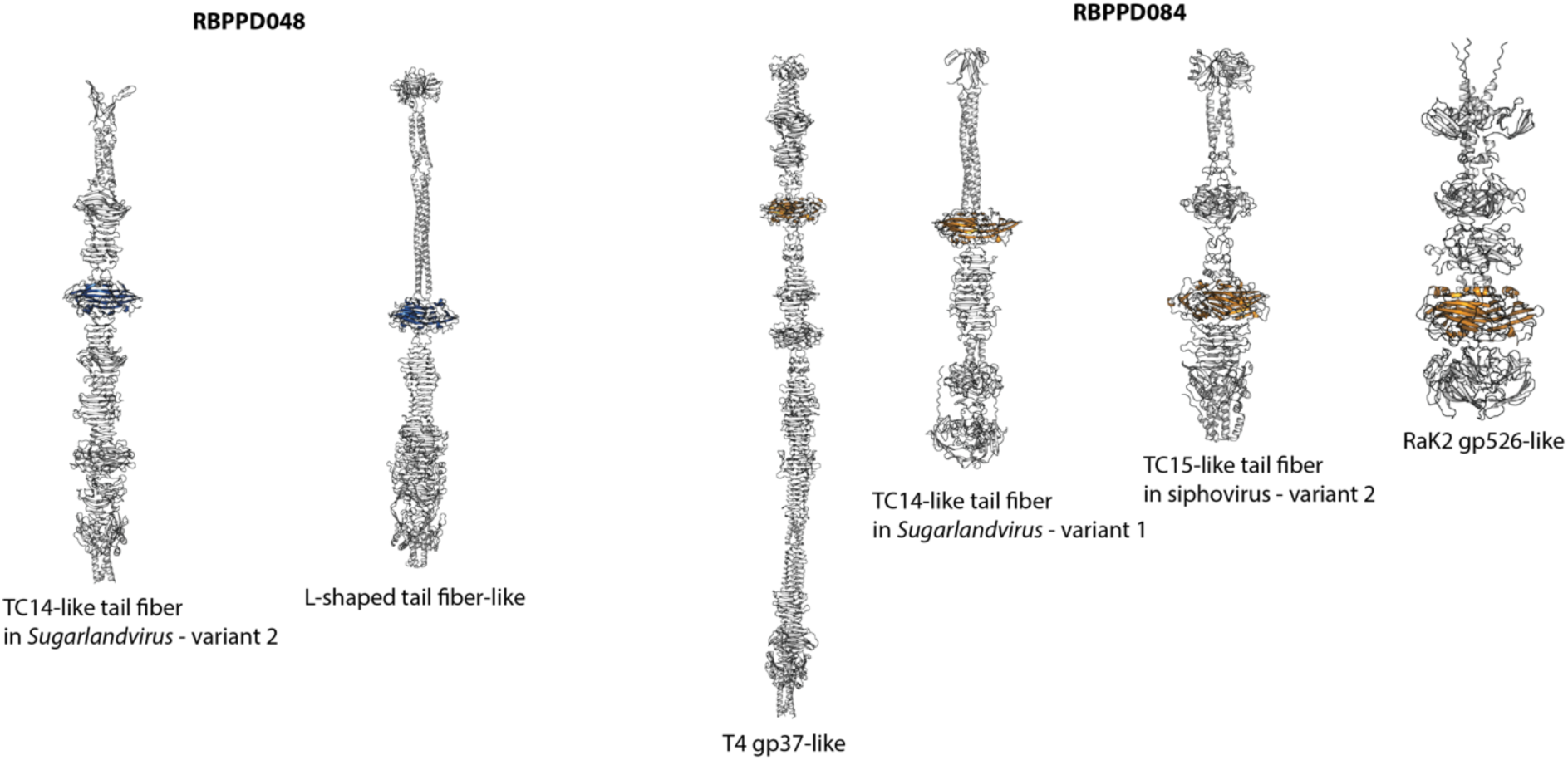
Sensitive detection of distinct pseudo-domain clusters in structurally similar knob regions. Pseudo-domain clusters mapped to the knob regions of members from different RBP-classes. Notably, despite the high structural similarity between these two knob regions, they are classified into distinct pseudo-domain clusters. This distinction highlights the sensitivity and resolution of the clustering approach.

**Figure S17.**
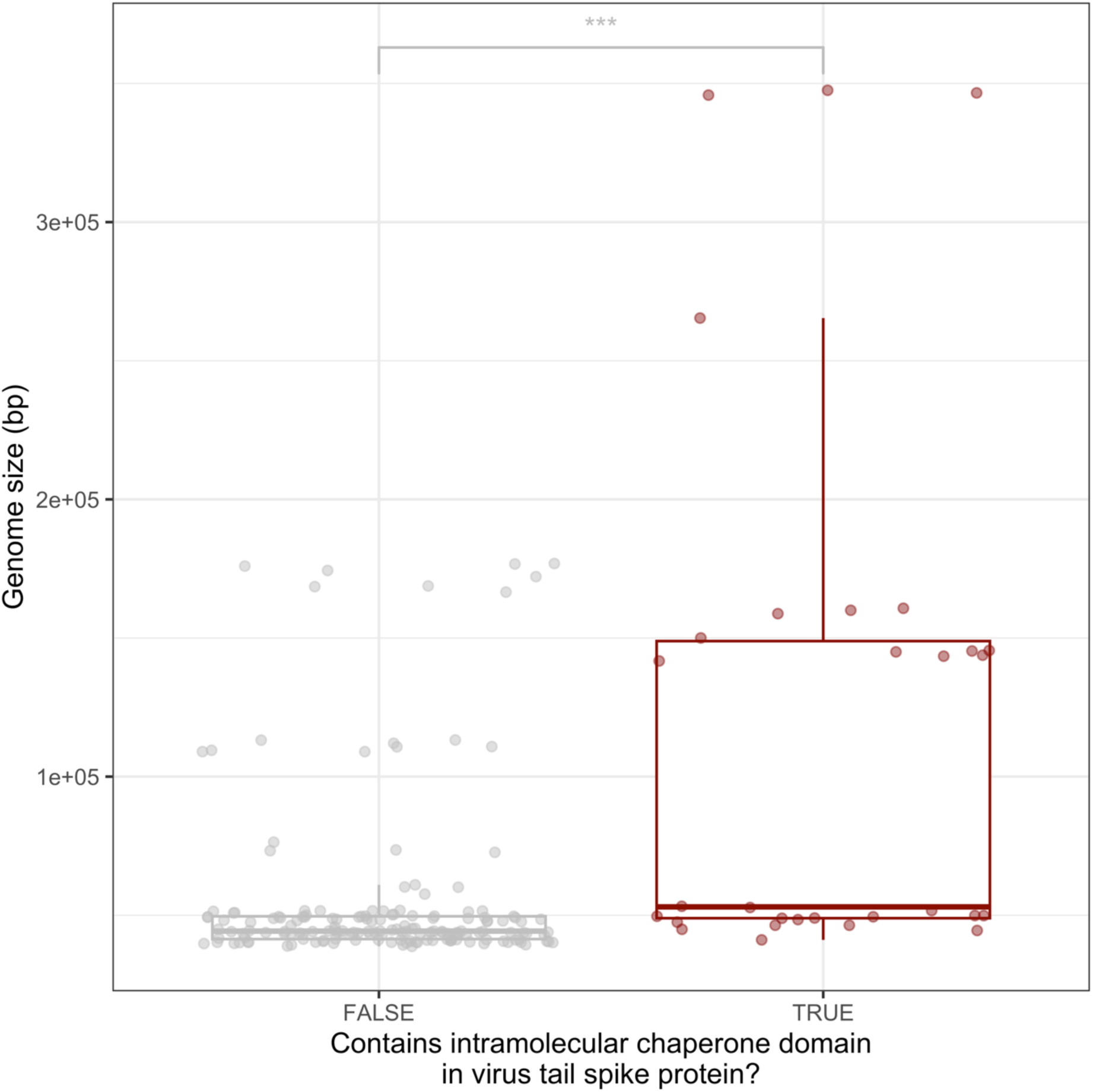
Phage genome size within phages that contain at least one Intramolecular chaperone domain in any of their tail spike proteins. The difference of genome size is significant between the two groups according to a two-sided Wilcoxon test (p.value < 0.001).

**Figure S18:**
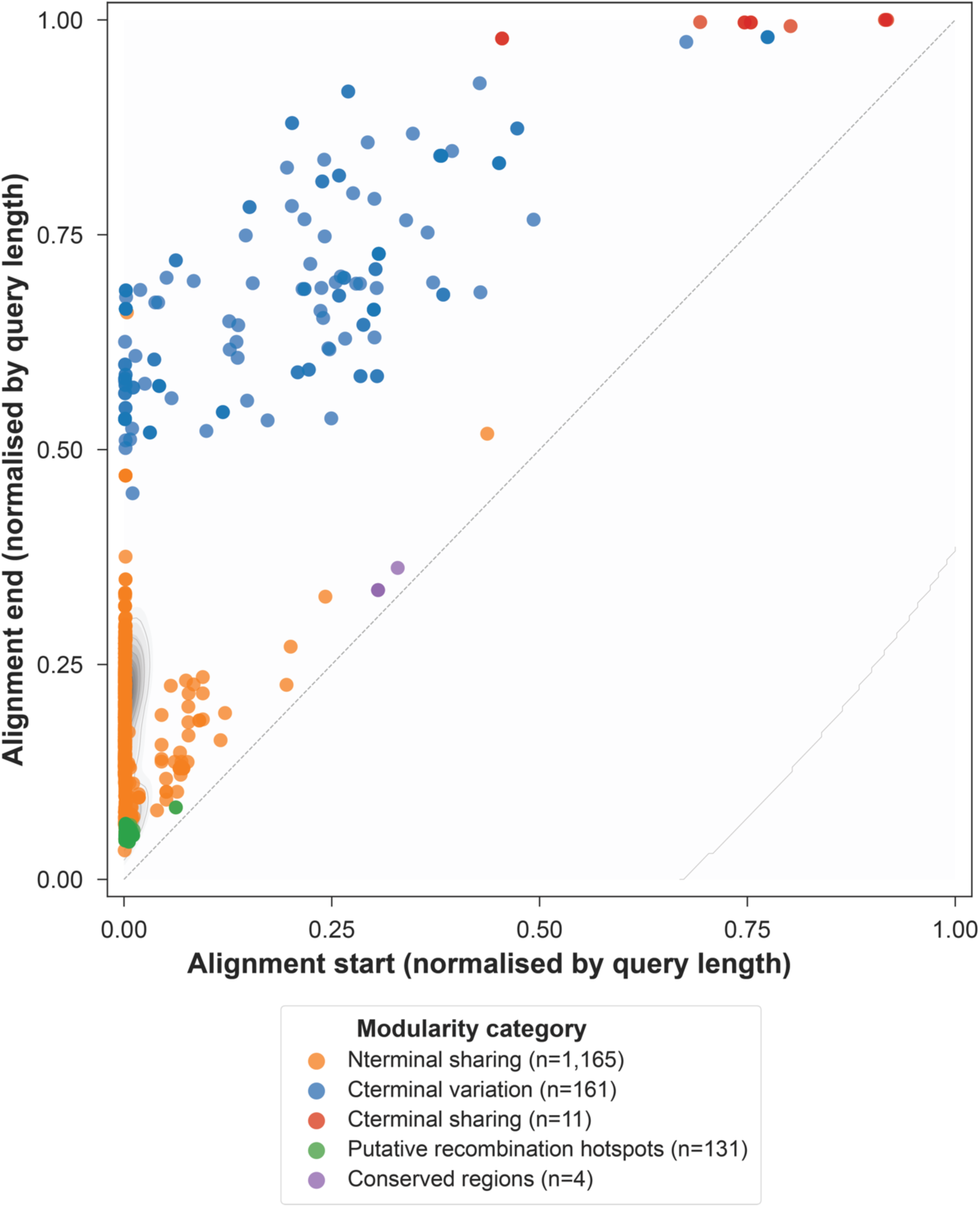
Empirical distribution of BLASTP hits of RBPs across the normalised protein sequence space. Each point represents one pairwise BLASTP hit plotted at its alignment start and end position, both divided by query length. Background contours show the kernel density estimate of all hits; coloured points overlay the five modularity categories. N-terminal sharing (orange) and C-terminal variation (blue) dominate the signal and occupy distinct, non-overlapping regions of the sequence space, consistent with modular exchange of functional domains. C-terminal sharing (red) co-localises with C-terminal variation but represents high-coverage, high-identity hits between RBPs of different RBP-classes. Putative recombination hotspots (green) cluster at intermediate normalised positions, marking internal exchange boundaries. Conserved regions (purple) fall near the diagonal, reflecting near-full-length similarity. Together, the five categories account for the large majority of observed BLASTP signals, validating the modularity classification scheme.

**Figure S19:**
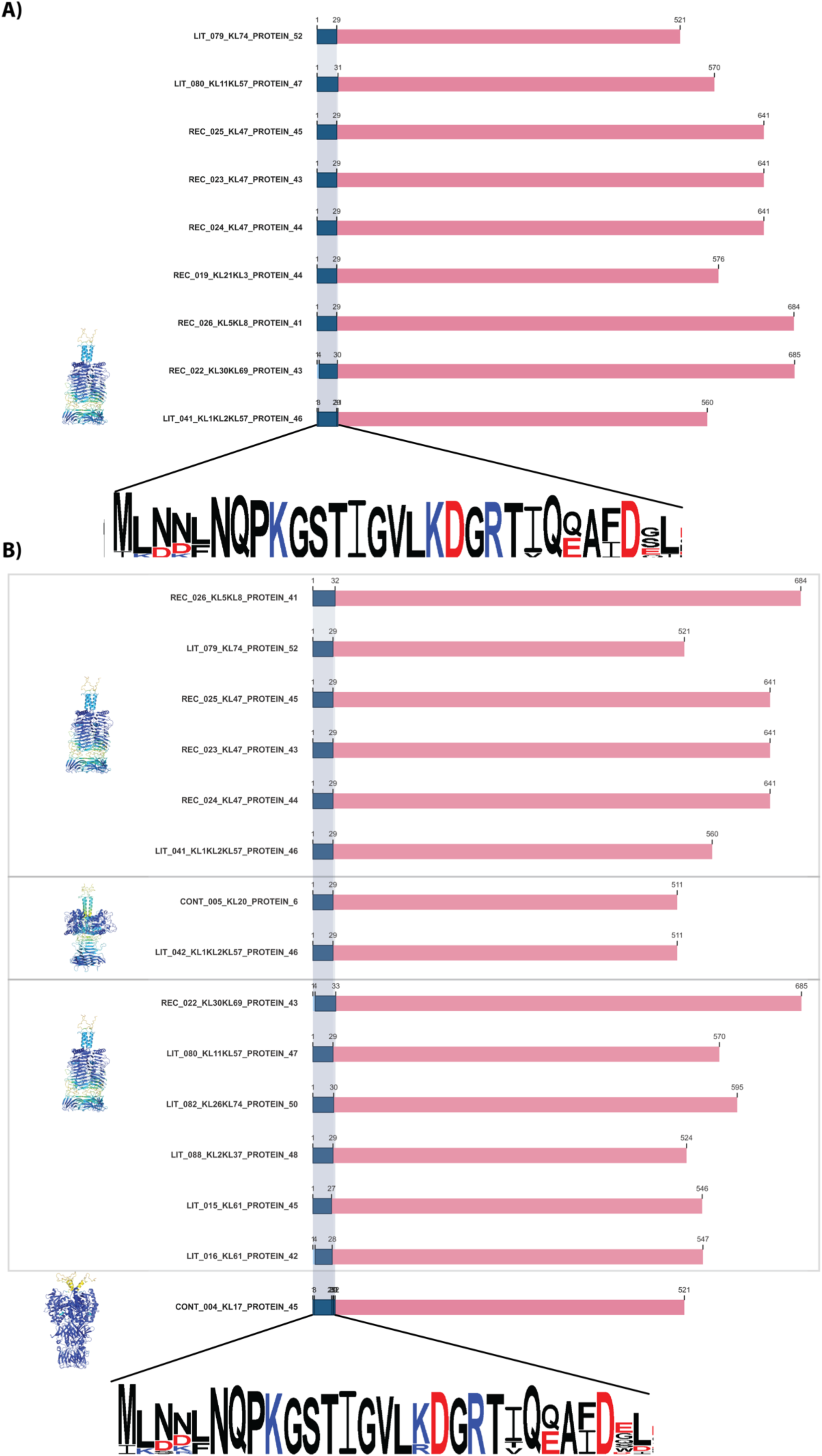
Mapping of recombination hotspots across RBP-classes at the amino acid level. These hotspots are approximately 30 amino acids in length and exhibit high sequence similarity regardless of class boundaries. (A) Recombination hotspots shared between members of the same RBP class (tailspike depolymerases). (B) Hotspots conserved between different RBP classes, showing alignments of a putative esterase-containing tailspike query against tailspike depolymerase and GDSL-like lipase-containing tailspike targets. Consensus sequence logos were generated with WebLogo 3 (https://weblogo.threeplusone.com/create.cgi).

**Figure S20:**
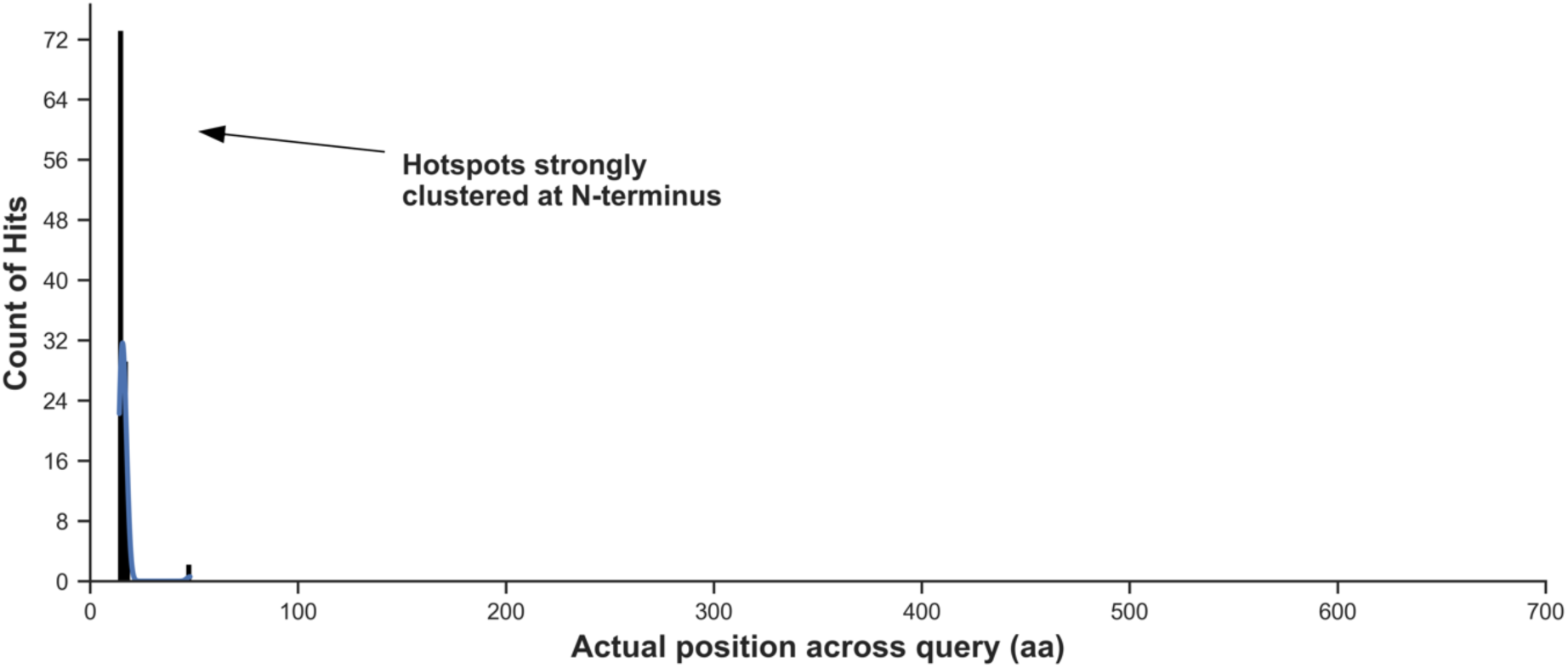
Recombination hotspots predominantly localize to the N-terminal region of the protein, regardless of RBP-class.

**Figure S21:**
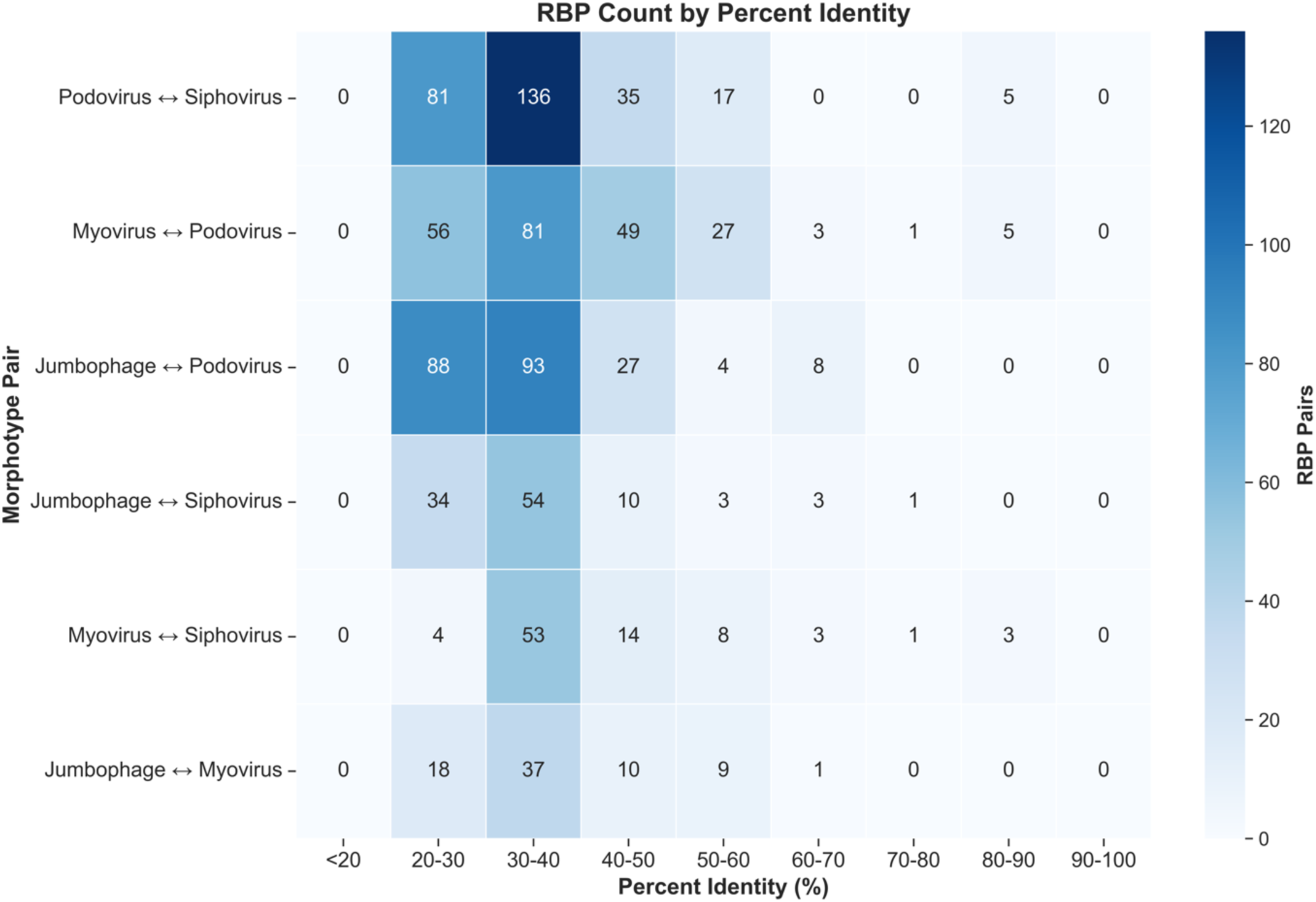
Heatmap of cross-morphotype sequence similarity for RBPs with structural depolymerase activity. Each cell shows the number of inter-morphotype pairs within a percent identity bin across all six morphotype combinations. The analysis includes 982 pairs in total, of which 876 exhibit low sequence similarity (<50%) and 106 exhibit higher sequence similarity (≥50%). Pairs are distributed across all six morphotype combinations, with Podovirus ↔ Siphovirus (274 pairs), Myovirus ↔ Podovirus (222 pairs), and Jumbophage ↔ Podovirus (220 pairs) contributing the most. Only same-class RBP pairs (i.e., query and target sharing the same RBP class) with e-value ≤ 0.001 are included.

**Figure S22:**
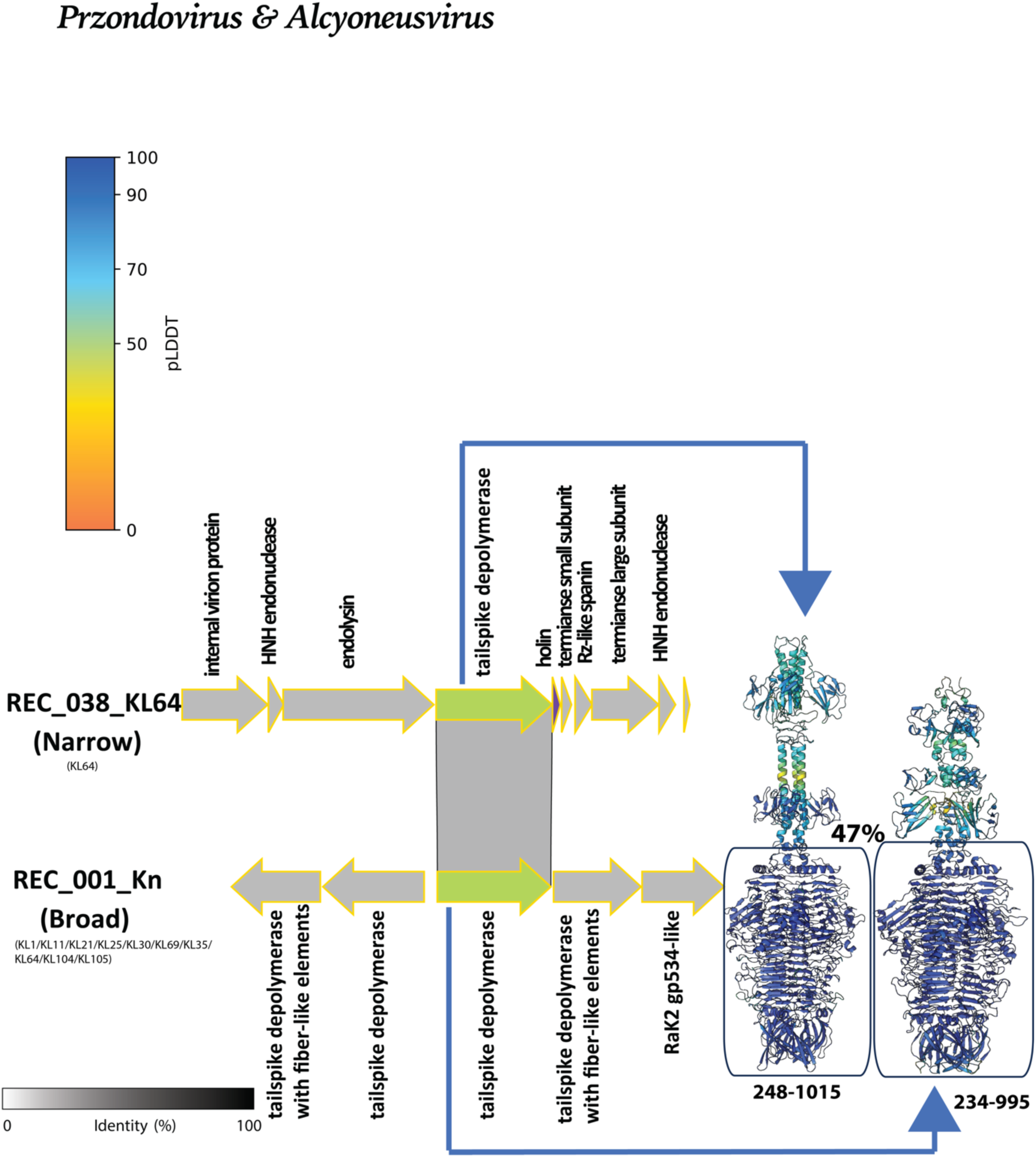
Cross-morphotype depolymerase sharing between *Przondovirus* and *Alcyoneusvirus* indicates ancient modular exchange with conserved capsule specificity. Gene neighborhoods of a *Przondovirus* podovirus and an *Alcyoneusvirus* jumbophage (drawn to scale; arrows indicate ORFs and transcriptional direction) with pairwise amino acid identity across the locus (grey-scale shading, 0–100% identity), alongside predicted RBP structures colored by residue confidence (pLDDT scale). Square boxes drawn between the two RBPs represent regions of amino acid sequence similarity. REC_038_KL64 (Narrow host range; KL64) encodes a single tailspike depolymerase flanked by lysis and accessory genes, while REC_001_Kn (Broad host range; KL1/KL11/KL21/KL25/KL30/KL35/KL64/KL69/KL104/KL105) encodes multiple RBPs including a tailspike depolymerase, a tailspike depolymerase with fiber-like elements, and a RaK2 gp534-like fiber. The shared tailspike depolymerase region (residues 248–1015 and 234–995, respectively) displays 47% amino acid identity, indicating conserved receptor-recognition function despite deep phylogenetic divergence across morphotypes.

**Figure S23:**
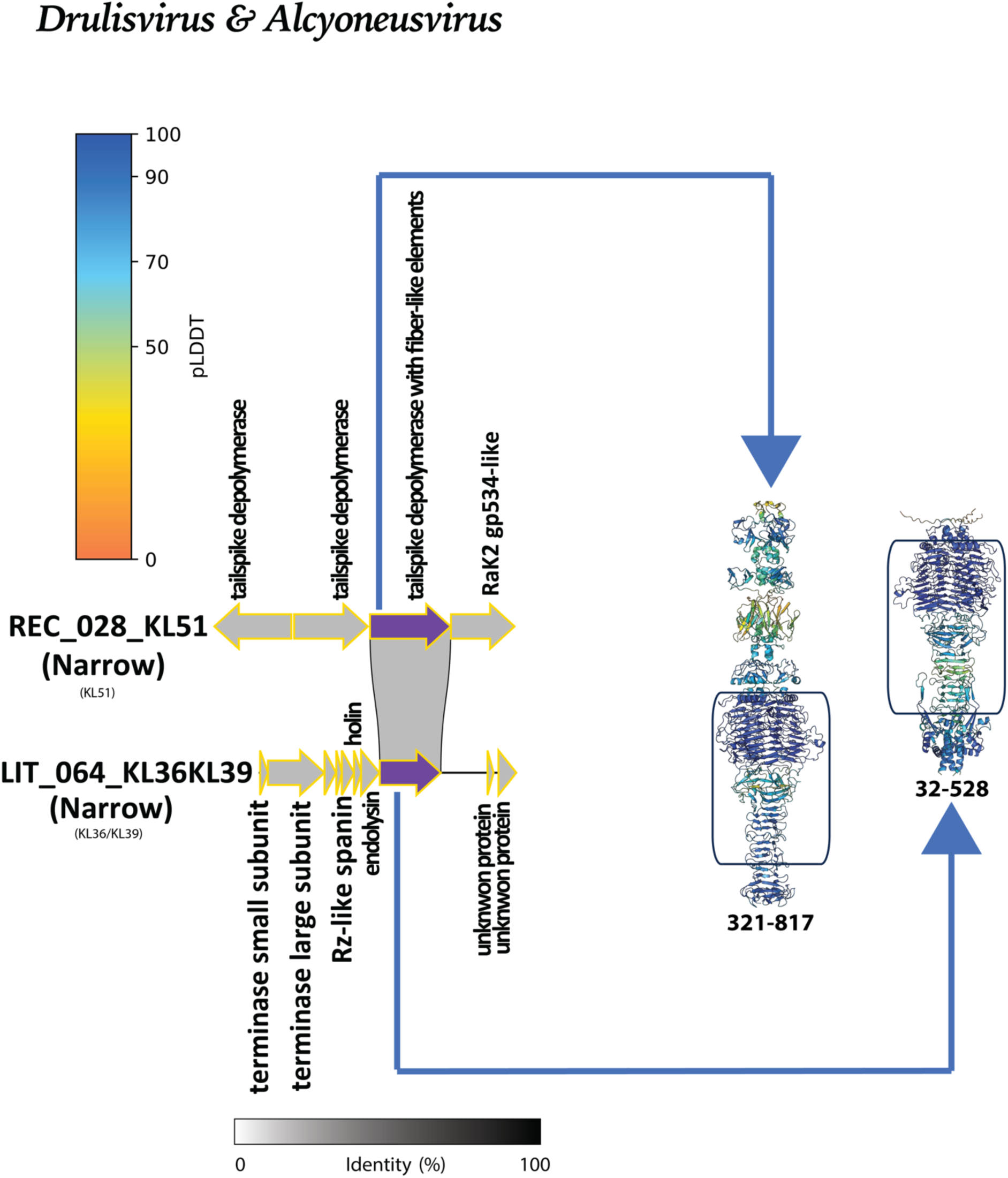
Cross-morphotype sharing of a tailspike depolymerase with fiber-like elements between *Drulisvirus* and *Alcyoneusvirus* suggests conserved capsule specificity across podovirus and jumbophage lineages. Gene neighborhoods of a *Drulisvirus* podovirus and an *Alcyoneusvirus* jumbophage (drawn to scale; arrows indicate ORFs and transcriptional direction) with pairwise amino acid identity across the locus (grey-scale shading, 0–100% identity), alongside predicted RBP structures colored by residue confidence (pLDDT scale). Square boxes drawn between the two RBPs represent approximate regions of amino acid sequence similarity; exact coordinates of the shared region are indicated adjacent to each structure. LIT_064_KL36KL39 (Narrow host range; KL36/KL39) encodes a minimal RBP locus with two depolymerases (tailspike depolymerase (not shown) and tailspike depolymerase with fiber-like elements) flanked by lysis genes, while REC_028_KL51 (Narrow host range; KL51) encodes a more complex locus comprising two tailspike depolymerases, a tailspike depolymerase with fiber-like elements, and a RaK2 gp534-like RBP. The shared region (residues 321–817 and 32–528, respectively) corresponds to the capsule-specific depolymerase body of the tailspike depolymerase with fiber-like elements, and notably excludes the intramolecular chaperone (IMC) domain present in the *Alcyoneusvirus* representative but absent from its *Drulisvirus* counterpart, indicating that the conserved module is restricted to the receptor-binding portion of the protein. Although REC_028_KL51 is classified as narrow host range based on KL51 specificity, this reflects a limited testing panel rather than genuine receptor restriction; the presence of a shared tailspike depolymerase with fiber-like elements suggests that this jumbophage likely harbours additional specificity towards KL36 or KL39, with individual depolymerases within its multi-RBP repertoire mediating distinct capsule-type recognition in a manner analogous to the two depolymerases conferring KL36 and KL39 specificity in the *Drulisvirus*.

**Figure S24:**
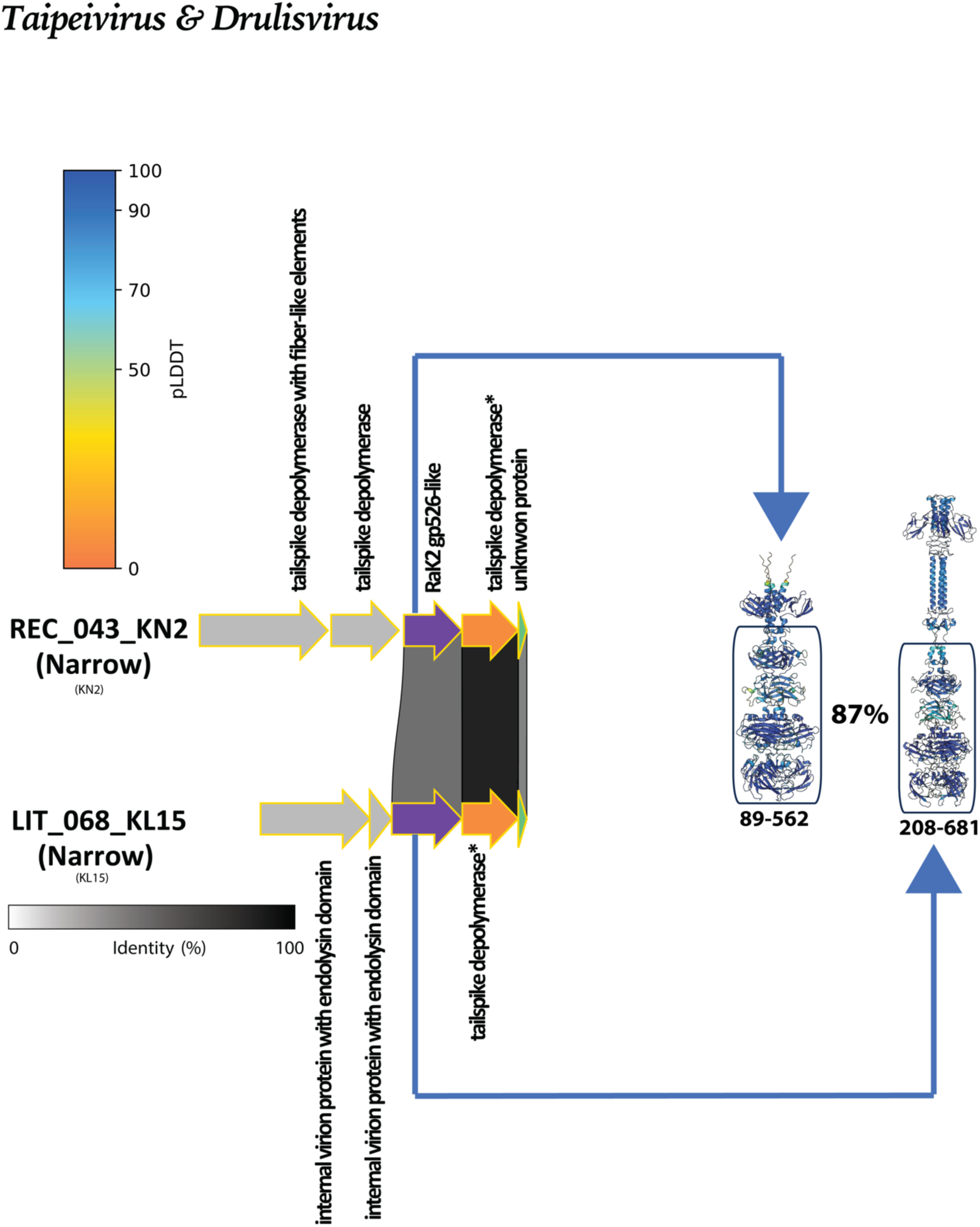
Cross-morphotype sharing of a RaK2 gp526-like RBP between *Taipeivirus* and *Drulisvirus* reveals recent transfer of an uncharacterised receptor-recognition module across myovirus and podovirus lineages. Gene neighborhoods of a *Taipeivirus* myovirus and a *Drulisvirus* podovirus (drawn to scale; arrows indicate ORFs and transcriptional direction) with pairwise amino acid identity across the locus (grey-scale shading, 0–100% identity), alongside predicted RBP structures colored by residue confidence (pLDDT scale). Square boxes drawn between the two RBPs represent regions of amino acid sequence similarity; asterisks (*) denote proteins for which AlphaFold3 generated poor-quality structural predictions. REC_043_KN2 (Narrow host range; KN2) encodes a multi-RBP locus comprising a tailspike depolymerase with fiber-like elements, a tailspike depolymerase, a RaK2 gp526-like RBP, and a tailspike depolymerase flanked by an unknown protein, while LIT_068_KL15 (Narrow host range; KL15) encodes a minimal locus with two internal virion proteins carrying endolysin domains paired with a single tailspike depolymerase. The shared region corresponds to the RaK2 gp526-like RBP (residues 89–562 and 208–681, respectively), a class first described in the jumbophage *Enterobacteria phage* vB_KleM-RaK2 and one of only four RBP classes in this dataset spanning more than one morphotype. The 87% amino acid identity between these representatives from phylogenetically distant genera (different morphotypes) indicates very recent transfer of this module and demonstrates that convergent receptor-recognition strategies — targeting a receptor whose identity remains to be determined — can be shared across deep taxonomic boundaries in *Klebsiella* phage populations.

**Figure S25:**
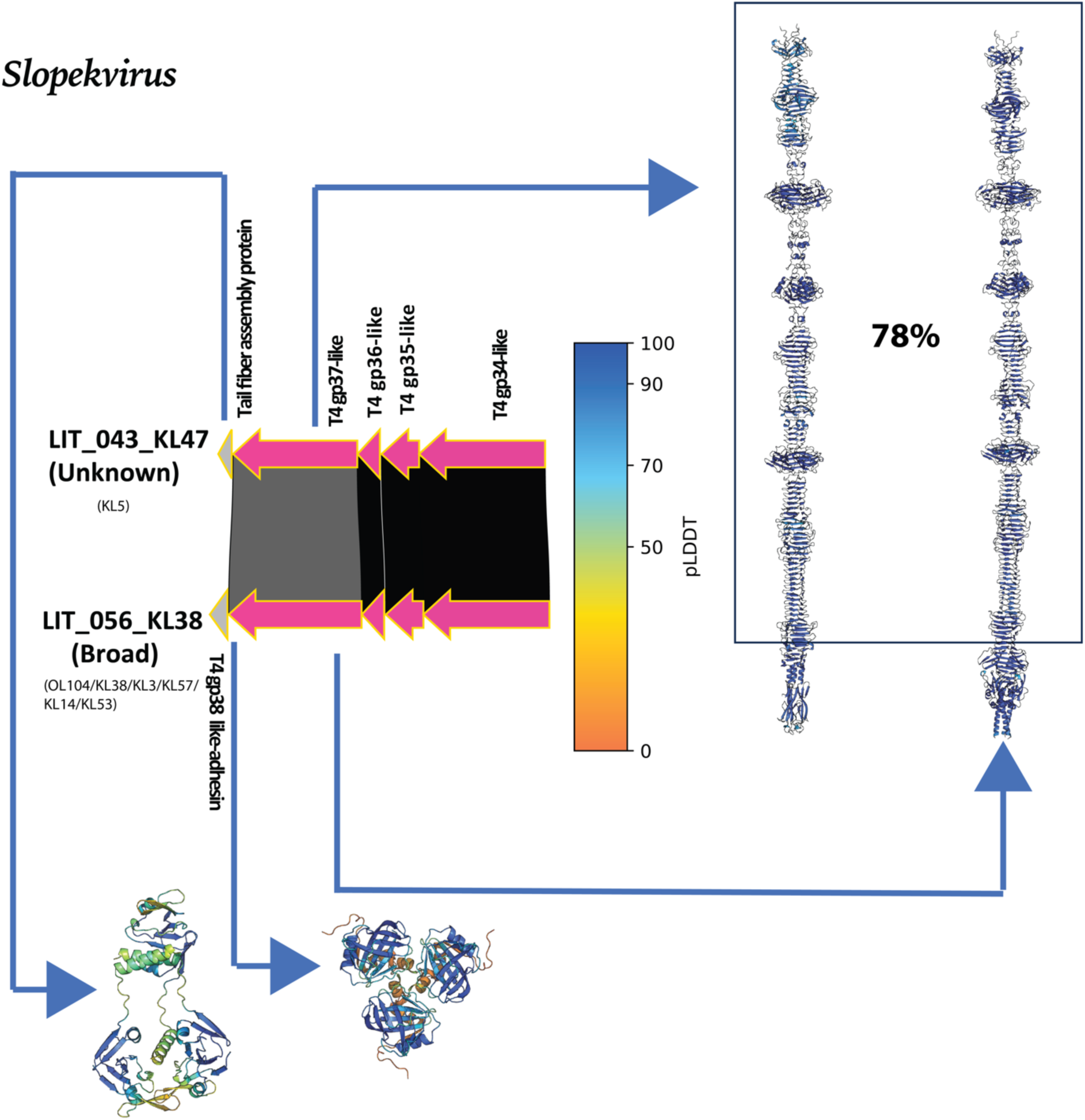
Modular C-terminal substitution within the T4 gp37-like tail fiber class in *Slopekvirus*. Gene neighborhoods of two *Slopekvirus* phages (drawn to scale; arrows indicate ORFs and transcriptional direction) with pairwise amino acid identity across the locus (grey-scale shading, 0–100% identity), alongside predicted RBP structures colored by residue confidence (pLDDT scale). Square boxes drawn between the two RBPs represent regions of amino acid sequence similarity. LIT_043_KL47 (Unknown host range; KL5) carries the canonical T4 gp37-like tail fiber with a conserved internal module (T4 gp34–gp37-like subunits) and a tail fiber assembly protein, while LIT_056_KL38 (Broad host range; OL104/KL38/KL3/KL57/KL14/KL53) carries a divergent C-terminal receptor-binding adhesin (T4 gp38-like adhesin) in place of the canonical C-terminal module. The 78% structural similarity between the resulting trimeric fiber assemblies (right) indicates that C-terminal domain identity co-evolves with the broader tail fiber assembly pathway.

## Supplementary Tables

- **Table S1:** Genome table with all the relevant metadata of the 192 phage genomes used in the study.
- **Table S2:** Metadata information of all the 457 putative RBPs detected in the study.
- **Table S3:** Diverse pseudo-domain clusters share identical ECOD domain assignments when mapped across structurally distinct RBP classes.
- **Table S4:** Fixed-effect results from a binomial GLMM relating genetic host-range (#SSBH-fold depolymerases) to ECOD domain content in RBPs. Host-range is coded as 1 for narrow (phages with 1-2 SSBH-fold depolymerases) and 0 for I/B (phages with either 0 or > 2 SSBH-fold depolymerases) and modelled with a binomial error distribution and a logit link. Predictors indicate the presence versus absence of ECOD domains detected in at least five phage clusters. Phage cluster is included as a random intercept to account for relatedness among genomes. The table reports odds ratios and corresponding 95% confidence intervals derived from the fixed-effect coefficients, together with p-values for each predictor.
- **Table S5:** Table comprising 238 SSBH-fold depolymerases detected across 192 phage genomes, with their manually identified N-terminal coordinates. N-terminal coordinates were identified based on the approach previously reported for *Klebsiella* phage SSBH-fold depolymerases^27,83^.

## Supplementary Text

- **Text S1:** A detailed methodology and structural description of RBP classification.

