## supplementary text - S1 for "Structural modularity of receptor-binding proteins underlies host-range strategy diversification in *Klebsiella pneumoniae* phages"

### Text S1: Detailed Description of RBP-Class Definitions

In this following section we will describe in detail how we defined each RBP-class. We used multiple assignment strategies to define RBP-classes from strong matches to PDB100, tail fiber atlas, matches to reported structures in relevant phage literature, remote similarities to tail fiber atlas, and functional inference via guilt by association. They comprise RBPs spanning multiple phage morphotypes as well as morphotype specific ones too. For convenience, we present here first the common RBPs, and then present morphotype-specific ones in the order of different functional assignment strategies.

#### Part I: Common RBP-Classes (Spanning Multiple Phage Morphotypes)

These four RBP-classes were identified across more than one phage morphotype and together account for the large majority of all RBPs in the dataset (n=227, 60%).

##### 1. Tailspike depolymerase

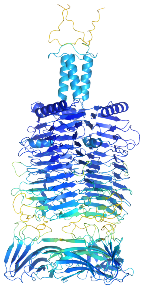

**Definition:** Tailspike depolymerases were identified manually by the presence of a single-stranded beta-helix (SSBH)-fold at the central domain — a structural hallmark of capsule-degrading enzymatic RBPs. Members of this class exhibit a canonical beta-solenoid spine terminating in a C-terminal receptor-binding domain without additional fiber-like structural elements. Foldseek clustering (multimer TMscore threshold 0.65) then partitioned these RBPs into individual clusters.

**Total representatives:** 186 RBPs

**RBP clusters:** 71 Foldseek clusters

**Morphotypes:** All four (Podovirus, Siphovirus, Myovirus, Jumbophage)

**Genera:** *Alcyoneusvirus, Bimevirus, Carvajevirus, Drulisvirus, Efbeekayvirus, Eowynvirus, Kayfunavirus, Kaypoctavirus, Koutsourovirus, Lastavirus, Mascletvirus, Mydovirus, Przondovirus, Sircambvirus, Taipeivirus, Webervirus*, and unclassified phages.

**Biological notes:** Tailspike depolymerases constitute the numerically dominant RBP-class in the dataset and are present across all phage morphotypes, with podoviruses contributing the most (n=132) and myoviruses the fewest (n=15). All members are inferred to target capsular polysaccharides (CPS) for K-type-specific infection, consistent with the characteristic halo phenotype produced by enzymatic degradation of the capsule around plaques. The broad phylogenetic distribution across divergent genera implies that this architecture is under strong positive selection as the primary recognition strategy in *K. pneumoniae* phage communities.

#### 2. Tailspike depolymerase with fiber-like elements

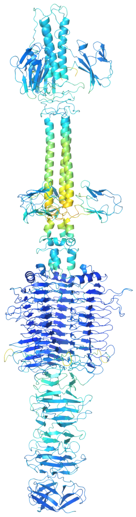

**Definition:** Like the tailspike depolymerase class, members were identified by the presence of an SSBH fold at the central domain. They were distinguished from canonical tailspike depolymerases based on variations in their C-terminal architecture, which contains additional structural elements resembling tail fibers. Foldseek clustering then partitioned these RBPs into individual clusters.

**Total representatives:** 35 RBPs

**RBP clusters:** 15 Foldseek clusters

**Morphotypes:** All four (Podovirus, Siphovirus, Myovirus, Jumbophage)

**Genera:** *Alcyoneusvirus, Drulisvirus, Eowynvirus, Przondovirus, Taipeivirus, Webervirus*, and unclassified phages.

**Biological notes:** This class retains the SSBH-fold enzymatic core of standard tailspike depolymerases but incorporates C-terminal domains with structural resemblance to tail fibers, suggesting a hybrid architecture that may modulate receptor engagement mode or broaden receptor compatibility. Members co-occur with canonical tailspike depolymerases and other non-depolymerase RBP-classes in multi-RBP genomes, particularly in *Taipeivirus* and *Alcyoneusvirus*. Like the canonical tailspike depolymerase class, this class is distributed across all four phage morphotypes, with podoviruses contributing the most (n=14) and myoviruses the fewest (n=6).

#### 3. RaK2 gp526-like

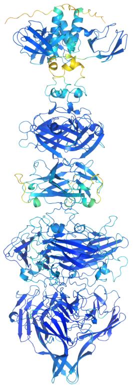

**Definition:** Defined by structural similarity to the characterised RBP gp526 of the JUMBOphage RaK2, a well-described *Klebsiella* phage whose RBP repertoire has been structurally studied in detail. Classification was based on direct comparison to published RBP structures from the RaK2 phage in the literature.

**PMID:** 37298271

**Total representatives:** 3 RBPs

**RBP clusters:** 1 Foldseek cluster (RBP71)

**Morphotypes:** Jumbophage, Myovirus, Podovirus

**Genera:** *Alcyoneusvirus* (jumbophage), *Taipeivirus* (myovirus), *Drulisvirus* (podovirus)

**Biological notes:** The RaK2 gp526-like class is notable for spanning three distinct phage morphotypes — jumbophage, myovirus, and podovirus — within a single Foldseek cluster, suggesting a conserved structural scaffold that has been recruited independently across architecturally distinct virion types. Structurally, members of this RBP-class lack depolymerase domains and instead feature an N-terminal six-stranded beta-sandwich resembling the CBA120 TSP1 D1 head domain, a body domain similar to the *Pseudomonas aeruginosa* R1 pyocin fiber Knob 2, and a C-terminal beta-sandwich resembling the *Pantoea stewartii* WceF glycan biofilm-modifying enzyme. The receptor target of this class in *K. pneumoniae* phages remains experimentally unconfirmed. In *Taipeivirus*, a tailspike depolymerase with fiber-like elements from the same genus shares 72% N-terminal sequence identity with RaK2 gp526-like RBPs, diverging completely beyond this region — consistent with a modular recombination event at a conserved N-terminal scaffold.

#### 4. GDSL-like lipase containing tailspike

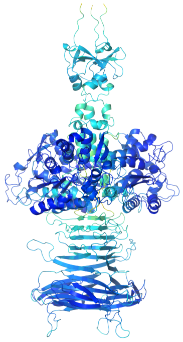

**Definition:** Defined by Foldseek structural search against PDB100 (v20240101), using the Foldseek cluster representative as the query. A significant match (TMscore >= 0.5) to an SGNH hydrolase/GDSL-like lipase-containing tailspike structure in PDB100 was the basis of classification.

**PDB100 best hit:** IVJG (all clusters)

**Total representatives:** 3 RBPs

**RBP clusters:** 2 Foldseek clusters (RBP6, RBP36)

**Morphotypes:** Myovirus and Podovirus

**Genera:** *Mydovirus* (myovirus), *Przondovirus* (podovirus)

**Biological notes:** GDSL-like lipase-containing tailspikes are found in both myovirus (*Mydovirus*) and podovirus (*Przondovirus*) morphotypes, distributed across two distinct Foldseek clusters, implying independent recruitment of the same SGNH hydrolase catalytic module into different RBP scaffolds in these two lineages. Characterised homologues of SGNH hydrolase-containing

RBPs — including the *E. coli* phage G7C gp63.1 tailspike — experimentally remove O-acetyl modifications from polysaccharide substrates, and direct functional evidence for this class in *Klebsiella* temperate phages has recently been reported. Members of this class invariably co-occur with at least one canonical SSBH-fold tailspike depolymerase or tailspike depolymerase with fiber-like elements in the same phage genome. In *Przondovirus*, co-occurrence with SSBH-fold depolymerases correlates with a narrow host-range phenotype, suggesting that the host-range outcome depends on the full RBP repertoire rather than the GDSL-like lipase tailspike alone.

#### Part II: Morphotype-Specific RBP-Classes

The remaining 35 RBP-classes are each restricted to a single phage morphotype. These are presented below grouped by morphotype, within each group ordered by the functional assignment strategy used.

##### IIa. Strong Matches to PDB100 (Foldseek+PDB100, TMscore ≥ 0.5)

For RBPs to which a confident tail fiber class (TC) assignment could not be made, we ran Foldseek structural search against PDB100 (v20240101) using the cluster representative as query. A hit was considered significant only at a minimum TMscore of 0.5. This approach defined 8 morphotype-specific RBP-classes across podoviruses, siphoviruses, and myoviruses.

**PODOVIRUS**

###### 5. HRP29 tail-like variant 1

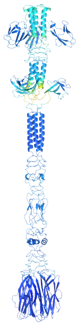

**Definition:** Defined by a significant Foldseek hit (TMscore >= 0.5) of the cluster representative against PDB100, where the best-scoring match corresponded to structural components of phage HRP29 (a *Klebsiella* phage). Two structurally distinct variants were identified within what was initially a broad HRP29-like match; variant 1 constitutes the more abundant group.

**PDB100 best hit:** 8ES4 (all clusters)

**Total representatives:** 5 RBPs

**RBP clusters:** 5 Foldseek clusters (RBP12, RBP40, RBP123, RBP124, RBP127)

**Morphotypes:** Podovirus

**Genera:** *Drulisvirus, Koutsourovirus*

**Biological notes:** HRP29 tail-like variant 1 RBPs share an N-terminal region homologous to the HRP29 gp44 tailspike adapter domain but diverge structurally in their overall architecture. The presence of this class in *Drulisvirus* and *Koutsourovirus* — genera otherwise dominated by SSBH-fold depolymerases — suggests a possible non-capsular receptor target (e.g. O-antigen or LPS components). Each of the five representatives occupies its own Foldseek cluster, indicating substantial structural diversity within this class, which may reflect rapid C-terminal diversification at the receptor-binding domain level. The receptor of HRP29 tail-like variant 1 in *K. pneumoniae* phages has not been experimentally confirmed.

#### 6. HRP29 tail-like variant 2

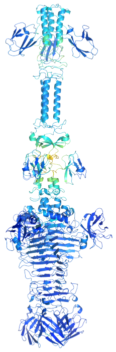

**Definition:** Defined by the same PDB100 structural match criterion as HRP29 tail-like variant 1, but representing a structurally distinct group that was separated into a second variant class based on divergence in overall architecture from variant 1.

**PDB100 best hit:** 8ES4

**Total representatives:** 1 RBP

**RBP clusters:** 1 Foldseek cluster (RBP69)

**Morphotypes:** Podovirus

**Genera:** *Drulisvirus*

**Biological notes:** HRP29 tail-like variant 2 shares the same N-terminal anchor as variant 1 (homologous to HRP29 gp44 tailspike adapter) but diverges structurally in a manner sufficient to warrant a separate class assignment. A pseudo-domain cluster (RBPPD053) containing the ECOD domain 3856.1.1 (putative tailspike protein Orf210 N-terminal domain) is shared between the canonical tailspike depolymerase class and HRP29 tail-like variant 2, suggesting a common N-terminal ancestry. With only one representative, the receptor identity and biological role of this class remain entirely uncharacterised. The presence of this class in *Drulisvirus* alongside the more abundant tailspike depolymerases and variant 1 highlights the RBP diversity within this genus.

#### 7. Putative esterase containing tailspike

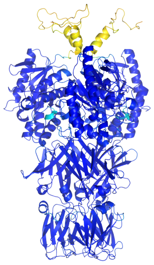

**Definition:** Defined by a significant Foldseek hit (TMscore >= 0.5) of the cluster representative against PDB100, where the best-scoring match corresponded to a structure containing an SGNH hydrolase/esterase domain. Classification as a distinct class from GDSL-like lipase tailspikes reflects differences in the PDB100 hit and overall structural context.

**PDB100 best hit:** 2APJ

**Total representatives:** 4 RBPs

**RBP clusters:** 1 Foldseek cluster (RBP4)

**Morphotypes:** Podovirus

**Genera:** *Przondovirus*

**Biological notes:** All four representatives fall within a single Foldseek cluster and are found exclusively in *Przondovirus*. SGNH hydrolase domain-containing RBPs with O-antigen specificity have been characterised in *E. coli* phages, and structurally analogous esterase-containing RBPs are inferred to target O-antigen or surface polysaccharide modifications rather than the capsule directly. In *Przondovirus*, the putative esterase tailspike co-occurs with canonical SSBH-fold depolymerases; the host-range phenotype of phages encoding this class is narrow, consistent with co-dependence on the depolymerase for capsule clearance. The specific bacterial receptor and substrate remain to be confirmed experimentally.

SIPHOVIRUS

#### 8. L-shaped tail fiber-like

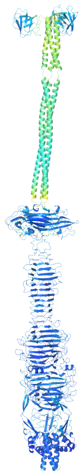

**Definition:** Defined by a significant Foldseek hit (TMscore >= 0.5) of the cluster representative against PDB100, where the best-scoring match corresponded to L-shaped tail fiber structures from well-characterised phages. The characteristic bent geometry that distinguishes L-shaped fibers from straight tail fibers was confirmed in the structural models.

**PDB100 best hit:** 4UW8 (all clusters)

**Total representatives:** 8 RBPs

**RBP clusters:** 3 Foldseek clusters (RBP105, RBP110, RBP112)

**Morphotypes:** Siphovirus

**Genera:** *Webervirus*, and one unclassified phage

**Biological notes:** The L-shaped tail fiber-like class is structurally similar to L-shaped tail fibers of phage T5, which recognise oligo-mannose units of the O-antigen on the *E. coli* surface. By inference, L-shaped tail fiber-like RBPs in *Klebsiella* phages are proposed to target oligo-mannose O-antigen residues rather than the capsule, classifying them as O-antigen generalists. Consistent with this receptor inference, *Webervirus* phages encoding the L-shaped tail fiber-like class show substantially broader empirical host-range than *Webervirus* members encoding canonical SSBH-fold depolymerases — a pattern explained by the limited O-antigen locus diversity in *K. pneumoniae* (13 O-loci) compared to the extensive capsular locus diversity (>150 KL-types). Within *Webervirus*, near-identical phages differing only at the RBP locus encode either an SSBH-fold tailspike or an L-shaped tail fiber, with the fiber-carrying variant shifting from narrow to broad host-range. The three Foldseek clusters spanning this class reflect structural divergence within the receptor-binding domain while retaining the shared L-shaped overall architecture.

#### 9. SU10 tail fiber-like

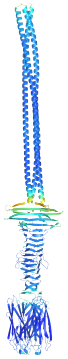

**Definition:** Defined by a significant Foldseek hit (TMscore >= 0.5) of the cluster representative against PDB100, where the best-scoring match corresponded to the long tail fiber of phage SU10, a *Klebsiella* phage. The N-terminal domain of this RBP-class exhibits coiled-coil structural similarity to the SU10 long tail fiber.

**PDB100 best hit:** 7Z4B

**Total representatives:** 1 RBP

**RBP clusters:** 1 Foldseek cluster (RBP9)

**Morphotypes:** Siphovirus

**Genera:** *Webervirus*

**Biological notes:** The SU10 tail fiber-like class is represented by a single RBP from a single *Webervirus* isolate. The structural similarity to SU10 long tail fiber (PDB match) and N-terminal coiled-coil features are consistent with a long, straight fiber architecture distinct from the L-shaped class found in the same genus. The receptor target of SU10 tail fiber-like RBPs in *Webervirus* has not been experimentally determined. Its co-occurrence alongside tailspike depolymerases in *Webervirus* underscores the structural diversity of the RBP repertoire within this genus.

##### MYOVIRUS

#### 10. GDSL-like lipase containing tail fiber

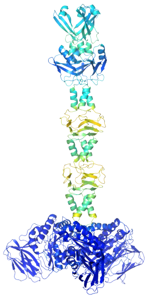

**Definition:** Defined by a significant Foldseek hit (TMscore >= 0.5) of the cluster representative against PDB100, where the best-scoring match indicated an SGNH hydrolase/GDSL-like lipase domain embedded in a tail fiber scaffold (as opposed to a tailspike). Classified as a distinct RBP-class from the cross-morphotype GDSL-like lipase containing tailspike class based on structural differences in overall architecture and morphotype restriction to myovirus.

**PDB100 best hit:** 4K9S

**Total representatives:** 1 RBP

**RBP clusters:** 1 Foldseek cluster (RBP34)

**Morphotypes:** Myovirus

**Genera:** *Mydovirus*

**Biological notes:** The GDSL-like lipase containing tail fiber is the myovirus-restricted counterpart to the GDSL-like lipase containing tailspike (found in both myovirus and podovirus). It is represented by a single RBP in *Mydovirus* and carries an SGNH hydrolase catalytic domain within a tail fiber architectural context rather than a tailspike. By analogy to characterised SGNH hydrolase RBPs, this class is inferred to enzymatically remove surface polysaccharide modifications. Together with the tailspike variant, the presence of SGNH hydrolase domains in both myovirus tail fibers and podovirus/myovirus tailspikes illustrates the repeated modular recruitment of this catalytic module across different RBP scaffolds. The specific receptor substrate remains to be experimentally confirmed.

#### 11. Pam3 fiber-like

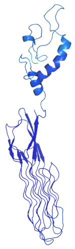

**Definition:** Defined by a significant Foldseek hit (TMscore >= 0.5) of the cluster representative against PDB100, where the best-scoring match corresponded to the fiber protein Pam3 from Myocyanophage, a cyanophage with a characterised monomeric adhesin structure.

**PDB100 best hit: 7YPX**

**Total representatives:** 2 RBPs

**RBP clusters:** 1 Foldseek cluster (RBP28)

**Morphotypes:** Myovirus

**Genera:** *Bimevirus, Mascletvirus*

**Biological notes:** Pam3 fiber-like RBPs are monomeric adhesin proteins with hypervariable C-terminal loops — a structural feature shared with the *Straboviridae* gp38 adhesin — that in related phages recognise diverse outer loop combinations of several outer membrane protein (OMP) receptors. In *K. pneumoniae* phages, pam3 fiber-like RBPs are found in *Bimevirus* and *Mascletvirus* (*Jameshumphriesvirinae*), where they co-occur paired with a tailspike depolymerase but notably without additional non-enzymatic tail fibers. This co-occurrence pattern contrasts with the architecturally analogous *Straboviridae* phages (*Slopekvirus*, *Jiaodavirus*), which carry the full gp34-gp35-gp36-gp37 long tail fiber complex in addition to gp12-like short tail fibers. *Bimevirus* and *Mascletvirus* are proposed to target OmpC or related OMPs via the pam3-like adhesin (Strategy 5: OMP gatekeeper), but the absence of additional non-capsule-targeting fibers means successful OMP engagement is contingent on prior capsule clearance by the co-encoded depolymerase. *Mascletvirus* displays a narrow empirical host-range, consistent with this dependence on depolymerase-mediated capsule removal before OMP access.

#### 12. Xylanase containing tail fiber

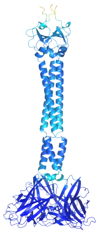

**Definition:** Defined by a significant Foldseek hit (TMscore >= 0.5) of the cluster representative against PDB100, where the best-scoring match corresponded to a xylanase-containing structure. The xylanase domain (a glycoside hydrolase family domain capable of cleaving xylan polymers) is embedded within a tail fiber scaffold.

**PDB100 best hit:** 7AY3

**Total representatives:** 2 RBPs

**RBP clusters:** 1 Foldseek cluster (RBP81)

**Morphotypes:** Myovirus

**Genera:** *Mydovirus*

**Biological notes:** The xylanase containing tail fiber class is an enzymatically active tail fiber restricted to *Mydovirus*. Both representatives fall within a single Foldseek cluster. Like other enzymatically active non-depolymerase RBP-classes identified here (GDSL-like lipase, putative esterase), the xylanase domain is inferred to target surface polysaccharide components, though the specific substrate and whether it targets capsular or non-capsular glycans in *K. pneumoniae* remains to be experimentally determined. Xylanases cleave beta-1,4-xylosidic linkages in xylan polysaccharides; their presence in a tail fiber context suggests a surface-remodelling function that may facilitate phage adsorption to the bacterial surface.

#### IIb. Tail Fiber Atlas Matches (RBPseg-classify; probability ≥ 0.95, qTMscore ≥ 0.5)

We used the RBPseg classify module to assign confident tail fiber classes (TCs) to RBPs not assigned via the depolymerase identification step. Six TCs were assigned to 72 RBPs at high confidence (probability >= 0.95 and query TMscore >= 0.5), defining six RBP-classes.

##### PODOVIRUS

#### 13. T7 gp12 nozzle-like

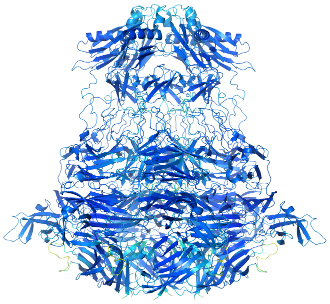

**Definition:** Assigned by RBPseg-classify at high confidence (TC2; probability >= 0.95, qTMscore >= 0.5). The TC2 class in the tail fiber atlas corresponds to nozzle-like hexameric proteins structurally related to phage T7 gp12, a well-characterised component of the T7 tail machinery.

**Total representatives:** 41 RBPs

**RBP clusters:** 1 Foldseek cluster (RBP1)

**Morphotypes:** Podovirus

**Genera:** *Przondovirus, Teetrevirus*

**Biological notes:** The T7 gp12 nozzle-like class is the second most abundant RBP-class in the dataset after tailspike depolymerases, accounting for 22% of all podovirus RBPs. All 41 representatives fall within a single Foldseek cluster (RBP1), indicating strong structural conservation across this class. These RBPs adopt homo-hexameric structures resembling TC2 tail fibers and the HRP29 gp40 nozzle. In the T7 phage, gp12 forms the tail needle nozzle and is involved in host recognition and tail tube assembly; by structural analogy, T7 gp12 nozzle-like RBPs in K. pneumoniae phages are presumed to function in a similar needle/nozzle role. This class is predominantly found in *Przondovirus* (n=40) and one *Teetrevirus* isolate. The receptor targeted by T7 gp12 nozzle-like RBPs in *Klebsiella* phages has not been definitively characterised.

#### 14. TC16 tail fiber

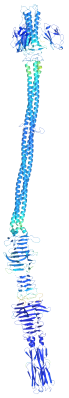

**Definition:** Assigned by RBPseg-classify at high confidence (TC16; probability >= 0.95, qTMscore >= 0.5), mapping to the TC16 class in the RBPseg tail fiber atlas.

**Total representatives:** 1 RBP

**RBP clusters:** 1 Foldseek cluster (RBP100)

**Morphotypes:** Podovirus

**Genera:** *Teetrevirus*

**Biological notes:** The TC16 tail fiber class is represented by a single RBP from a *Teetrevirus* podovirus. Its confident assignment to TC16 by RBPseg-classify places it within a structurally defined tail fiber class in the atlas. *Teetrevirus* also encodes a T7 gp12 nozzle-like RBP (TC2), suggesting that this genus employs at least two distinct tail fiber architectures. A distantly similar class, TC16-like tail fiber, was also found in jumbophages (*Eowynvirus*), but the two classes were kept distinct due to insufficient confidence scores for the latter. The receptor target of TC16 tail fiber in *Klebsiella* phages has not been experimentally determined.

#### MYOVIRUS

##### 15. T4 gp34-like

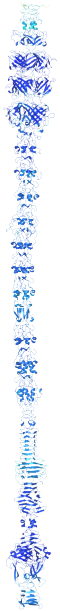

**Definition:** Assigned by RBPseg-classify at high confidence (TC0; probability >= 0.95, qTMscore >= 0.5), mapping to the TC0 class in the RBPseg tail fiber atlas, which corresponds to the T4 long tail fiber proximal subunit gp34.

**Total representatives:** 8 RBPs

**RBP clusters:** 1 Foldseek cluster (RBP49)

**Morphotypes:** Myovirus

**Genera:** *Slopekvirus, Jiaodavirus*

**Biological notes:** T4 gp34-like RBPs correspond to the proximal subunit of the T4 long tail fiber complex and are found exclusively in *Straboviridae* myoviruses (Slopekvirus, Jiaodavirus). All 8 representatives fall within a single Foldseek cluster, reflecting high structural conservation. In these phages, T4 gp34-like invariably co-occurs with T4 gp36-like, T4 gp37-like, and T4 gp12-like RBPs, reconstituting a near-complete T4-like long tail fiber complex (gp34-gp35-gp36-gp37) with a short tail fiber (gp12). Consistent with T4 biology, the distal tip of the gp37 subunit engages OmpC by structural analogy, proposing OMP-targeted adsorption (Strategy 5: OMP gatekeeper) for these phages. In *Slopekvirus* and *Jiaodavirus*, this long tail fiber complex is the primary non-depolymerase host-recognition apparatus, and phages carrying it show broad empirical host-range.

#### 16. T4 gp36-like

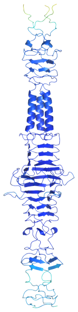

**Definition:** Assigned by RBPseg-classify at high confidence (TC11; probability >= 0.95, qTMscore >= 0.5), mapping to the TC11 class in the tail fiber atlas, corresponding to the T4 long tail fiber intermediate subunit gp36.

**Total representatives:** 8 RBPs

**RBP clusters:** 1 Foldseek cluster (RBP50)

**Morphotypes:** Myovirus

**Genera:** *Slopekvirus, Jiaodavirus*

**Biological notes:** T4 gp36-like RBPs correspond to the intermediate subunit of the T4 long tail fiber and are found in the same genera (*Slopekvirus, Jiaodavirus*) and same phages as T4 gp34-like. All 8 representatives fall within a single Foldseek cluster. Gp36 in T4 forms a short globular connector between gp34 and gp37 within the long tail fiber; the structural conservation of this subunit across *Klebsiella* myoviruses underscores the functional integration of the full T4-like long tail fiber complex. As with T4 gp34-like, the biological role of T4 gp36-like is inferred from its context within the co-encoded T4-like tail fiber gene cluster (gp34-gp35-gp36-gp37).

#### 17. TC8 tail fiber

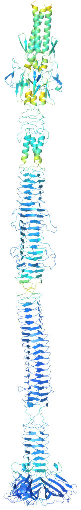

**Definition:** Assigned by RBPseg-classify at high confidence (TC8; probability >= 0.95, qTMscore >= 0.5), mapping to the TC8 class in the RBPseg tail fiber atlas.

**Total representatives:** 7 RBPs

**RBP clusters:** 1 Foldseek cluster (RBP33)

**Morphotypes:** Myovirus

**Genera:** *Mydovirus*

**Biological notes:** TC8 tail fibers are *Vequintavirinae*-specific and are found exclusively in *Mydovirus*. All 7 representatives fall within a single Foldseek cluster (RBP33). The C-terminal domain of TC8 tail fibers may serve as an adapter for tail tip adhesins or longer fiber complexes, analogous to adaptors described in T4-related phages. This RBP-class is structurally distinct from the *vequintavirinae*-like short tail fiber (TC1) class also found in *Mydovirus*, suggesting that this genus encodes at least two architecturally distinct tail fiber types. The receptor target of TC8 tail fiber in *K. pneumoniae* phages has not been experimentally confirmed.

##### 18. *Vequintavirinae*-like short tail fiber

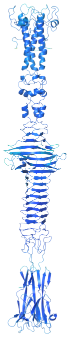

**Definition:** Assigned by RBPseg-classify at high confidence (TC1; probability >= 0.95, qTMscore >= 0.5), mapping to the TC1 class in the RBPseg tail fiber atlas, which corresponds to short tail fibers of *Vequintavirinae*.

**Total representatives:** 7 RBPs

**RBP clusters:** 1 Foldseek cluster (RBP32)

**Morphotypes:** Myovirus

**Genera:** *Mydovirus*

**Biological notes:** The *vequintavirinae*-like short tail fiber class is restricted to *Mydovirus* and all 7 representatives cluster into a single Foldseek cluster (RBP32). This class corresponds to short tail fibers of the *Vequintavirinae* subfamily, with C-terminal resemblance to the Mu phage tail fiber. In T4-like phages, short tail fibers are typically involved in reversible initial contact and host surface scanning before irreversible adsorption. In *Mydovirus*, this class co-occurs with TC8 tail fibers and SSBH-fold depolymerases, contributing to the exceptionally diverse RBP repertoire (up to eleven RBP-classes per genome) observed in this genus. A shared pseudo-domain cluster (RBPPD011, ECOD domain 79.1.1: phage tail fiber protein trimerisation domain) links this class to other structural groups, suggesting a common evolutionary scaffold. The receptor target in *K. pneumoniae* phages has not been experimentally determined.

#### IIc. Reported Structures in Relevant Phage Literature

For RBPs that received no confident TC assignment and no significant PDB100 hit, we assigned RBP-classes based on structural similarity to phage RBPs described in published literature. This approach defined three morphotype-specific RBP-classes (T4 gp12-like in myoviruses, and RaK2 gp534-like and RaK2 gp98-like in jumbophages). The cross-morphotype RaK2 gp526-like class was also defined by this strategy and is described in Part I above.

MYOVIRUS

#### 19. T4 gp12-like

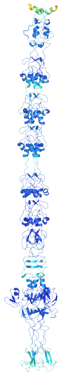

**Definition:** Defined by direct structural comparison to the short tail fiber gp12 of phage T4, based on published structural and functional data from T4 phage. Classification was supported by co-occurrence of gp12-like proteins alongside T4 gp34-like, gp36-like, and gp37-like proteins within the same phage genomes.

**PDB100 best hit:** PMID: 27193680

**Total representatives:** 5 RBPs

**RBP clusters:** 1 Foldseek cluster (RBP48)

**Morphotypes:** Myovirus

**Genera:** *Slopekvirus*

**Biological notes:** T4 gp12-like proteins correspond to the short tail fibers of phage T4, which make final irreversible contact with the LPS inner core of the bacterial surface and function in tandem with the long tail fiber complex during adsorption. In *K. pneumoniae* phages, T4 gp12-like RBPs are found exclusively in *Slopekvirus* myoviruses. In T4, gp12 targets the heptose-rich LPS inner core directly; by structural analogy, *Slopekvirus* phages carrying this RBP-class are inferred to target the conserved LPS inner core (Strategy 6: LPS core anchor). Supporting this inference, five of seven *Slopekvirus* phages in the KlebPhaCol collection were isolated on capsule-null hosts and two on encapsulated strains, consistent with LPS core access being facilitated by, but not requiring, reduced capsule density. A shared pseudo-domain cluster (RBPPD026, ECOD domain 1083.1.1: Phage T4 gp12 N-terminal repeating units) also links this class to the TC17-like tail fiber class, suggesting a common N-terminal structural ancestry. All 5 representatives fall within a single Foldseek cluster.

JUMBOPHAGE

#### 20. RaK2 gp534-like

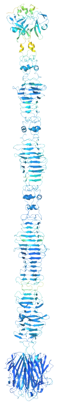

**Definition:** Defined by structural similarity to the RBP gp534 of phage RaK2, a giant *Klebsiella* phage whose RBP repertoire has been characterised in detail in published literature.

**PDB100 best hit:** PMID: 37298271

**Total representatives:** 3 RBPs

**RBP clusters:** 3 Foldseek clusters (RBP121, RBP140, RBP144)

**Morphotypes:** Jumbophage

**Genera:** *Alcyoneusvirus*

**Biological notes:** RaK2 gp534-like RBPs are found exclusively in *Alcyoneusvirus* jumbophages, distributed across three distinct Foldseek clusters — indicating notable structural diversity within this class at the inter-cluster level. The N-terminal domain resembles the CBA120 TSP3 D2 head domain, while the C-terminal domain adopts a jelly-roll fold — a beta-barrel topology commonly used for receptor binding by diverse phage RBPs. *Alcyoneusvirus* genomes encode multiple depolymerases in addition to non-depolymerase RBPs such as RaK2 gp534-like and RaK2 gp98-like, achieving broad host-range through RBP repertoire accumulation (Strategy 2: K-type arsenal) rather than individual RBP promiscuity. The specific *K. pneumoniae* receptor recognised by RaK2 gp534-like RBPs has not been experimentally confirmed.

#### 21. RaK2 gp98-like

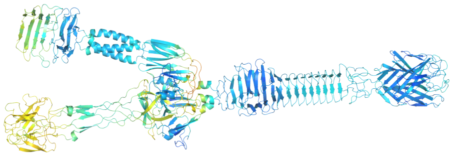

**Definition:** Defined by structural similarity to the RBP gp98 of phage RaK2, based on published structural characterisation of the RaK2 phage RBP repertoire.

**PDB100 best hit:** PMID: 37298271

**Total representatives:** 3 RBPs

**RBP clusters:** 2 Foldseek clusters (RBP116, RBP141)

**Morphotypes:** Jumbophage

**Genera:** *Alcyoneusvirus*

**Biological notes:** RaK2 gp98-like RBPs are also restricted to *Alcyoneusvirus* jumbophages. This class presents a novel architecture with a compact jelly-roll receptor-binding domain at the C-terminus, structurally distinct from known straight-fiber classes. The jelly-roll fold at the C-terminus, shared with RaK2 gp534-like, suggests that both *Alcyoneusvirus*-exclusive literature-defined classes employ this versatile beta-barrel topology for receptor engagement, despite having distinct overall architectures. As noted for RaK2 gp534-like, these RBPs operate in the context of a massively encoded RBP arsenal in *Alcyoneusvirus*, and their specific receptor targets in *K. pneumoniae* phages remain experimentally uncharacterised.

#### IId. Remote Similarities to Tail Fiber Atlas (RBPseg likely TC; qTMscore < 0.5)

For RBPs with no significant PDB100 hit and no literature precedent, we assigned RBP-classes based on distant TC similarities detected by the RBPseg-classify module operating at relaxed thresholds (probability <= 1 and qTMscore < 0.5). These represent structurally novel RBP-classes with only remote structural homology to characterised tail fiber classes, and are designated as candidates for future experimental characterisation. This approach defined 18 RBP-classes, presented below by morphotype.

###### MYOVIRUS

##### 22. T4 gp37-like

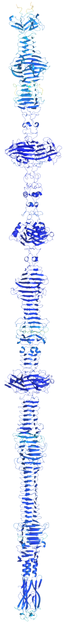

**Definition:** Assigned based on remote similarity to the TC class corresponding to T4 gp37 via RBPseg-classify at relaxed thresholds. Notably, this class is classified as structurally novel: the structure described here is substantially different from the T4 gp37 structure in published literature, and there is no direct literature precedent for this class in *Klebsiella* phages. Assignment is therefore also supported by guilt-by-association inference: T4 gp37-like proteins invariably co-occur with T4 gp34-like, T4 gp36-like, and T4 gp12-like RBPs in the same phage genomes, consistent with participation in a T4-like long tail fiber system.

**Total representatives:** 8 RBPs

**RBP clusters:** 4 Foldseek clusters (RBP51, RBP66, RBP76, RBP94)

**Morphotypes:** Myovirus

**Genera:** *Slopekvirus, Jiaodavirus*

**Biological notes:** T4 gp37-like RBPs are the distal subunit of the T4 long tail fiber system in *K. pneumoniae Straboviridae* myoviruses (*Slopekvirus, Jiaodavirus*). In T4, gp37 carries the receptor-binding tip of the long tail fiber, with the C-terminal domain directly engaging OmpC. By co-occurrence logic and structural analogy, T4 gp37-like RBPs in *Klebsiella* phages are proposed to mediate OmpC recognition (Strategy 5: OMP gatekeeper). The four distinct Foldseek clusters within this class reflect greater structural diversification at the distal tip compared to the proximal (gp34-like) and intermediate (gp36-like) subunits, consistent with rapid C-terminal evolution driven by host receptor diversity. In some *Slopekvirus*, the T4 gp37-like tail fiber carries an intramolecular chaperone (IMC) domain, but in a minority variant this is replaced by a receptor-binding adhesin, co-occurring with substitution of the tail fiber assembly protein by a gp38-like adhesin — suggesting co-evolution of C-terminal domain identity with the broader assembly pathway.

#### 23. TC10-like tail fiber in *Mydovirus* — variant 1

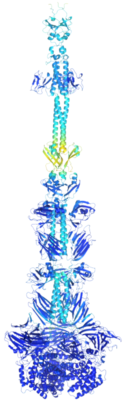

**Definition:** Assigned by RBPseg-classify at relaxed thresholds, with the best distant TC match corresponding to TC10. Classified as variant 1 to distinguish it structurally from two additional TC10-like classes found in *Mydovirus* (variants 2 and 3) and a further TC10-like class in *Alcyoneusvirus* jumbophages.

**Total representatives:** 1 RBP

**RBP clusters:** 1 Foldseek cluster (RBP30)

**Morphotypes:** Myovirus

**Genera:** *Mydovirus*

**Biological notes:** TC10-like tail fiber variant 1 is one of three TC10-like classes found in *Mydovirus* myoviruses, reflecting the exceptional RBP diversity of this genus (which harbours up to eleven distinct RBP-classes). Each variant represents a structurally distinct group with only remote similarity to the TC10 reference class in the tail fiber atlas. A pseudo-domain cluster (RBPPD057, ECOD domain 3856.1.1: putative tailspike protein Orf210 N-terminal domain) links TC10-like variant 3 in *Mydovirus* to other RBP-classes, suggesting some degree of shared N-terminal ancestry. The receptor target of this class is unknown.

#### 24. TC10-like tail fiber in *Mydovirus* — variant 2

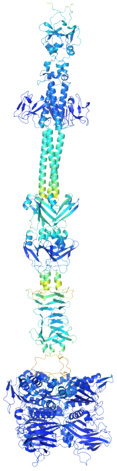

**Definition:** Assigned by RBPseg-classify at relaxed thresholds; remote TC10 match. Classified as a separate variant from TC10-like variants 1 and 3 based on structural divergence at cluster level.

**Total representatives:** 1 RBP

**RBP clusters:** 1 Foldseek cluster (RBP79)

**Morphotypes:** Myovirus

**Genera:** *Mydovirus*

**Biological notes:** A second structurally distinct TC10-like tail fiber in *Mydovirus*, this class contributes to the remarkable RBP repertoire diversity within this genus. Like variant 1 and variant 3, this class carries only a remote structural resemblance to the TC10 reference and is classified as structurally novel. The receptor target is unknown.

#### 25. TC10-like tail fiber in *Mydovirus* — variant 3

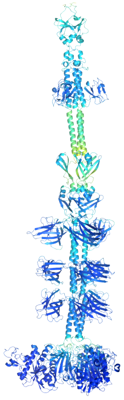

**Definition:** Assigned by RBPseg-classify at relaxed thresholds; remote TC10 match. Classified as variant 3 based on structural divergence from variants 1 and 2.

**Total representatives:** 1 RBP

**RBP clusters:** 1 Foldseek cluster (RBP82)

**Morphotypes:** Myovirus

**Genera:** *Mydovirus*

**Biological notes:** The third TC10-like variant in *Mydovirus*. A pseudo-domain cluster (RBPPD057, ECOD domain 3856.1.1: putative tailspike protein Orf210 N-terminal domain) is shared between this variant and additional RBP-classes, indicating a conserved N-terminal module. The receptor target is unknown.

#### 26. TC12-like tail fiber in *Mydovirus*

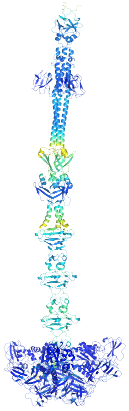

**Definition:** Assigned by RBPseg-classify at relaxed thresholds, with the best distant TC match corresponding to TC12. Classified separately from the TC12-like tail fiber in podovirus due to morphotype restriction and structural divergence.

**Total representatives:** 3 RBPs

**RBP clusters:** 1 Foldseek cluster (RBP78)

**Morphotypes:** Myovirus

**Genera:** *Mydovirus*

**Biological notes:** TC12-like tail fiber in *Mydovirus* myoviruses carries only remote structural similarity to the TC12 reference class, and is classified as structurally novel. All 3 representatives cluster within a single Foldseek cluster. This class was detected as sharing sequence-level modularity with additional RBP-classes previously undetected by ECOD or pseudo-domain methods, suggesting connectivity at a relatively recent evolutionary timescale. The receptor target is unknown.

#### 27. TC12/TC17-like tail fiber in *Mydovirus*

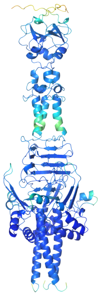

**Definition:** Assigned by RBPseg-classify at relaxed thresholds, where the cluster representative showed ambiguous distant similarity to both TC12 and TC17 classes in the tail fiber atlas. The combined designation TC12/TC17-like reflects this dual remote match.

**Total representatives:** 7 RBPs

**RBP clusters:** 1 Foldseek cluster (RBP31)

**Morphotypes:** Myovirus

**Genera:** *Mydovirus*

**Biological notes:** TC12/TC17-like is one of the more numerically abundant novel RBP-classes in *Mydovirus*, with all 7 representatives in a single Foldseek cluster. The ambiguous dual-TC assignment indicates that this class falls between the TC12 and TC17 reference clusters in structural space, without confidently belonging to either. A shared pseudo-domain cluster (RBPPD088, ECOD domain 3240.1.1: intramolecular chaperone domain in virus tail spike protein) is shared with TC15-like variant 1 in siphovirus, TC7-like, and tailspike depolymerase with fiber-like elements — suggesting a conserved internal chaperone-like module spanning multiple RBP-classes. The receptor target is unknown.

#### 28. TC14-like long tail fiber in *Taipeivirus*

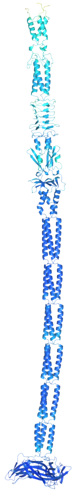

**Definition:** Assigned by RBPseg-classify at relaxed thresholds; remote TC14 match. Designated "long" to distinguish from the structurally distinct TC14-like short tail fiber class in the same genus (*Taipeivirus*).

**Total representatives:** 3 RBPs

**RBP clusters:** 1 Foldseek cluster (RBP96)

**Morphotypes:** Myovirus

**Genera:** *Taipeivirus*

**Biological notes:** TC14-like long tail fiber is one of two TC14-like classes found in *Taipeivirus* myoviruses, which are notable for encoding a diverse RBP complement including SSBH-fold depolymerases, depolymerases with fiber-like elements, and both TC14-like tail fiber variants. The broad host-range of *Taipeivirus* (>60% of phages classified as intermediate/broad host-range) is proposed to emerge from this multi-class RBP accumulation. This class is classified as structurally novel based on the low qTMscore against the TC14 reference. The receptor target is unknown.

#### 29. TC14-like short tail fiber in *Taipeivirus*

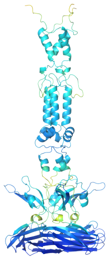

**Definition:** Assigned by RBPseg-classify at relaxed thresholds; remote TC14 match. Designated "short" based on structural size relative to the long tail fiber variant in the same genus.

**Total representatives:** 3 RBPs

**RBP clusters:** 2 Foldseek clusters (RBP99, RBP130)

**Morphotypes:** Myovirus

**Genera:** *Taipeivirus*

**Biological notes:** TC14-like short tail fiber in *Taipeivirus* is structurally distinct from its long counterpart despite sharing a TC14-like distant assignment, as evidenced by clustering into two separate Foldseek clusters. This class was excluded from pseudo-domain segmentation analysis due to insufficient segmentation signal. Together with the TC14-like long tail fiber, this class contributes to the multi-RBP architecture of *Taipeivirus*, a genus proposed to achieve broad host-range through RBP repertoire breadth rather than individual RBP promiscuity. The receptor target is unknown.

#### 30. TC17-like tail fiber

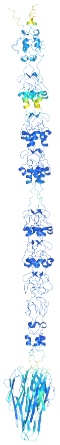

**Definition:** Assigned by RBPseg-classify at relaxed thresholds, with the best distant TC match corresponding to TC17. A shared pseudo-domain cluster (RBPPD026, ECOD domain 1083.1.1: Phage T4 gp12 N-terminal repeating units) was also detected between TC17-like and T4 gp12-like, linking these two classes structurally at the N-terminal domain level.

**Total representatives:** 3 RBPs

**RBP clusters:** 1 Foldseek cluster (RBP65)

**Morphotypes:** Myovirus

**Genera:** *Jiaodavirus*

**Biological notes:** TC17-like tail fibers are found in *Jiaodavirus* myoviruses (*Straboviridae*), which also carry the full T4-like long tail fiber system (gp34-like, gp36-like, gp37-like). The structural link between TC17-like and T4 gp12-like via a shared N-terminal pseudo-domain cluster may indicate an evolutionary relationship or shared structural module between these co-encoded classes. It could also be a C-terminal innovation specific to *Jiaodavirus* phages of T4 gp12-like short tail fiber seen in *Slopekvirus.* Classification as structurally novel reflects the remote nature of the TC17 similarity score. The receptor target in K. pneumoniae phages is unknown.

##### SIPHOVIRUS

#### 31. TC5-like tail fiber

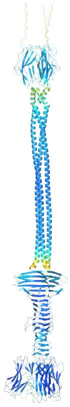

**Definition:** Assigned by RBPseg-classify at relaxed thresholds, with the best distant TC match corresponding to TC5.

**Total representatives:** 1 RBP

**RBP clusters:** 1 Foldseek cluster (RBP43)

**Morphotypes:** Siphovirus

**Genera:** *Webervirus*

**Biological notes:** TC5-like is one of six structurally novel RBP-classes found in siphoviruses. It is represented by a single RBP from a Webervirus isolate — a genus that also encodes L-shaped tail fiber-like and SU10 tail fiber-like RBPs, making *Webervirus* one of the most RBP-diverse siphovirus genera in the collection. The remote TC5 match provides minimal functional inference, and the receptor target of TC5-like in *K. pneumoniae* phages is unknown.

#### 32. TC7-like tail fiber

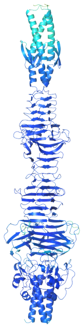

**Definition:** Assigned by RBPseg-classify at relaxed thresholds, with the best distant TC match corresponding to TC7.

**Total representatives:** 3 RBPs

**RBP clusters:** 2 Foldseek clusters (RBP113, RBP114)

**Morphotypes:** Siphovirus

**Genera:** Unclassified phages

**Biological notes:** TC7-like tail fibers are found in two phages that could not be assigned to any established genus. Both Foldseek clusters are structurally distinct, and a shared pseudo-domain cluster (RBPPD088, ECOD domain 3240.1.1: intramolecular chaperone domain in virus tail spike protein) links TC7-like to TC12/TC17-like (*Mydovirus*), TC15-like variant 1 (siphovirus), and tailspike depolymerase with fiber-like elements — suggesting a shared structural module across morphotype boundaries. The receptor target is unknown.

#### 33. TC14-like tail fiber in *Sugarlandvirus* — variant 1

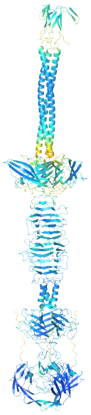

**Definition:** Assigned by RBPseg-classify at relaxed thresholds; remote TC14 match. Classified as variant 1 to distinguish from variant 2 found in the same genus (*Sugarlandvirus*).

**Total representatives:** 3 RBPs

**RBP clusters:** 2 Foldseek clusters (RBP60, RBP95)

**Morphotypes:** Siphovirus

**Genera:** *Sugarlandvirus*

**Biological notes:** TC14-like variant 1 and variant 2 together represent the two structurally distinct TC14-like tail fiber classes found in *Sugarlandvirus* siphoviruses. *Sugarlandvirus* phages carrying these classes show within-genus RBP diversification where different phages encode different TC14-like variants, potentially expanding the receptor repertoire of this genus without changing the overall functional assignment category. The receptor targets are unknown.

#### 34. TC14-like tail fiber in *Sugarlandvirus* — variant 2

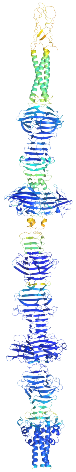

**Definition:** Assigned by RBPseg-classify at relaxed thresholds; remote TC14 match. Classified as a separate variant from variant 1 based on structural divergence, supported by clustering into distinct Foldseek clusters.

**Total representatives:** 3 RBPs

**RBP clusters:** 3 Foldseek clusters (RBP62, RBP63, RBP64)

**Morphotypes:** Siphovirus

**Genera:** *Sugarlandvirus*

**Biological notes:** With three representatives across three separate Foldseek clusters, TC14-like variant 2 displays greater within-class structural diversity than variant 1. Both TC14-like variants in *Sugarlandvirus* illustrate the within-genus RBP diversification strategy whereby closely related phages encode structurally distinct variants of the same broadly assigned TC class, consistent with receptor repertoire expansion through modular C-terminal domain substitution. The receptor target is unknown.

#### 35. TC15-like tail fiber in siphovirus — variant 1

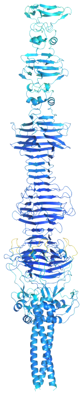

**Definition:** Assigned by RBPseg-classify at relaxed thresholds; remote TC15 match. Two structurally distinct TC15-like classes were identified in siphoviruses and designated variants 1 and 2 accordingly.

**Total representatives:** 1 RBP

**RBP clusters:** 1 Foldseek cluster (RBP52)

**Morphotypes:** Siphovirus

**Genera:** *Henuseptimavirus*

**Biological notes:** TC15-like variant 1 is represented by a single RBP from a *Henuseptimavirus* siphovirus. A shared pseudo-domain cluster (RBPPD088, ECOD domain 3240.1.1: intramolecular chaperone domain in virus tail spike protein) links TC15-like variant 1 to TC12/TC17-like tail fiber in *Mydovirus*, TC7-like tail fiber, and tailspike depolymerase with fiber-like elements — indicating a conserved structural module spanning morphotype boundaries. The receptor target is unknown.

#### 36. TC15-like tail fiber in siphovirus — variant 2

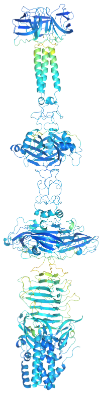

**Definition:** Assigned by RBPseg-classify at relaxed thresholds; remote TC15 match. Classified as a separate variant from variant 1 based on structural divergence.

**Total representatives:** 2 RBPs

**RBP clusters:** 1 Foldseek cluster (RBP111)

**Morphotypes:** Siphovirus

**Genera:** Unclassified phages

**Biological notes:** TC15-like variant 2 is found in two phages that could not be assigned to a named genus. Both fall within a single Foldseek cluster. The structural distinction from variant 1 (which is in *Henuseptimavirus*) and the different genus context indicate that this variant likely represents an independent evolutionary origin of a TC15-like architecture within siphoviruses. The receptor target is unknown.

##### JUMBOPHAGE

#### 37. TC10-like tail fiber in *Alcyoneusvirus*

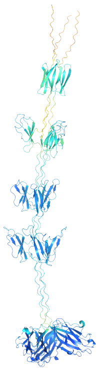

**Definition:** Assigned by RBPseg-classify at relaxed thresholds; remote TC10 match. Classified as a distinct RBP-class from the three TC10-like variants found in *Mydovirus* myoviruses based on morphotype (jumbophage) and structural divergence.

**Total representatives:** 3 RBPs

**RBP clusters:** 1 Foldseek cluster (RBP122)

**Morphotypes:** Jumbophage

**Genera:** *Alcyoneusvirus*

**Biological notes:** TC10-like tail fiber in *Alcyoneusvirus* is one of two uncharacterised RBP-classes in this jumbophage genus, alongside TC16-like in *Eowynvirus*. All 3 representatives cluster within a single Foldseek cluster (RBP122). *Alcyoneusvirus* is dominated by SSBH-fold depolymerases (n=30, 73%) and also carries the literature-defined RaK2 gp534-like and RaK2 gp98-like classes, making the TC10-like class a minor structural component of an otherwise well-characterised RBP repertoire. Its receptor target in *K. pneumoniae* phages is unknown and represents a candidate for future structural and functional validation.

#### 38. TC16-like tail fiber

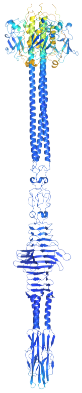

**Definition:** Assigned by RBPseg-classify at relaxed thresholds; remote TC16 match. Classified as a distinct class from TC16 tail fiber (in podoviruses) due to insufficient confidence scores for a direct TC16 assignment.

**Total representatives:** 1 RBP

**RBP clusters:** 1 Foldseek cluster (RBP53)

**Morphotypes:** Jumbophage

**Genera:** *Eowynvirus*

**Biological notes:** TC16-like tail fiber is the sole non-depolymerase RBP class in *Eowynvirus*, a jumbophage genus that otherwise relies on a large repertoire of SSBH-fold depolymerases for host recognition. The single representative clusters into RBP53 and shows only remote structural similarity to the TC16 reference; as such, it is classified as structurally novel. Its receptor target in *K. pneumoniae* phages is unknown.

#### PODOVIRUS

##### 39. TC12-like tail fiber in podovirus

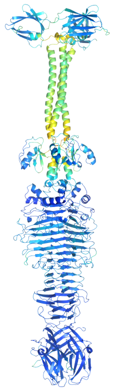

**Definition:** Assigned by RBPseg-classify at relaxed thresholds; remote TC12 match. Classified separately from TC12-like tail fiber in *Mydovirus* based on morphotype (podovirus) and structural divergence.

**Total representatives:** 1 RBP

**RBP clusters:** 1 Foldseek cluster (RBP83)

**Morphotypes:** Podovirus

**Genera:** Unclassified

**Biological notes:** TC12-like tail fiber in a podovirus is the sole uncharacterised RBP-class in podoviruses, found in a phage that could not be assigned to a named genus. This RBP-class was detected as sharing sequence-level mosaicism with additional RBP-classes previously undetected by ECOD or pseudo-domain analysis, suggesting evolutionary connectivity at a recent timescale despite structural novelty. This is also the only structurally novel RBP-class found in podoviruses; in contrast to siphoviruses and myoviruses where multiple novel classes were identified, podoviruses were the most functionally well-characterised morphotype. The receptor target is unknown.

### Summary Table

| RBP-class | Source | n RBPs | n clusters | Morphotype(s) | Genera |
|---|---|---|---|---|---|
| Tailspike depolymerase | SSBH identification | 186 | 71 | All four | Multiple (17 genera) |
| Tailspike depolymerase with fiber-like elements | SSBH identification | 35 | 15 | All four | Multiple (7 genera) |
| RaK2 gp526-like | Published literature | 3 | 1 | Jumbo, Myo, Podo | *Alcyoneusvirus, Taipeivirus, Drulisvirus* |
| GDSL-like lipase containing tailspike | Foldseek+PDB100 | 3 | 2 | Myo, Podo | *Mydovirus, Przondovirus* |
| HRP29 tail-like variant 1 | Foldseek+PDB100 | 5 | 5 | Podovirus | *Drulisvirus, Koutsourovirus* |

| RBP-class | Source | n RBPs | n clusters | Morphotype(s) | Genera |
|---|---|---|---|---|---|
| HRP29 tail-like variant 2 | Foldseek+PDB100 | 1 | 1 | Podovirus | *Drulisvirus* |
| Putative esterase containing tailspike | Foldseek+PDB100 | 4 | 1 | Podovirus | *Przondovirus* |
| L-shaped tail fiber-like | Foldseek+PDB100 | 8 | 3 | Siphovirus | *Webervirus*, Unclassified |
| SU10 tail fiber-like | Foldseek+PDB100 | 1 | 1 | Siphovirus | *Webervirus* |
| GDSL-like lipase containing tail fiber | Foldseek+PDB100 | 1 | 1 | Myovirus | *Mydovirus* |
| Pam3 fiber-like | Foldseek+PDB100 | 2 | 1 | Myovirus | *Bimevirus, Mascletvirus* |
| Xylanase containing tail fiber | Foldseek+PDB100 | 2 | 1 | Myovirus | *Mydovirus* |
| T7 gp12 nozzle-like (TC2) | RBPseg-classify | 41 | 1 | Podovirus | *Przondovirus, Teetrevirus* |
| TC16 tail fiber (TC16) | RBPseg-classify | 1 | 1 | Podovirus | *Teetrevirus* |
| T4 gp34-like (TC0) | RBPseg-classify | 8 | 1 | Myovirus | *Slopekvirus, Jiaodavirus* |
| T4 gp36-like (TC11) | RBPseg-classify | 8 | 1 | Myovirus | *Slopekvirus, Jiaodavirus* |
| TC8 tail fiber (TC8) | RBPseg-classify | 7 | 1 | Myovirus | *Mydovirus* |
| *Vequintavirinae*-like short tail fiber | RBPseg-classify | 7 | 1 | Myovirus | *Mydovirus* |
| T4 gp12-like | Published literature | 5 | 1 | Myovirus | *Slopekvirus* |
| RaK2 gp534-like | Published literature | 3 | 3 | Jumbophage | *Alcyoneusvirus* |
| RaK2 gp98-like | Published literature | 3 | 2 | Jumbophage | *Alcyoneusvirus* |
| T4 gp37-like | RBPseg likely TC | 8 | 4 | Myovirus | *Slopekvirus, Jiaodavirus* |
| TC10-like tail fiber in *Mydovirus* — variant 1 | RBPseg likely TC | 1 | 1 | Myovirus | *Mydovirus* |
| TC10-like tail fiber in *Mydovirus* — variant 2 | RBPseg likely TC | 1 | 1 | Myovirus | *Mydovirus* |
| TC10-like tail fiber in *Mydovirus* — variant 3 | RBPseg likely TC | 1 | 1 | Myovirus | *Mydovirus* |
| TC12-like tail fiber in *Mydovirus* | RBPseg likely TC | 3 | 1 | Myovirus | *Mydovirus* |

| RBP-class | Source | n RBPs | n clusters | Morphotype(s) | Genera |
|---|---|---|---|---|---|
| TC12/TC17-like tail fiber in *Mydovirus* | RBPseg likely TC | 7 | 1 | Myovirus | *Mydovirus* |
| TC14-like long tail fiber in *Taipeivirus* | RBPseg likely TC | 3 | 1 | Myovirus | *Taipeivirus* |
| TC14-like short tail fiber in *Taipeivirus* | RBPseg likely TC | 3 | 2 | Myovirus | *Taipeivirus* |
| TC17-like tail fiber | RBPseg likely TC | 3 | 1 | Myovirus | *Jiaodavirus* |
| TC5-like tail fiber | RBPseg likely TC | 1 | 1 | Siphovirus | *Webervirus* |
| TC7-like tail fiber | RBPseg likely TC | 3 | 2 | Siphovirus | Unclassified |
| TC14-like tail fiber in *Sugarlandvirus* — variant 1 | RBPseg likely TC | 3 | 2 | Siphovirus | *Sugarlandvirus* |
| TC14-like tail fiber in *Sugarlandvirus* — variant 2 | RBPseg likely TC | 3 | 3 | Siphovirus | *Sugarlandvirus* |
| TC15-like tail fiber in siphovirus — variant1 | RBPseg likely TC | 1 | 1 | Siphovirus | *Henuseptimavirus* |
| TC15-like tail fiber in siphovirus — variant 2 | RBPseg likely TC | 2 | 1 | Siphovirus | Unclassified |
| TC10-like tail fiber in *Alcyoneusvirus* | RBPseg likely TC | 3 | 1 | Jumbophage | *Alcyoneusvirus* |
| TC16-like tail fiber | RBPseg likely TC | 1 | 1 | Jumbophage | *Eowynvirus* |
| TC12-like tail fiber in podovirus | RBPseg likely TC | 1 | 1 | Podovirus | Unclassified |
